# High Resolution Slide-seqV2 Spatial Transcriptomics Enables Discovery of Disease-Specific Cell Neighborhoods and Pathways

**DOI:** 10.1101/2021.10.10.463829

**Authors:** Jamie L. Marshall, Teia Noel, Qingbow S. Wang, Silvana Bazua-Valenti, Haiqi Chen, Evan Murray, Ayshwarya Subramanian, Katherine A. Vernon, Katie Liguori, Keith Keller, Robert R. Stickels, Breanna McBean, Rowan M. Heneghan, Astrid Weins, Evan Z. Macosko, Fei Chen, Anna Greka

## Abstract

High resolution spatial transcriptomics is a transformative technology that enables mapping of RNA expression directly from intact tissue sections; however, its utility for the elucidation of disease processes and therapeutically actionable pathways remain largely unexplored. Here we applied Slide-seqV2 to mouse and human kidneys, in healthy and in distinct disease paradigms. First, we established the feasibility of Slide-seqV2 in human kidney by analyzing tissue from 9 distinct donors, which revealed a cell neighborhood centered around a population of *LYVE1+* macrophages. Second, in a mouse model of diabetic kidney disease, we detected changes in the cellular organization of the spatially-restricted kidney filter and blood flow regulating apparatus. Third, in a mouse model of a toxic proteinopathy, we identified previously unknown, disease-specific cell neighborhoods centered around macrophages. In a spatially-restricted subpopulation of epithelial cells, we also found perturbations in 77 genes associated with the unfolded protein response (UPR). Our studies illustrate and experimentally validate the utility of Slide-seqV2 for the discovery of disease-specific cell neighborhoods.

**One-Sentence Summary:** High resolution Slide-seqV2 spatial transcriptomics in human and mouse kidneys.

## Main Text

Known for its structural complexity, the kidney performs vital functions that rely on spatially-distinct cellular compartments, yet an understanding of spatially resolved, cell type-specific responses to perturbations using high resolution technologies^1–3^ is lacking. The nephron, the functional unit of the kidney, is composed of epithelial, mesenchymal, and endothelial cells, as well as a network of immune cells that contribute to organ defense and repair from injury. Anatomically, each kidney has an outer layer, the cortex, containing the glomeruli, through which blood is filtered, and an inner layer, the medulla, where urine is concentrated. Urine flows within tubules that coalesce in the renal pelvis, draining into the ureter and on to the bladder^4^. Given its structural and functional complexity, second only to the brain, there is a significant need to develop spatial transcriptomics from intact kidney tissue both as a reference, and to uncover disease-specific processes.

Several studies using single cell (sc-) or single nucleus (sn-) RNA sequencing of human and mouse kidney in health and disease have been performed^5–21^, and have used spatial validation methods including immunofluorescence microscopy, fluorescence *in-situ* hybridization, and targeted panels of a few dozen RNA probes or antibodies. However, these spatial methods are hampered by relatively low throughput^22–28^. To date, efforts to spatially capture transcriptome-wide profiles in kidney cells *in situ* have been limited in resolution^29–31^. Given that several cell types in spatially restricted areas are the hypothesized drivers of injury as a result of ischemia and inflammation^32–36^, diabetic conditions^20,37–40^, and genetic disorders^41–47^, high resolution spatial transcriptomics in the kidney could catalyze mechanistic and therapeutic insights.

We recently generated a comprehensive cross-species scRNA-seq atlas^20^, that identified shared broad cell classes and unique cellular states between mouse and human kidney across three regions (cortex, medulla, and renal pelvis). Among many insights, this work revealed multiple macrophage subsets including *C1QB+LYVE1+* macrophages in human adult kidneys. A unique *TREM2+* subset was expanded with age in kidneys of diabetic and obese mice and humans^20^, mirroring obesity-associated macrophages in other tissues^48^. Based on these results, we hypothesized that these specialized macrophage populations might be localized in disease-related microenvironments, and that we could define these cell neighborhoods using high resolution spatial transcriptomics.

To understand the way in which cells act in concert in the kidney, we employed Slide-seqV2, a high resolution method for unbiased spatial transcriptomics, with a feature (or bead) size of 10μm most commonly capturing 1-2 cells/feature, and no more than 3 cells/feature^1,2^. Leveraging this near-single cell spatial resolution and single cell atlases^20^, we developed SlideSeqV2 working protocols and analysis pipelines for human and mouse kidney tissue. Furthermore, we compared tissue from healthy and diseased kidneys, because the side-by-side analysis allowed us to measure changes in the organization of cellular neighborhoods, thus taking full advantage of the high spatial resolution.

We first established the applicability of Slide-seqV2 in human kidney, in samples from 9 individual donors. We generated libraries from 4 arrays per sample with 2 arrays applied to cortex and 2 arrays applied to medulla in nephrectomy tissue (Table S1; samples were also used to build the single cell reference^20^). Histopathologic analysis performed by a kidney pathologist showed that seven samples were consistent with normal age-appropriate kidney tissue, one sample had subtle signs of early diabetic kidney disease (DKD), and one sample displayed clear evidence of injury (ischemia due to tumor compression; for relevant clinical data, see Table S1). The arrays were positioned at different locations in each tissue cross-section to cover as much area in each 3mm array as possible. Following library preparation, we mapped cell types using our recently developed single cell reference^20^ (Methods). Cell type classifications of Slide-seqV2 features were confirmed based on established markers of gene expression (Fig S1). We compared two different cell type assignment methods, NMFreg^1^ and the label transfer method from Seurat^49,50^. We found that Seurat was best able to map all cell types in human kidneys with expected spatial patterns of known cell types in the cortex and medulla, the two main anatomical layers of the kidney. For example, proximal convoluted tubules (PCTs) were enriched in cortex, whereas distal convoluted tubules (DCTs) and collecting ducts (CDs) were more densely packed in the medulla (Fig 1B; Fig S2-7; Fig S9A; Fig S10A). We thus used the Seurat label transfer method for all downstream analyses. As expected, glomeruli were found in the cortex and contained the appropriate cell types, including podocytes, endothelial cells (ECs), and mesangial cells (MCs) (Fig S2-7, S9A). We did not detect any significant changes in the early DKD sample (consistent with early disease lacking overt structural changes, as also previously shown^20^). We also found no significant changes in injured cortex (where the cell mappings showed numerous fibroblasts, in agreement with Periodic acid-Schiff (PAS) staining that showed extensive fibrosis) (Fig S8-9). Therefore, in human kidney, we focused our efforts on studying injured medulla in the sample that showed ischemic injury due to tumor compression.

**Fig. 1.**
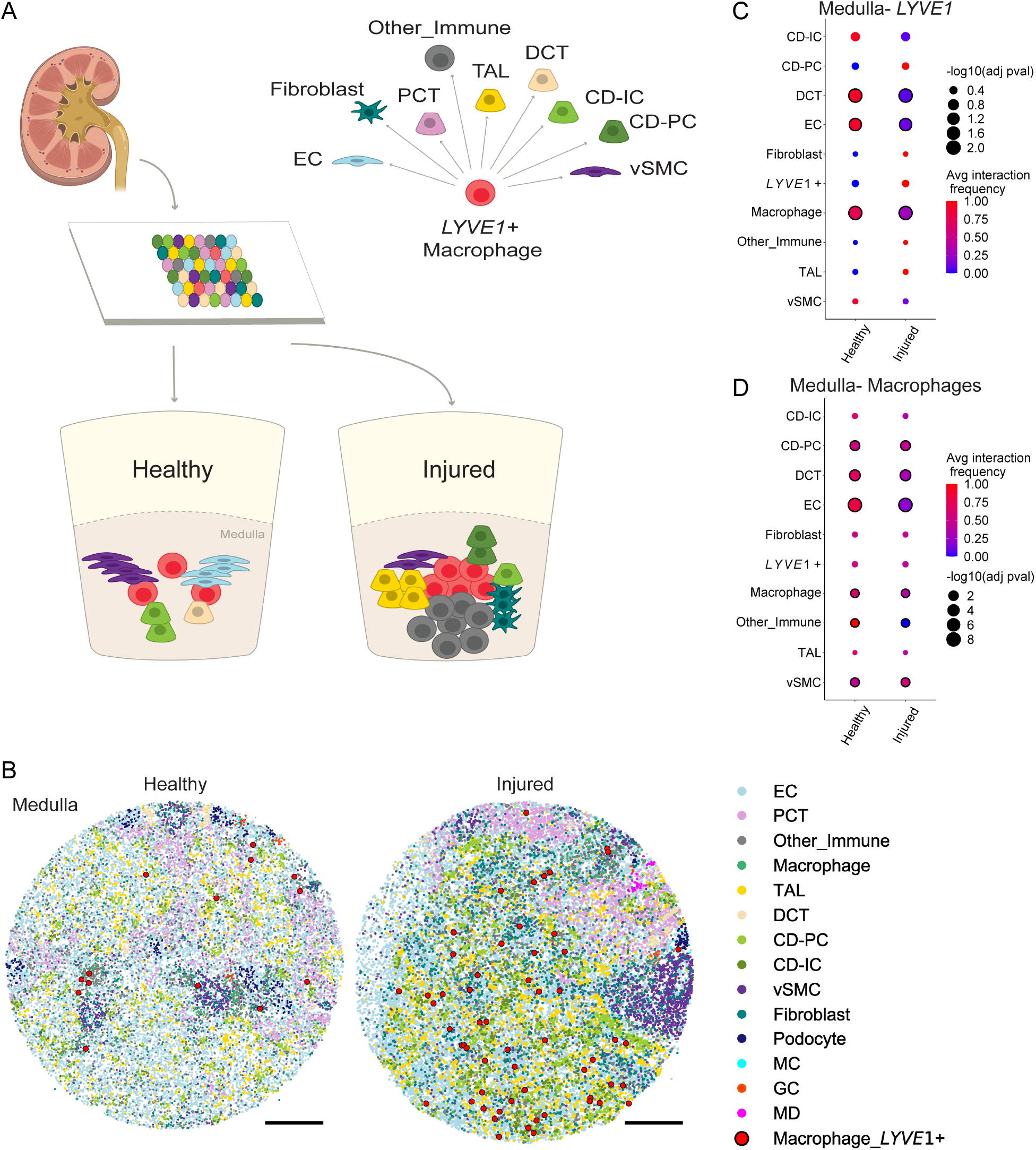
Slide-seqV2 spatial transcriptomics in human kidney informs methods to identify and quantify cell-cell interactions frequency and cell neighborhoods. (**A**) Schematic demonstrating the Slide-seqV2 method. A 10μm sagittal section of kidney is placed onto a Slide-seqV2 array. The arrays bind to RNA in the tissue and result in a spatial transcriptome with cDNA containing a barcode from each bead. (**B**) Medulla arrays showing all cell mappings and spatial locations of *LYVE1+* macrophages in large red circles. Images of individual cell populations are plotted in Fig S61-62. Scale bars, 500μm. (**C-D**) Uniquely enabled by the spatial resolution, we identified neighboring cell types in the medulla of healthy and injured human kidney. *LYVE1+* macrophages interact with endothelial cells (ECs) and distal convoluted tubular epithelial cells (DCTs) in healthy human medulla. These interactions are decreased in injured medulla. Dot plot shows the interaction frequency of (**C**) *LYVE1+* macrophages and (**D**) all macrophages versus every other cell type in the tissue array. The color of the dots indicates the intensity of the interaction (red, increased; blue, decreased interaction). The size of the dots indicates the adjusted p-value for cell-cell interactions, and significance is shown by a black outline around the dot (adjusted p-value < 0.05). Cell types not represented in the dot plot had no interactions with macrophages of interest.

Given that ischemic injury is associated with inflammatory changes that involve macrophage populations^32,36^, we hypothesized that focusing on macrophages^20^ might help us identify injury-specific changes. First, we identified macrophage beads in medulla from age matched healthy and injured human tissue arrays (Fig S10A-B, D, F; validated by quantification of *C1QB+* cells, a canonical macrophage cell marker^51^, in hybridization chain reaction (HCR) of the entire kidney tissue section). Among these *C1QB+* macrophages, we prioritized looking for changes in macrophage subtypes that have been previously implicated in disease processes in the kidney and other tissues^20,48,52–58^. We thus detected an increase in *LYVE1+* macrophages (large red beads) in medullary injured tissue (Fig 1B, Fig S10C, S10E-F; validated by HCR showing *C1QB+ LYVE1+* cells). Taking advantage of the high spatial resolution, we calculated the average interaction frequency in the cellular neighborhood between *LYVE1+* macrophages and immediately adjacent cell types (Fig 1A-D). In human healthy medulla, *LYVE1+* macrophage expansion led to a spatial neighborhood composed of DCT epithelial cells and EC (Fig 1C). In order to determine if these interactions were specific to *LYVE1+* macrophages, the same analysis was performed with all medullary macrophages that, in addition to DCT and EC, also interacted significantly with principal cells (CD-PC), other immune cells, and vascular smooth muscle cells (vSMC) (Fig 1D). Overall, *LYVE1+* macrophage interactions with DCT and EC and all macrophage interactions with other immune cells were reduced in human injured medulla, perhaps consistent with increased fibrosis (Fig 1C-D; Table 1; Fig S8A, C, E). Most importantly, this work established the applicability of Slide-seqV2 in human kidney, including new computational approaches (Methods) that provide the blueprint for future, more detailed studies in human kidney tissue.

Since mouse tissue of the quality required for successful implementation of spatial transcriptomics is more readily attainable than human tissue, we turned our attention to established mouse models of kidney disease. We thus generated libraries from BTBR *ob/ob* mice, a genetic model of DKD, where the first signs of injury localize to the glomeruli^38–40^. We chose this well-characterized mouse model with an available single cell reference^20^ because we wanted to take advantage of the high spatial resolution of Slide-seqV2 to detect transcriptome-wide changes in or near glomeruli, spatially defined structures composed of rare cell types^59^. We were particularly interested in the Juxtaglomerular apparatus (JGA), a structurally distinct, spatially compact structure located adjacent to glomeruli, that is comprised of renin (*Ren1*)-producing granular cells (GCs), and rare macula densa (MD) cells, which, working in concert, regulate blood flow to the kidney filter^60–62^. In DKD, the JGA is known to shift its cellular composition^63,64^, but, to date, the underlying cellular changes have not been described with high spatial, near single cell resolution.

Seven arrays were captured from 10μm sagittal sections of kidney from four BTBR *wt/wt* controls and four BTBR *ob/ob* mice. We used our recently developed scRNA-seq reference^20^ for cell type mappings (Methods; Fig S11-13), and characterized the transcriptional effects of disease in three ways. First, we captured a significant increase in the measured area of individual glomeruli in BTBR *ob/ob* mice compared to controls (Fig 2A-E; Fig S14A-D) ^38,40^. Second, we examined the composition of the JGA. We identified glomeruli-adjacent beads positive for *Ren1*^65^ and negative for *Slc12a3* (a classical distal nephron marker^6,66^), and another set positive for *Ptger3, Klf6*, or *Nos1* (MD marker genes^6,67^), and labeled the corresponding cells as GC or MD, respectively. MDs were rare (~0.6% of unfiltered TAL calls were classified as MD). However, in diabetic BTBR *ob/ob* tissue, we detected a significant increase in the percentage of GCs (Fig 2F; Fig S14E) and in the distance between the center of each glomerulus and GCs (Fig 2G-H; Fig S15A). Consistent with overall tissue hypertrophy, as observed in BTBR *ob/ob* mice^38–40^, we found a significant increase in the distance between the centers of glomerular and GC structures. In contrast, the distance measured between the edges of the glomeruli and GC structures did not show a significant increase (Fig S14G), arguing against GCs migrating away from glomeruli. This spatial expansion of the JGA in diabetes, reminiscent of previous work in glomeruli using labor-intensive lineage tracing studies^63,68–70^, was readily detected by Slide-seqV2.

**Fig. 2.**
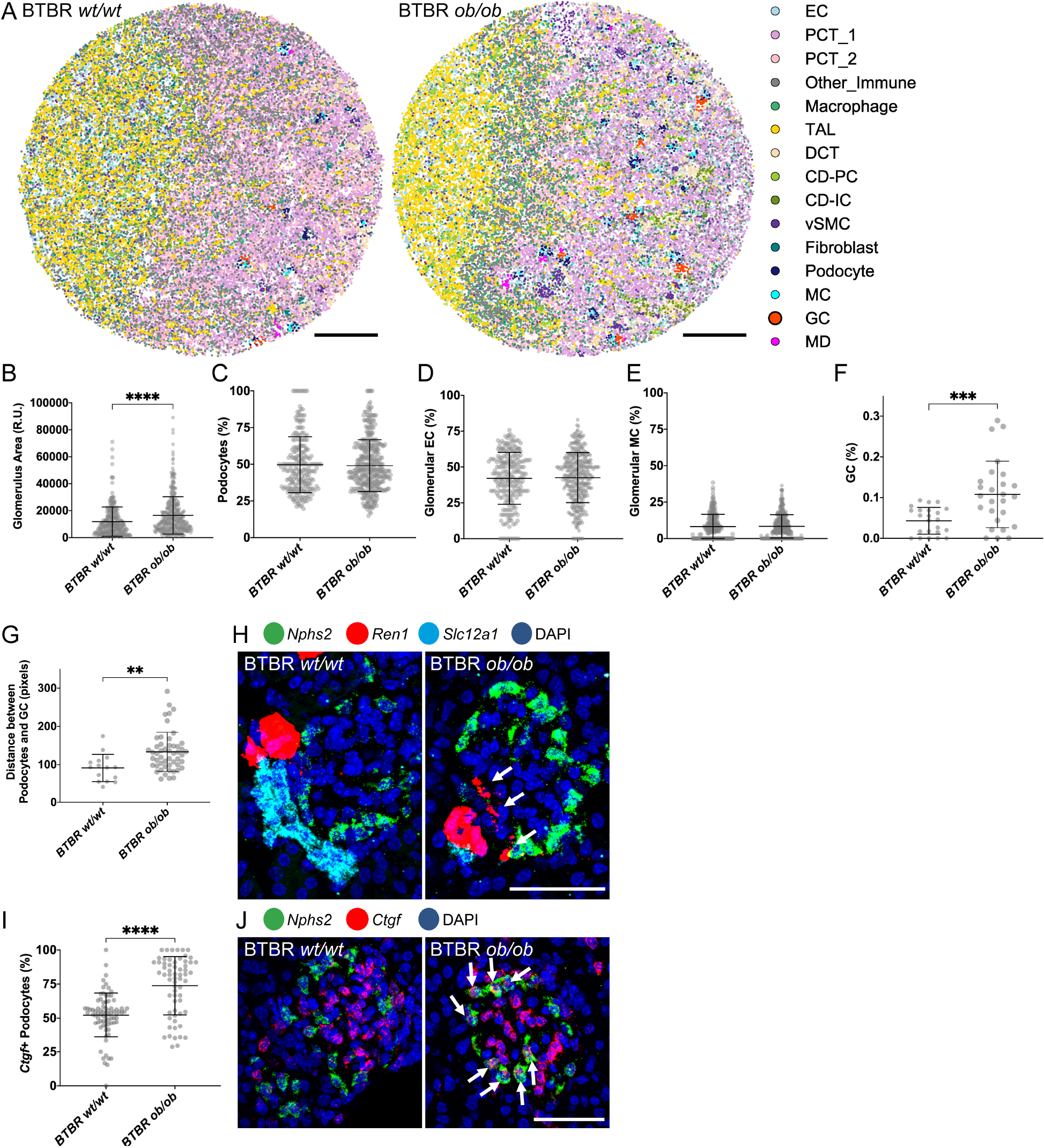
Near single cell spatial resolution in diabetic mouse kidney reveals an expansion of granular cells, and a disrupted blood flow regulating apparatus. (**A**) Arrays displaying cell types in BTBR *wt/wt* (left) and BTBR *ob/ob* diabetic mice (right). Images of individual cell populations are plotted in Fig S64. Scale bars, 500μm. (**B-F)** Plots from Slide-seq arrays showing (**B**) average area of glomeruli (p-value <0.0001), (**C**) percentage of beads classified as podocytes, (**D**) percentage of beads classified as glomerular endothelial cells, (**E**) percentage of beads classified as glomerular mesangial cells, (**F**) percentage of beads classified as granular cells (p-value 0.0009) in BTBR *wt/wt* and BTBR *ob/ob* mice. (**G**) Plot from Slide-seq arrays showing the average distance between the center of glomerulus and granular cell cluster in BTBR *wt/wt* and BTBR *ob/ob* mice (p-value 0.0018). Data was obtained from cross sections of 4 mice per genotype. (**H**) HCR validation images, showing *Nphs2+* podocytes, *Ren1+* granular cells, *Slc12a1+* TAL, and all cells in DAPI. Scale bar 50μm. Disorganized *Ren1+* cells are denoted with white arrows. (**I**) HCR quantification of (p-value <0.0001) and (**J**) HCR images of *Ctgf+ Nphs2+* injured podocytes (arrows). Scale bar 50μm. Data obtained from entire cross sections of a kidney from 4 mice per genotype.

Taking advantage of the fact that each cell in the spatial array is linked to a transcriptome comprising of several hundred genes (Methods), we developed a clustering and differential gene expression (DGE) pipeline. Focusing on podocytes, whose injury is a hallmark of DKD^38^, we compared the transcriptional profiles for all podocyte beads. Specifically, we examined their clustering in an unbiased fashion, irrespective of whether the podocyte beads came from a BTBR *wt/wt* or diabetic BTBR *ob/ob* array; we let their transcriptional profiles dictate whether they were considered “healthy” or “diseased” (Fig S16; Tables S2-5). Based on transcriptomes alone, podocyte beads from the “diseased” cluster that mapped to diabetic BTBR *ob/ob* glomeruli were found to express *Ctgf*, a known podocyte injury marker gene (Table 3). Furthermore, these results were validated by HCR measuring co-expression of *Ctgf* and *Nphs2*, a podocyte marker gene^71,72^ (Fig 2I-J; Fig S15B). Pathway analysis highlighted mechanisms of high relevance to podocyte pathobiology such as lipid-mediated signaling, regulation of stress fiber assembly, and apoptosis (Table S6, Fig S14F)^73–76^. In sum, we captured disease-specific changes in spatially-restricted neighborhoods of podocytes and granular cells in a mouse model of DKD. More generally, implementing Slide-seqV2 in mouse tissue provided a valuable foundation for our subsequent studies of a significantly understudied toxic proteinopathy.

Seeking to define actionable disease mechanisms, we turned to homozygous UMOD-C125R knockin (KI) mice^41^. These mice are a model of a poorly understood, monogenic disorder called UMOD Kidney Disease (UKD)^42^, caused by the toxic accumulation of mutant UMOD protein in the kidney^41^. We generated libraries in five arrays from kidneys of three WT control and five homozygous UMOD-KI six-month-old mice^41^. At this age, homozygous UMOD-KI mice have detectable kidney disease associated with inflammation and fibrosis^41^. Cell types were mapped to arrays^20^, and a border was drawn to separate cortex from medulla based on the presence of PCTs in the cortex and the concentration of TAL and CDs in the medulla (Fig S17-19). A significant increase in fibroblast and macrophage beads was observed in the medulla of homozygous UMOD-KI mice compared to controls (validated by HCR for *Ctgf+* and *Itga8+* fibroblasts^77^ and *C1qb+* macrophages^51^; Fig 3A-C; Fig S20). Significant *Ctgf* expression was detected by HCR in medullary *Umod+* TAL epithelial cells^78^ of homozygous UMOD-KI mice, suggesting cellular injury in *Umod+* TALs (Fig 3E; Fig S20F). Unbiased clustering and DGE analysis of cortex and medulla identified disease-specific clusters for TAL, fibroblasts, macrophages, vSMCs, and CD-PCs in the medulla, as well as fibroblast, glomerular EC, and CD-PC in the cortex (Tables S7-14; Fig S21-22).

**Fig. 3.**
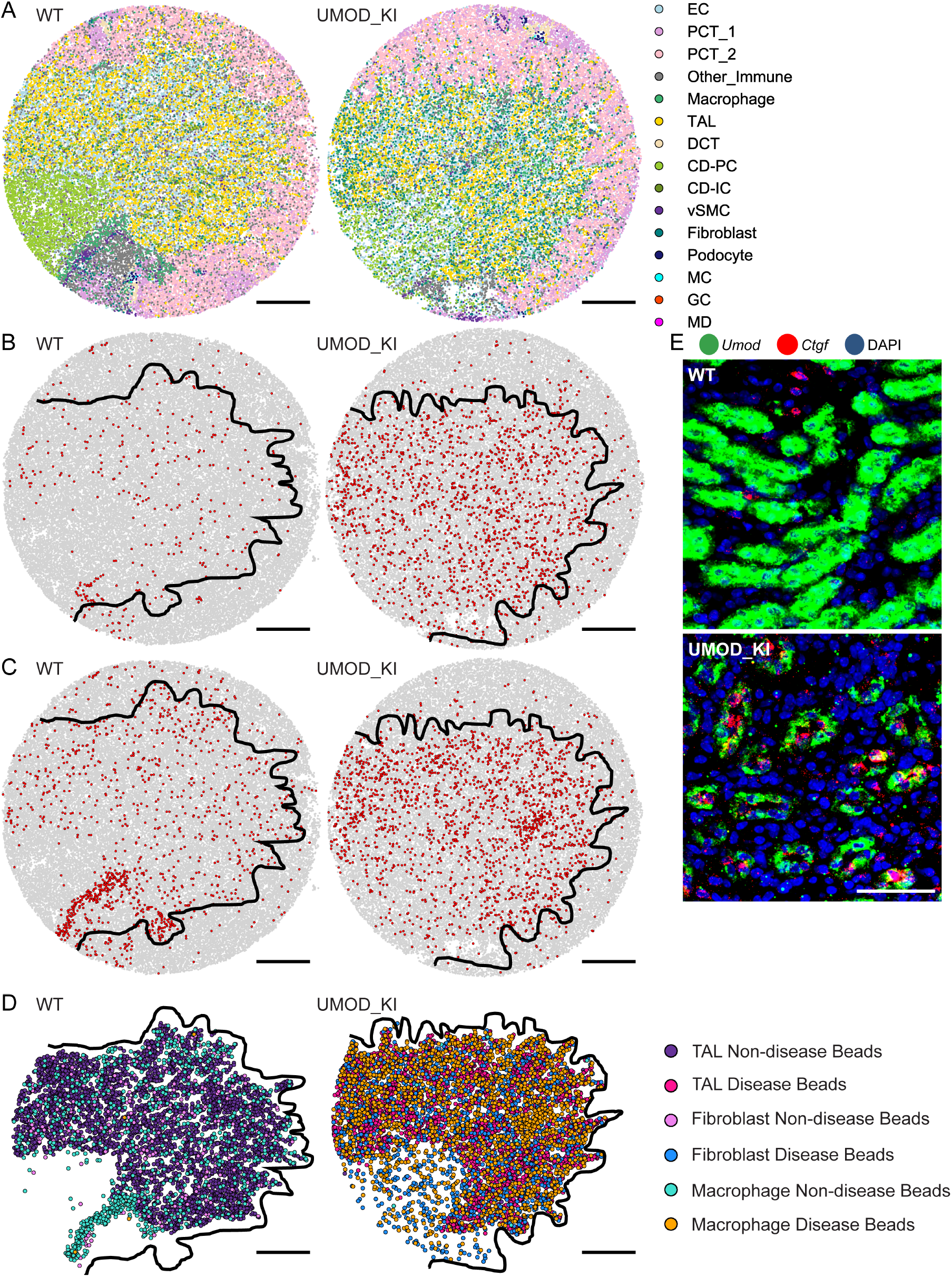
Slide-seqV2 reveals disease-specific, medulla-restricted cell neighborhoods and injured epithelial cells in a mouse model of a toxic proteinopathy due to *Umod* mutations. (**A**) Arrays displaying all cell types in WT (left) and UMOD-KI mice (right). Images of individual cell populations are plotted in Fig S65. Arrays showing delineation of cortex vs medulla in WT and UMOD-KI arrays where red beads show the spatial location and quantity of (**B**) fibroblasts and (**C**) macrophages. (**D**) Based on unbiased DGE, array plots show spatial mapping of TAL (non-disease in purple; disease in red), fibroblast (non-disease in pink; disease in blue), and macrophage beads (non-disease in green; disease in orange) in WT and UMOD-KI tissue. Beads classified by DGE as “disease” map primarily onto the medulla of UMOD-KI arrays, compared to beads classified as “non-disease” that map primarily onto the medulla of WT arrays (fisher exact test p<10^-100). Scale bars, 500μm. (**E**) HCR images of *Umod+ Ctgf+* double positive injured TALs. Scale bar, 50μm.

Taking advantage of the spatial resolution, we wondered whether we could determine the location of medullary disease-associated cell types compared to healthy cells based only on unbiased clustering, without any prior knowledge of their localization in the tissue. We thus classified individual beads into either healthy or disease states based on gene expression profile alone, and characterized their spatial distribution across all of the arrays from WT and UMOD-KI mice. This analysis showed that healthy (“non-disease”) cells largely mapped onto the medulla of WT arrays (99.8%), while disease-associated cell types were predominantly localized in the medulla of homozygous UMOD-KI arrays (Fig 3D; 63.1%, fisher exact test p<10^-100). This analysis, uniquely afforded by high resolution spatial transcriptomics, suggested that fibroblasts (blue), TAL epithelial cells (red), and macrophages (orange) in the medulla of UMOD-KI mice may form disease-specific cell neighborhoods.

To probe the question of cell neighborhoods further, we employed the spatial neighbor interaction frequency method, as we previously established in human tissue (Fig 1C-D), to determine the identity of cell types immediately adjacent to macrophages (Fig 4A-C; Methods). We thus identified medulla-specific cell neighborhoods centered on macrophages interacting with diverse kidney cell types (Fig 4A-C). Of interest, specifically in the setting of disease, we observed enhanced interactions between medullary macrophages and CD-PC, collecting duct intercalated cells (CD-IC), and ECs (Fig 4C). Focusing further on macrophage subsets, we found a significantly higher percentage of *Trem2+* macrophages in the medulla of UMOD-KI mice (large red beads; Fig 4D-E). HCR confirmed a significant expansion of double-positive *C1qb+Trem2+* cells in the medulla of UMOD-KI mice (Fig 4F-G). These results suggest that *Trem2+* macrophages in the medulla of UMOD-KI mice may expand in response to interactions with specific cell neighbors, a previously unknown mechanism in this disease.

**Fig. 4.**
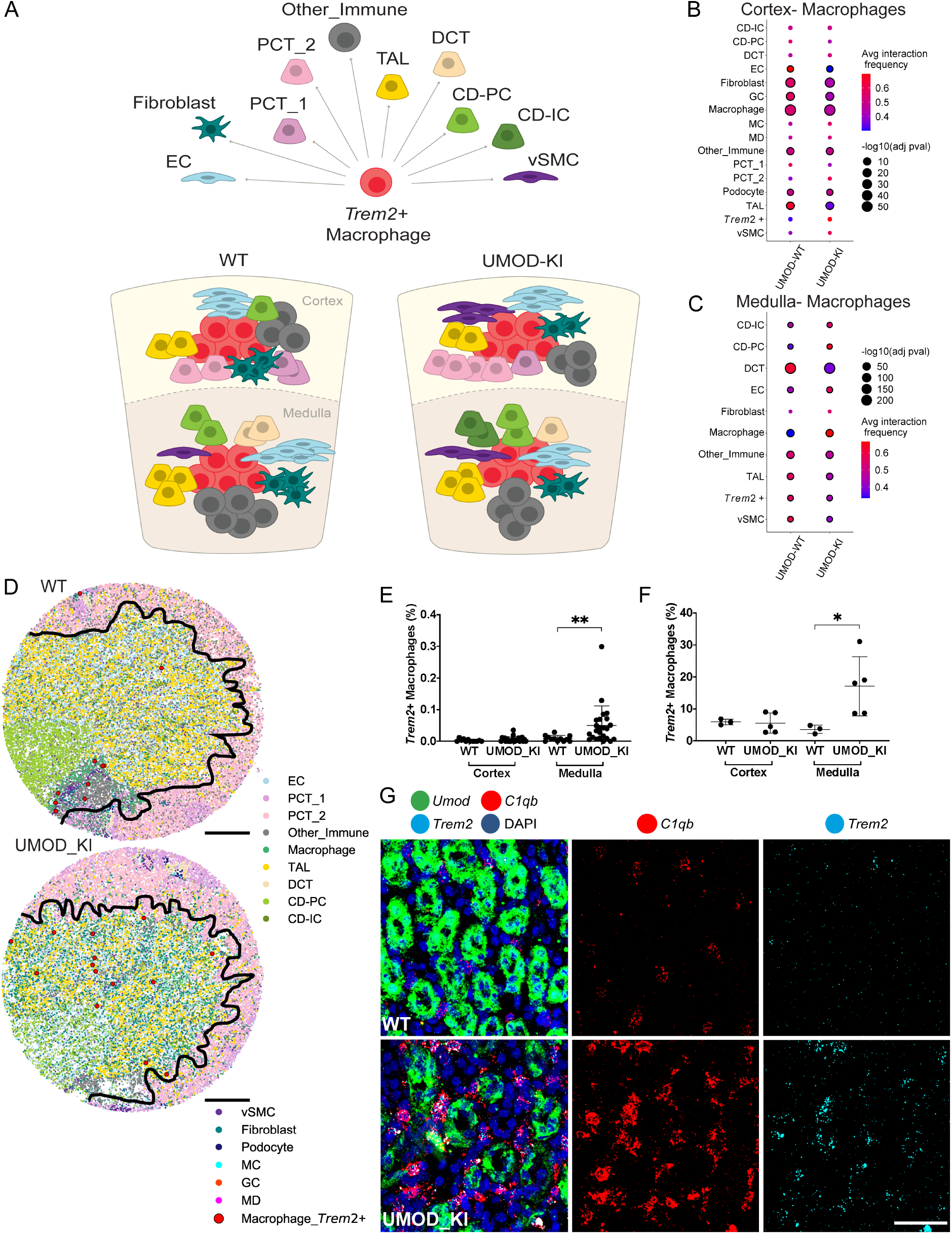
Rare *Trem2+* macrophage expansion associated with disease and localized specifically to kidney medulla. (**A**) Schematic showing location, cortex vs medulla, of *Trem2+* macrophages and immediately adjacent neighboring cell types. (**B-C**) Uniquely enabled by high spatial resolution, we quantified macrophage-neighbor cell interactions. Dot plots show average cell-cell interaction frequencies (relative proportions) for all macrophages in (**B**) cortex and (**C**) medulla. Significant interactions are displayed with a black border around colored circles. Cell types not represented in the dot plot had no interactions with macrophages of interest. (**D**) Arrays showing all cell mappings, cortex and medulla delineation, and spatial localization of *Trem2+* macrophages in large red circles. Scale bars, 500μm. (**E**) Plots from Slide-seqV2 arrays showing percentage of beads classified as *Trem2+* macrophages (p-value 0.003). Each point represents an array from 3 WT and 5 UMOD-KI mice. Error bars represent standard deviation of the mean. (**F**) Plots generated from HCR validation showing percentage of cells that are *Trem2+* macrophages (p-value 0.0357). Each point represents a mouse. Error bars represent standard deviation of the mean. (**G**) HCR images from 2 different WT and UMOD-KI mice showing *Trem2+ C1qb+* macrophages in the medulla (*Umod*). Scale bar 50μm.

To gain a deeper understanding of disease-relevant pathways, we probed our data for detectable changes in the UPR, a cellular mechanism triggered by the accumulation of misfolded mutant UMOD protein in UMOD-KI mouse kidneys^41,42^. Using a validated UPR pathway signature^79^, DGE analysis between WT and UMOD-KI arrays (Fig 5A, Fig S23) revealed changes in the UPR^80,81^ specifically in medullary TALs (Fig 5A; 77 genes). In contrast, when we aggregated all TALs (cortex and medulla), we did not detect any significant DGE changes between WT versus KI tissue (Fig S23A). Similar analyses focusing on cortical TALs alone, or PCTs alone did not yield any significant differences in UPR gene expression either (Fig S23B-C). The loss of signal when looking at the totality of TALs (or other cell subsets) suggests that if we only relied on bulk RNA-Seq or even on scRNA-Seq without the high spatial resolution afforded by Slide-SeqV2, we would not have been able to identify this disease-associated UPR signature.

**Fig. 5.**
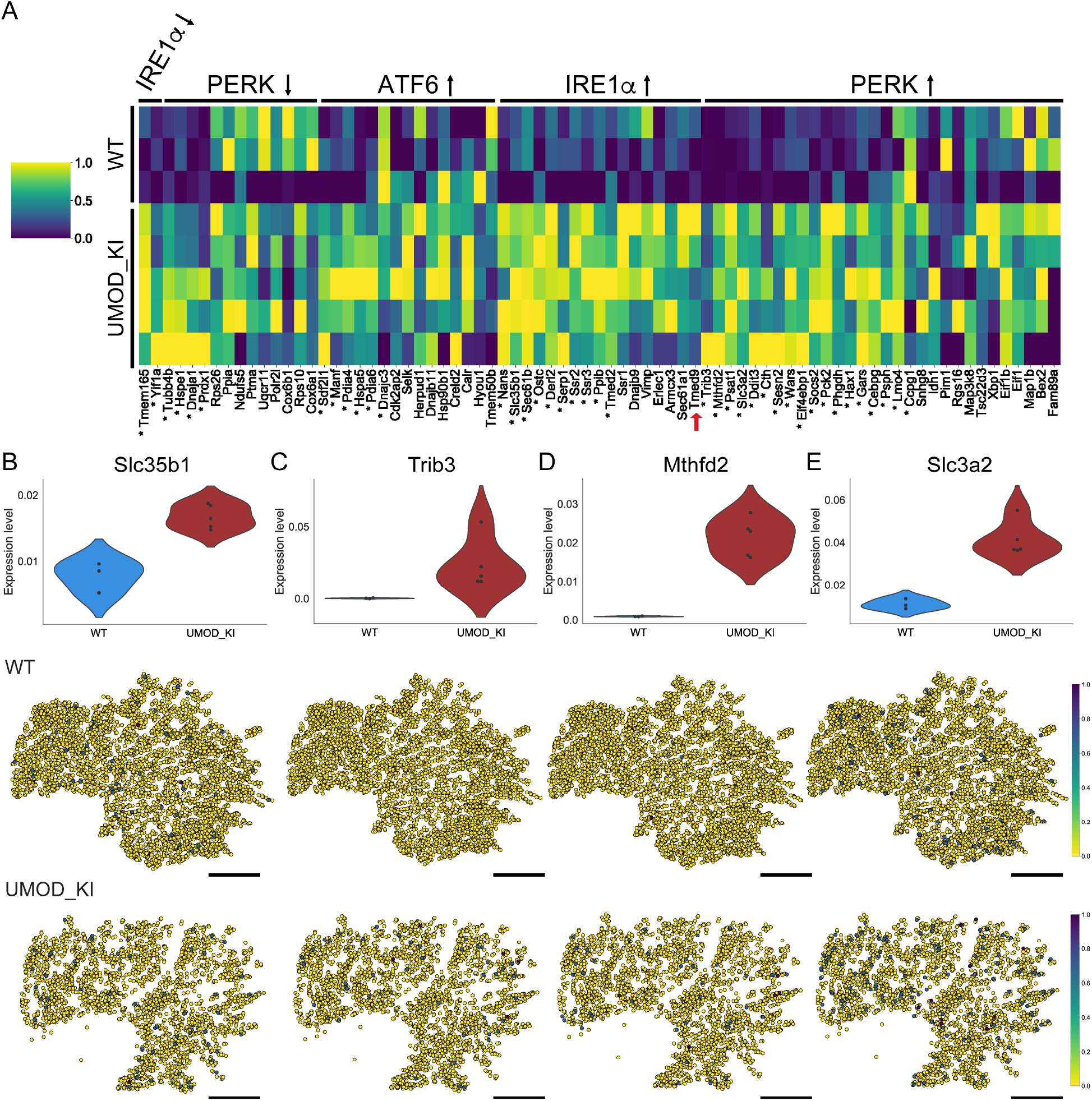
Spatially-restricted, cell-specific and disease-associated perturbations in 77 genes associated with the UPR in medullary TAL epithelial cells uniquely revealed by Slide-seqV2. (**A**) Heatmap showing relative expression level of 77 UPR genes averaged across arrays in WT and UMOD-KI mice from medullary TAL beads. *Tmed9* is indicated with a red arrow. Significant genes are denoted with an asterisk (Benjamani-Hochberg-corrected p<0.05). Violin plots showing expression level in medulla TAL (top) and all medulla TAL beads in arrays (middle, bottom) are shown with expression level for notable genes in (**B**) IRE1α upregulated pathway gene, *Slc35b1* and PERK upregulated pathway genes, (**C**) *Trib3*, (**D**) *Mthfd2*, and (**E**) *Slc3a2*. Scale bars, 500μm.

Three to four markers of the UPR have been previously used to implicate this pathway in UKD^41,42^, however, coordinate changes in the expression of 77 UPR genes in *Umod*-expressing medullary TALs have not been described before. Among significant genes in medullary TAL features were *Slc35b1* (IRE1a pathway) and *Trib3, Mthfd2*, and *Slc3a2* (PERK pathway; Fig 5B-E; Fig S24-29), which showed that the medullary TAL epithelial cells were selectively and most highly affected by two specific UPR pathways, IRE1α and PERK (Fig S27-29). Among UPR-associated genes, *Tmed9* was upregulated and selectively localized to *Umod*-expressing medullary TALs (Fig 5A, red arrow). We have previously shown that the small molecule (BRD4780) targets TMED9 for the treatment of MUC1 Kidney Disease, a toxic proteinopathy caused by the accumulation of mutant MUC1 protein in epithelial cells in the kidney^43^. We reasoned that the specific co-localization of *Umod* and *Tmed9* in cells with detectable UPR-mediated injury (medullary TALs) pointed to the utility of Slide-seqV2 for the detection of therapeutically relevant targets.

## Discussion

Spatial transcriptomics has revolutionized the field of single cell genomics and provided new insights into many biological systems such as the brain, heart, liver, and testes^82–88^. However, the potential of spatial transcriptomics to identify therapeutically actionable pathways has not yet been explored. In this study, we explored the utility of spatial transcriptomics to uncover disease-associated cell-cell interactions and pathways, and, importantly, we performed rigorous experiments (HCR, mouse studies) to experimentally validate these findings. Several critical conclusions can be drawn from these studies.

First, we developed tools and methods to identify disease-specific cell neighborhoods. In human kidney, we found a cell neighborhood centered around *LYVE1+* macrophages, most notably in the medulla. In other tissues, *LYVE1+* macrophages have been shown to protect from fibrosis^53^. We can therefore speculate that *LYVE1+* cell neighborhoods in human kidneys may reflect an adaptive response mounted to prevent or abrogate the pro-fibrotic sequelae of injury. Disease-specific cell neighborhoods were also identified in a mouse with diabetic kidney disease. In line with prior studies^63,68–70^, we characterized a disordered kidney filter and blood flow regulating apparatus (JGA) associated with an expanding population of rare GCs in diabetic kidneys. These findings illustrated how the high spatial resolution of Slide-seqV2 empowered us to detect changes within and between spatially restricted structures.

Second, drawing from the blueprint established in these initial studies in human and mouse, we made a new discovery in mice with a toxic proteinopathy (UKD). Specifically, we identified a disease-specific, medulla-restricted cell neighborhood composed of *Trem2+* macrophages^20^, and specific epithelial (TAL) and fibroblast cell populations. Since resident macrophage *Trem2* acts as an immunomodulatory receptor on resident macrophages that senses tissue damage and negatively regulates inflammation^48,54–58^, our results suggest that medullary *Trem2+* macrophages may expand in response to neighboring cell injury. More generally, resident macrophage-epithelial cell neighbor interactions may underlie several kidney diseases^20^ associated with inflammation and fibrosis^89^.

Lastly, a critical aspect of this study was the development of new analytical methods and pipelines to map cell states and interactions onto spatial arrays from mouse and human kidney, relying exclusively on spatial near single cell transcriptome-wide profiles. Taken together, our experiments illustrate how we can harness the power of Slide-seqV2, and spatial transcriptomics more broadly, to gain biological insights by a “one and done” approach that simultaneously monitors cell-cell interactions and numerous genes and pathways in an unbiased fashion. Similar to RNA-Seq supplanting the need for RT-PCR, we speculate that this technology may mature to the point that it may minimize or even replace “one-by-one” molecule detection by labor intensive *in situ* and immunolocalization methods.

In sum, our work provides a foundational framework for near single cell spatial transcriptomics in the kidney, a functionally complex tissue with intricate architecture. Building on the paradigm that many diseases, from cancer to neurodegeneration, depend on dynamic cell-cell interactions in spatially restricted microenvironments^90,91^, our studies illustrate the utility of high resolution spatial transcriptomics, to uncover disease-specific cell neighborhoods and pathways.

## Materials and Methods

All code can be found on github: https://github.com/marshalljamie/Kidney-Slide-seq.

### Mouse Models/Tissue

BTBR *wt/wt* and BTBR *ob/ob* mice were purchased from JAX labs (strain 004824)^38^. Male mice were aged to 13 weeks in the Broad Institute vivarium with alpha dry bedding and acidified water. UMOD-KI mice were transferred to the Broad Institute from the Thakker lab at the University of Oxford^41^. UMOD-KI heterozygous mice were mated to produce 6 month old WT and UMOD-KI homozygous male progeny used in these studies. All animals were subjected to intracardiac perfusion of PBS to remove blood. Tissues were dissected, mounted in OCT, and flash frozen in liquid nitrogen cooled isopentane for 1 min. Samples were then placed on dry ice until long-term storage in the −80°C. All procedures performed are IACUC approved on Broad Institute animal protocol # 0061-07-15-1.

### Human Tissue

All human tissues were obtained from healthy regions of tumor nephrectomy samples on Partners IRB protocol # 2011P002692 at Mass General Brigham (AW). Cortex and medulla samples were collected from each patient and samples were flash frozen in liquid nitrogen prior to storage in the −80°C.

### Slide-seqV2

Bead synthesis.

Bead barcodes were synthesized either by the ChemGenes Corporation or in house on an Akta Oligopilot 10 on one of two polystyrene supports, Agilent PLRP-S-1000A 10μm particles or 10μm custom polystyrene from AMBiotech. Oligonucleotide synthesis was performed as described below. Beads were used with one of the two following sequences: ChemGenes Corporation beads (5′-TTTTTTTTCTACACGACGCTCTTCCGATCTJJJJJ JJJTCTTCAGCGTTCCCGAGAJJJJJJJNNNNNNNNT30-3′) and custom synthesis beads (5′-TTT_PC_GCCGGTAATACGACTCACTATAGGGCTACACGACGCTCTTCCGATCTJJJJJJJ JTCTTCAGCGTTCCCGAGAJJJJJJJTCNNNNNNNNT25-3′ (pawpuck3); PC, a photocleavable linker; J, bases generated by split-pool barcoding, such that every oligonucleotide on a given bead has the same J bases; N, bases generated by standard base mixing of a 1:1:1:1 ratio of A, C, T and G, such that every oligonucleotide on a given bead has different N bases; TX, a sequence of X thymidines; V, an A, C or G but not T. Bead synthesis. PLRP-S resin (~10-μm mean particle diameter; Agilent) was functionalized with a non-cleavable linker by ChemGenes. The functionalized beads were then used as a solid support for reverse-direction phosphoramidite synthesis (5′ to 3′) on an Akta OligoPilot 10 using a standard solid-phase DNA synthesis protocol. 5′-CE (b-cyanoethyl) phosphoramidites were purchased from Glen Research and were dissolved in anhydrous acetonitrile to obtain a concentration of 0.1M. Successive phosphoramidites were coupled for 5min using 5-benzylmercaptotetrazole (0.30M in acetonitrile) as an activator. Oxidation of the phosphite backbone to a phosphate backbone was achieved using iodine. Failure sequences were capped using acetic anhydride. Dichloroacetic acid was used as a detritylation reagent. For split-pool synthesis cycles, beads were suspended in acetonitrile and divided into four equal portions. These bead aliquots were then placed in four separate synthesis columns and reacted with dG, dC, dT or dA phosphoramidites. After each cycle, beads were pooled, suspended in acetonitrile and aliquoted into four equal portions. The split-pool procedure was repeated 15 times in total (two blocks of eight and seven cycles) to obtain 415 (~109) unique barcode sequences. After synthesis completion, the protecting groups from the nucleobases and phosphate backbone were removed by treating beads with 30% ammonium hydroxide containing 10% diethylamine for 40h at room temperature. The beads were centrifuged, and the supernatant was discarded. Beads were then washed three times with 1% acetone in acetonitrile, three times with water and three times with a buffer consisting of 10mM Tris and 1mM EDTA pH 8.

### Puck preparation

Puck preparation was performed as described previously^2^. Briefly, beads were pelleted and resuspended in water with 10% DMSO at a concentration between 20,000 and 50,000 beads per μl. Then, 10μl of the resulting solution was pipetted into each position on the gasket. The coverslip gasket filled with beads was centrifuged at 850g for at least 30min at 40°C until the surface was dry.

### Puck sequencing

Puck sequencing was performed in a Bioptechs FCS2 flow cell using an RP-1 peristaltic pump (Rainin) and a modular valve positioner (Hamilton MVP). Flow rates between 1ml min–1 and 3ml min–1 were used during sequencing. Imaging was performed using a Nikon Eclipse Ti microscope with a Yokogawa CSU-W1 confocal scanner unit and an Andor Zyla 4.2 Plus camera. Images were acquired using a Nikon Plan Apo × 10, 0.45-NA objective. After each ligation, images were acquired in the following channels: 488-nm excitation with a 525/36-nm emission filter (MVI, 77074803), 561-nm excitation with a 582/15-nm emission filter (MVI, FF01-582/15-25), 561-nm excitation with a 624/40-nm emission filter (MVI, FF01-624/40-25) and 647-nm excitation with a 705/720-nm emission filter (MVI, 77074329). The final stitched images varied in size depending on the size of the Slide-seq array. For the arrays presented in this work the final stitched images were 6,030 pixels by 6,030 pixels. Pucks were sequenced using a sequencing-by-ligation approach with a SOLiD dibase-encoding and with a monobaseencoding strategy previously described^1,2^.

This protocol was used for all slide-seq version 2 arrays http://dx.doi.org/10.17504/protocols.io.bpgzmjx6^2^. Briefly, 10μm sections (Leica, CM1950) were overlaid and melted onto spatial arrays. Seven arrays were collected from four BTBR *wt/wt* and four BTBR *ob/ob* mice, one kidney per mouse. Seven arrays were necessary to cover the entire cortex and collect enough glomeruli for analysis. Five arrays were collected for three WT and five UMOD-KI mice, one kidney per mouse. Five arrays ensured coverage of the entire medulla. Two arrays were collected from each human tissue sample, which resulted in two cortex and two medulla arrays per individual.

All reagents were diluted in ultrapure water (Life Technologies, Inc., 10977023). Arrays covered in tissue were then transferred to tubes with 6xSSC (Life Technologies, Inc., 15557044) containing RNAase inhibitor (Takara Bio, 2313B) and incubated for 15min. Arrays were then dipped in 1xRT buffer (Life Technologies, Inc., EP0753) and transferred to RT tubes (Maxima RT: Life Technologies, Inc., EP0753; 10mM dNTP Life Technologies, Inc., 4303443; RNAase inhibitor: Takara Bio, 2313B; 50μM Template Switch Oligo: IDT, AAGCAGTGGTATCAACGCAGAGTGAATrG+GrG) for a 30min room temperature incubation. RT tubes were then transferred to 52°C for a 90min incubation. Proteinase K and tissue clearing solution (Tris-HCl, pH 7.5: Life Technologies, Inc., 15567027; NaCl: American Bioanalytical, AB01915-01000; SDS (w/v): Life Technologies, Inc., 15553027; EDTA: Life Technologies, Inc., 15575020; Proteinase K: New England BioLabs, Inc., P8107S) is then added to the RT tube and the array is incubated at 37°C for 30min. Beads are removed from glass and resuspended in TE-TW (TE buffer: Sigma-Aldrich, Inc., 8910-1L; Tween-20: VWR International, LLC, 100216-360) and subjected to 2 TE-TW washes followed by centrifugation for 2min at 3000rcf. Beads are then resuspended in Exonuclease I mix (New England Biolabs, Inc., M0293L) and incubate at 37°C for 50min. This is followed by 2 washes in TE-TW, 5min room temperature incubation in 0.1N NaOH (Sigma-Aldrich, Inc., SX0607N-6), another TE-TW wash, and incubation in the second strand synthesis buffer (Maxima RT: Life Technologies, Inc., EP0753; 10mM dNTP Life Technologies, Inc., 4303443; dnSMRT oligo: IDT, AAGCAGTGGTATCAACGCAGAGTGANNNGGNNNB; Klenow: New England BioLabs, Inc., M0212L) for 1 hour at 37°C. Beads are then subjected to three TE-TW washes and loaded into WTA PCR (100 μM Truseq PCR primer: IDT, CTACGACGCTCTTCCGATCT; 100 μM SMART PCR primer: IDT, AAGCAGTGGTATCAACGCAGAGT; Terra PCR mix: Takara Bio, 639284) with cycling conditions 98°C for 2min; 4 cycles of 98°C for 20sec, 65°C for 45sec, 72°C for 3min; 7 cycles of 98°C for 20sec, 67°C for 20sec, 72°C for 3min; and 72°C for 5min. PCR clean up was performed twice with 0.6x SPRI (AmPureXP: Beckman Coulter, Inc., A63881) on cDNA libraries. Final and cDNA libraries were then QCed on a bioanalyzer (Bioanalyzer High Sensitivity DNA kit: Agilent Technologies, Inc., 5067-4626) and qubit (dsDNA high sensitivity kit: Life Technologies, Inc., Q32854) following manufacturer protocols. Tagmentation of 600pg of cDNA is performed according to Nextera DNA sample preparation manufacturer instructions (Illumina, Inc., FC-131-1096) using a Truseq-P5 hybrid constant oligo (IDT, AATGATACGGCGACCACCGAGATCTACACTCTTTCCCTACACGACGCTCTTCCGATCT) and Nextera N7XX indexing primer (Illumina, Inc., FC-131-1001). Final libraries (4nM) were sequenced on a NovaSeq S2 or S4 with 100-200 million reads per sample at the Genomics Platform at the Broad Institute using read structure Read 1 42bp, Index 1 8bp, Read 2 41-60bp, and Index 2 0bp.

### Hybridization Chain Reaction (HCR)

All HCR v3 reagents (probes, hairpins, and buffers) were purchased from Molecular Technologies^92^. Thin sections of tissue (10μm) were mounted in 24-well glass bottom plates (VWR International, LLC, 82050-898) coated with a 1:50 dilution of APTES (Sigma-Aldrich, Inc., 440140). The following solutions were added to the tissue: 10% formalin (VWR International, LLC, 100503-120) for 15min, 2 washes of 1x PBS (ThermoFisher Scientific, AM9625), ice cold 70% EtOH at −20 2 hours to overnight (VWR International, LLC, 76212-358), 3 washes 5x SSCT (ThermoFisher Scientific 15557044, with 0.2% Tween-20), Hybridization buffer (Molecular Technologies) for 10min, probes in Hybridization buffer overnight, 4 15min washes in Wash buffer (Molecular Technologies), 3 washes 5x SSCT, Amplification buffer (Molecular Technologies) for 10min, heat denatured hairpins in Amplification buffer overnight, 3 15min washes in 5x SSCT (DAPI, VWR International, LLC, TCA2412-5MG, 1:10,000, in the second wash), and storage/imaging in 5x SSCT. Imaging was performed on a spinning disk confocal (Yokogawa W1 on Nikon Eclipse Ti) operating NIS-elements AR software. All images acquired were imaged using a Nikon Plan Apo ×40 1.15-NA water immersion objective. Image analysis and processing was performed on ImageJ Fiji.

### Image analysis

Images were first processed using ImageJ2 (National Institutes of Health). Raw ND2 files were background subtracted using the Rolling Ball method (rolling=50 sliding stack). Max intensities of the Z-stack images were then projected, and image channels were split and saved separately. CellProfiler (version 3.1.5, Broad Institute) was then used for cell segmentation based on the fluorescence intensity of DAPI channel and for measuring integrated fluorescence intensity in the rest of the channels (CellProfiler pipeline provided in https://github.com/marshalljamie/Kidney-Slide-seq).

Slide-seqV2 cell percentage and HCR validation plots were generated using Graphpad Prism version 9.1.1. Mean and standard deviation are shown and significance was determined by p<0.05 using a Mann-Whitney U-test. Each dot represents an individual mouse or human sample, except in glomerular plots where each dot represents an individual glomerulus collected from 4 BTBR *wt/wt* or 4 BTBR *ob/ob* mice or an individual human in healthy, DKD, or injured samples.

### Periodic Acid Schiff (PAS) Staining

10μm cryosections of flash frozen tissue were mounted onto superfrost plus micro slides (VWR, 48311-703). Tissue was fixed in 10% formalin (VWR International, LLC, 100503-120) for 15min. Slides were then transferred to the Brigham and Womens Pathology Department for PAS staining, which briefly summarized starts with tissue oxidized in 0.5% Periodic Acid solution for 5min, and rinsed 3 times with distilled water. Slides were then placed in Schiff’s reagent for 15min and washed with tap water for 5min. Slides were counterstained in Mayer’s hematoxylin for 1 min and washed with tap water for 5min and then rinsed with distilled water. Slides were finally dehydrated and mounted using Xylene based mounting media. Imaging was performed on a Zeiss Observer.Z1 microscope using the Zeiss Zen software. Scale bars were added using Fiji ImageJ version 2.1.0/1.53c.

### Cell Type Classification

In order to assign a cell type identity to each Slide-seq bead, we used two methods: (1) NMFreg (Non-negative matrix factorization regression)^1^, and (2) the label transfer method from the R package Seurat (v. 3.0.1)^49^. In the former, a reduced gene space of “metagene” markers for each cell type was identified by non-negative matrix factorization of a single cell reference data set^20^, in which cell types were previously annotated. The reduced gene space for mice and humans was comprised of 40 metagenes and 70 metagenes, respectively, from a single cell reference^20^. Every gene expression profile in the Slide-seq query data set was decomposed into a weighted combination of these metagenes by non-negative least squares regression. Each data point in the query was assigned the cell type corresponding to the metagene of maximum weight. Lastly, cell type annotations were eliminated based on the confidence thresholding protocol defined by the NMFreg developers.

With regards to the Seurat V3 label transfer method, we assigned cell type annotations from the same single cell reference data set^20^ to each of our Slide-seq query data sets. Mappings between highly similar gene expression profiles of the single cell reference^20^ and Slide-seq query data points were established using the FindTransferAnchors() function on SCT-normalized data, with 50 PCs derived from the reference data. Cell type annotations of the query data were acquired from TransferData(), using PCA-derived dimensionality reduction on the reference. This function provided vectors with cell type prediction scores for every bead in our Slide-seq arrays. We defined the cell type identity of every bead to be the cell type class corresponding to the maximum prediction loading.

Upon running both methods on our data and generating scatterplots of the resulting cell type loadings, we found that while podocyte identification was comparable between the two methods, Seurat performed better in confidently identifying all other cell types. We concluded that Seurat cell type calls reliably formed structures across 2-D tissue space that we would expect to find in true kidney biology.

Pronounced tubular structures with fewer scattered points were more consistently characteristic of Seurat’s PCT, DCT, CD-IC, and CD-PC calls than of NMFreg’s calls. Additionally, we consistently found a clear delineation of cortex and medulla regions from Seurat’s TAL calls, which was less often true of NMFreg’s TAL calls (Fig S2-7; Fig S11-12; S17-18).

Consequently, we used podocyte classifications defined by NMFreg and classifications output from Seurat for all other cell types.

### Segmentation of Medulla and Cortex Regions

While most Slide-seq arrays from human kidney sections spanned either the cortex or medulla, several human arrays and all mouse arrays contained both regions. In order to refine our characterization of the cell type and genetic composition of our tissue to these distinct regions of the kidney, we utilized a segmentation algorithm to separate beads in the cortex from beads in the medulla. We first plotted the coordinates of all beads in an array, colored by their cell type class. Visually, we were able to approximate the boundary separating the cortex from medulla because the former contains exclusively PCTs and podocytes, while the latter contains a higher density of DCT, CD-IC, CD-PC, and TAL cell types. Using the methodology adapted from the Python lasso selector widget (https://matplotlib.org/3.1.0/gallery/widgets/lasso_selector_demo_sgskip.html), we were able to hand-draw on these plots our proposed boundary, and in response, the program returned the bead coordinates both within and outside this border.

### Cell Type Curation

While many of the cell types formed dense structural patterns that aligned with the known histology of structures found in kidney tissue, we found relatively isolated instances of cell type calls. We sought to remove these uncertain cell type calls from our analysis. Here, we used a custom method utilizing K-nearest neighbors (KNN). First, we isolated marker beads for each structure, whether that be beads having a certain cell type label or expression of a marker gene, labeling these 1, and labeled all other beads 0. Next, from the Python package scikit-learn (v. 0.23.1)^93^, we generated the KNN adjacency matrix based on the coordinates of all beads with the function NearestNeighbors(), using the ball tree algorithm and user-specified k. We then filtered all marker beads with fewer than a threshold number of 1-labeled neighbors. Lastly, we added back in any nearest neighbors of our isolated beads having a label of 1. The motivation here was that marker beads occurring along the edge of structures may be filtered out because their composition of surrounding 0-labeled neighbors is too high.

The number of k-nearest neighbors we allowed the algorithm to search for, and the threshold number of neighbors used to filter out points were chosen by trial and error in order to maintain a clean boundary around the known biological structures of each cell type.

### Glomerulus, GC, and MD Detection

Because podocytes are known to be solely contained within glomeruli, and glomeruli have a circular shape in 2-D tissue space, we used our KNN-filtration method on podocyte-annotated beads. We chose parameters that maintained these circular structures, while throwing out spurious, isolated points (Fig S53). We then assigned other glomerulus-specific cell types, including mesangial cells and endothelial cells, to these regions containing podocytes. To do so, we computed the circular area of each glomerulus defined by its approximate center and radius. The centers were defined as the cluster centers of curated podocyte coordinates output by cluster.KMeans() from the Python package scikit-learn. Here, K was determined by the number of podocyte clusters visualized in a scatter plot of podocyte coordinates. We approximated the radius of each glomerulus to be the maximum of the distances between all podocytes in a cluster and its cluster center. We then assigned instances of endothelial and mesangial-annotated beads to a podocyte cluster if their coordinates occurred within the circle defined by its radius and center (Fig S60).

GC clusters were identified by running KNN-filtration on all beads expressing *Ren1*, or *REN*, with parameters chosen to isolate elliptical, dense groupings of these points. Although we know that GC cluster near glomeruli, our curated granular cell regions disagreed with this assumption. Therefore, we hand-selected GC clusters that occurred within 250 pixels to a curated glomerulus (Fig S54). Approximately 39% of GC clusters met this criterion.

In order to identify MD structures, we iterated through several filtration steps. First, we ran our KNN-filtration method on TAL, glomerulus, and granular cell-annotated beads, aiming to hone in on any TAL-annotated beads that occur near clusters of glomerulus-specific cells or GC, whether they themselves occur in a dense cluster or not. We then eliminated any of our filtered TAL-annotated beads that expressed the gene *Slc12a3* - a gene characteristic of TAL, but not MD structures. Lastly, in mice we hand-filtered the remaining beads based on the expression of *Ptger3/PTGER3, Klf6/KLF6*, and *Nos1/NOS1*, or vicinity to either a glomerulus or granular cell structure (within 250 pixels) (Fig S55). Ultimately, we selected approximately 22% of the MD structures curated with KNN.

Lastly, we assigned unique cell type labels to the beads having more than one label, with the following criteria. Amongst beads classified as both GC and glomerulus-specific cells, we found that the overlapping beads on average had higher expression of Ren1 than the rest of the granular cell population (Fig S52). As a result, overlapping beads maintained their glomerulus cell type assignments. Beads classified as both GC and MD cells were assigned to the group in which they had the higher gene expression marker.

### PCT, DCT, and CD-PC Detection

We curated PCT, DCT, and CD-PC-annotated beads in a similar manner. Polygons encapsulating glomeruli, granular cell clusters, and MD clusters were identified with alpha = 0.01, and all instances of curated CD-IC, CD-PC, DCT, and PCT beads were removed from their areas. On calls of DCT and CD-PC in the cortex and medulla separately and all PCT, we ran our KNN-filtration method, selecting parameters that maintained beads contained within tubule structures. Additional filtering was done with the Python package alphashape (v. 1.0.2) (https://alphashape.readthedocs.io/en/latest/), which output a polygon encapsulating most of the beads of the cell type of interest. All beads outside of the alphashape were filtered. Remaining PCT beads within our proposed medulla region of each array were removed (Fig S56-58).

### CD-IC Detection

We first combined CD-A-IC and CD-B-IC cell calls and called them CD-IC. Because it is possible to find singular CD-IC beads that occur within CD-PC tubular structures, CD-IC isolation was done by concatenating CD-IC beads with curated CD-PC beads in the cortex and medulla separately, and running KNN-filtration. Following this step, we aimed to remove remaining CD-IC beads that occurred within dense clusters outside of CD-PC tubules. We further filtered any instances of CD-IC beads that were located further than a distance of 100 pixels from the polygon surrounding CD-PC tubules using the distance() method from the Python package Shapely (v. 1.7.0) (https://github.com/Toblerity/Shapely) (Fig S59).

Polygons encapsulating glomeruli, granular cell clusters, and MD clusters were identified with alpha = 0.05, and all instances of curated CD-IC, CD-PC, DCT, and PCT beads were removed from their areas. TAL beads were removed only from glomerulus and granular cell areas.

### Immune Cell, Fibroblast, and vSMC Detection

Because our human single cell reference dataset^20^ had only an immune cell cluster, we parsed out macrophages from our immune-annotated beads by identifying those that expressed either *C1qa/C1QA* or *C1qb/C1QB*^51^.

Because fibroblasts, vascular smooth muscle cells, and immune cells do not have a clear underlying structure, we did not use KNN-filtration here.

TAL beads were removed only from glomerulus and granular cell areas.

### Spatial Outlier Detection

Abundant cell types such as PCTs, TAL, and endothelial cells that reach the edges of arrays sometimes spilled over outside of the array area. Because these distant beads are no longer associated with coordinates across tissue space, they were deemed spatial outliers and eliminated from the analysis. Detection of these outliers was accomplished by the anomaly detection algorithm ensemble.IsolationForest() from the Python package scikit-learn. For every Slide-seq array, all raw coordinates were fed into the isolation forest algorithm, and outliers were thrown out. Lastly, curated beads from each cell type class were merged with the filtered set of coordinates.

### Cell Type Structure Assignment

Certain cell types formed discrete, classifiable groupings across arrays. CD-IC, CD-PC, and DCT formed tubular islands, GC and MD cells formed elliptical structures, and mesangial cells, endothelial cells, and podocytes comprised circular glomeruli. Formulating structure aggregates were motivated by two downstream analyses: (1) morphology quantifications and (2) transcriptomic characterizations.

First, we identified individual structures with cluster.KMeans(), from the Python package scikit-learn, on the coordinates of each curated cell type. With podocytes, mesangial cells, and endothelial cells within glomeruli, GC, and MD cells, K was determined to be the number of dense clusters of points visualized in a scatter plot of curated coordinates. For CD-PC, and DCT, where each cell type comprises a single biological structure, but is characterized by natural partitions, K was determined by a silhouette analysis. Here, we searched for a K that maximizes the difference between minimum average out-of-cluster distances and average within-cluster distances. For every k in {2,…,max(number of beads, 50)}, we computed metrics.silhouette_score() of cluster.KMeans(K=k) from the Python package scikit-learn. We ultimately used the K-means clustering results with K being the number of clusters corresponding with the maximum silhouette score. CD-IC beads were assigned to their nearest CD-PC cluster (Fig S34-51).

### Quantifying the Morphology of Glomeruli

In order to address the hypothesis that glomerular morphology differs between the disease states of kidney tissue, we computed the following morphology metrics for every glomerulus: (1) area, (2) percentage of podocytes, (3) percentage of mesangial cells, and (4) percentage of endothelial cells. In order to compute the area of glomeruli, the convex hull of each glomerulus was found with the Python package alphashape (v. 1.0.2), using the function alphashape() with alpha = 0. This step returned a convex polygon enclosing all points in every glomerulus, along with an area attribute.

The percentages of podocytes, mesangial cells, and endothelial cells in glomeruli were computed with the following protocol: for every glomerulus, we divided the number of beads classified as the cell type of interest by the total number of beads in the glomerulus. Statistical tests were performed with the Mann-Whitney U-test.

### Cell Type Proportion Quantification

In order to address whether there was a significant difference in the cell type composition of kidneys in diseased versus wild-type kidney tissue, we computed the proportion of every cell type in the medulla, cortex, and whole kidney in each of our arrays. That is, for every cell type in every region, we divided the number of beads classified as this cell type by the total number of beads in the region of interest or entire array. We then compared these quantifications between UMOD-KI and WT mice, BTBR *ob/ob* and BTBR *wt/wt* mice, and the healthy, DKD, and injured humans. Statistical tests were performed with the Mann-Whitney U-test (Fig S13, S19, S32-33).

### Transcriptomic Data Preprocessing

We conducted quality control by computing the number of genes and UMIs per array and UMIs per cell type were quantified (Fig S30-31). Prior to running our transcriptomic analyses, we applied normalization and batch effect removal methods to our data, all done using the R package Seurat. In order to account for sequencing depth per bead, we normalized our data using the SCTransform() function^50^ on each array. Next, we combined the transcriptomic data in arrays with genotypes or disease states we planned on comparing (e.g., BTBR *wt/wt* and BTBR *ob/ob* mice, WT and UMOD-KI mice, and all humans) into a single data matrix. We ran SelectIntegrationFeatures() to find the top 3000 consistently variable features in the sctransform data across all arrays in each comparison grouping. SCTransform() was then run across all the data in each group at once, this time using the shared variable feature space discovered in the previous step.

In order to remove batch effects, within each genotype or disease state grouping of mice and humans, we searched for differentially expressed genes across batches. We used the function FindAllMarkers() with the negative binomial test on the raw counts, filtering for genes characterized by a log fold-change of at least 0.05 and occurring in a minimum fraction of 0.05 of the beads in each of the batch groups under comparison. The union of the sets of batch markers derived from disease states that we intended on comparing downstream were removed from the data (Fig S16, S21-22).

### Disease State Clustering and Differential Expression Analysis

With regards to glomerulus-specific cell types, DCT, CD-IC, and CD-PC, where we had clusters of cells forming small, discrete structures, we decided to cluster on the structures rather than cells. Because Slide-seq beads don’t always capture mRNA purely from a single cell type, we reasoned that in smaller structures, surrounded by a mixture of other cell types, edge beads would be prone to picking up mRNA from external cell types. To remove some of this noise, we averaged the sctransform residuals of the gene expression profiles of beads belonging to each of our previously-defined structures. Gene profiles consisting of sctransform residuals of cell types that did not form these smaller, discrete structures, such as TAL, vSMC, fibroblasts, macrophages, and other immune cells were not aggregated in this way.

For dimensionality reduction, we performed uniform manifold approximation and reduction (UMAP) with the Python package umap-learn (v. 0.4.6)^94^. We chose the clustering algorithm that best fit the biological labels of our data in UMAP-space: hdbscan (v. 0.8.26)^95^ in the cases where we clustered on cell type aggregates, and cluster.SpectralClustering() from scikit-learn in the cases where we clustered on the gene expression profiles of individual beads.

Clusters were assigned a disease status based on the composition of points contained within them. That is, if a cluster contained > 70% structure aggregates or beads coming from diseased arrays, all data points within it were labeled as diseased (Fig S16, S21-22).

Differential expression across clusters was performed with the function scanpy.tl.rank_genes_groups() from the Python package scanpy^96^, using the Wilcoxon rank-sum test and filtering for the top 100 differentially expressed genes in each cluster. Genes were called differentially expressed if their adjusted p-value, computed with the Benjamani-Hochberg correction method, fell below 0.05. For each DE gene, the average log fold-change was computed from the sctransform corrected counts, defined as the log10 transform of the difference in the average expression of the gene between the two clusters, added by one.

### Macrophage–Cell Neighbor Analysis and Statistics

In order to explore cell neighborhoods and cell-cell interactions between macrophages and other cell types, we used a custom analysis. For every macrophage bead of interest, we looked at all of its neighbors within a radius of 25 pixels using the function neighbors.radius_neighbors_graph() from the Python package scikit-learn. For every array, we were then able to compute the spatial interaction of a cell type, A, with a cell type, B, as the number of times A is a neighbor of B, normalized to the square-root of the product of the total number of cells A and total number of cells B. This normalization accounted for overall frequency of each cell type and all possible interactions between the two cell types of interest across arrays.

In order to see if the cell-cell interaction neighborhoods were unique, we controlled for all other cell types. For humans, to determine if interactions between *LYVE1+* macrophages versus their newly defined neighbors were unique, we also looked at cell interaction frequencies of *all* macrophages versus *all other cell types* in the medulla. For mice, to determine if interactions between all macrophages and their newly defined neighbors in the medulla were unique, we also looked at cell interaction frequencies of all macrophages versus all other cell types in mouse cortex.

In order to define differences between healthy and injured tissue, we performed further statistical analysis. For humans, cell type interaction frequencies for all *LYVE1+* macrophages versus all cells, and all macrophages versus all cells were compared across healthy and injured medulla. For mice, cell type interaction frequencies for all macrophages versus all cells were compared across the cortex and medulla in UMOD-KI and UMOD-WT mice. The statistical test used to compare interaction frequencies between healthy and disease tissue was the Mann-Whitney U-test, correcting p-values with the Benjamani-Hochberg method.

These new analyses are now presented as dot plots showing relative proportions of cell types normalized across healthy and disease tissue. Specifically, dot plots were generated showing the interaction frequency of macrophages (all macrophages or a subset, i.e. *LYVE1+*) versus every other cell type in the tissue array. The color of the dots indicates the intensity of the interaction (red, increased; blue, decreased interaction). The size of the dots indicates the adjusted p-value for cell-cell interactions and significance is shown by a black outline around the dot (adjusted p-value < 0.05). Cell types not represented in the dot plot had no interactions with macrophages of interest.

### Glomerulus and Granular Cell Distance Computation

In order to analyze the spatial relationship between glomeruli and GC, we aimed to compare the distances between the two structures across genotypes in mice and used two different methods. In the first method, using the previously-computed K-means cluster centers, we computed the Euclidean distances between the cluster centers of all glomeruli and granular cell structures in every array. We then searched for glomerulus-granular cell structure pairs at a minimum distance to each other. Distances were then compared between BTBR *wt/wt* and BTBR *ob/ob* mice. In the second method, we computed the minimum Euclidean distances between the edges of the convex hulls encompassing glomeruli and granular cell structures using the distance() method from the Python package Shapely. The distribution of distances was then compared between arrays of BTBR *wt/wt* and BTBR *ob/ob* mice. Statistical tests were performed with the Mann-Whitney U-test.

### UPR Pathway Analysis in UMOD Mice

A list of UPR-specific genes was acquired from Adamson *et al*., 2016^79^. Genes appearing in < 50% of the UMOD-KI and WT transcriptomic data matrices were removed. The select set of genes were then subset from the raw transcriptomic data of each of our UMOD-KI and WT mouse arrays and all data was combined into a single matrix. Beads expressing at least one feature of the UPR signature were maintained for the rest of the analysis. In order to account for sequencing depth of each bead, we ran the SCTransform() method from the R package Seurat. Next, we averaged the sctransform-normalized gene expression profiles of beads classified as being TAL in the medulla, per mouse. Differential expression was performed across aggregated gene expression profiles of TAL-medulla beads in UMOD-KI and WT mice. Here, we used the function scanpy.tl.rank_genes_groups() from the Python package scanpy with the Wilcoxon rank-sum test. We called genes differentially expressed if their adjusted p-value, computed with the Benjamani-Hochberg method, fell below 0.05. In order to visualize these gene expression differences across UMOD-KI and WT mice, we used the method clustermap() from the python package seaborn (v. 0.10.1)^97^, performing min-max normalization for every gene. The same protocol was followed in order to conduct DE across TAL-cortex and PCT beads in UMOD-KI and WT mice as negative controls.

## Acknowledgments

We thank Rajesh Thakker at the University of Oxford for providing the UMOD-KI mice used in this study, Sophia Liu for discussions and advice on immune cells, Pawan Kumar for generation of beads used in the arrays, Jilong Li for developing the computational pipeline used to map sequencing data to spatial locations, Dylan Cable for advice on RCTD, Aleks Goeva for advice and code for mapping cell types with NMF-reg, Terri Woo for performing PAS staining on kidney tissue, Michael Howard for mouse husbandry and treatments, and Jillian Shaw, Juanchi Pablo, and Aviv Regev for comments on the manuscript. The authors thank all members of the Greka and Chen labs for comments and suggestions.

## Author contributions

JLM conceived of and designed the study, collected mouse tissue, performed all slide-seqV2 and HCR experiments, supervised data analysis, drafted figures, and wrote the manuscript. TN performed data analysis for slide-seqV2 data, drafted figures, and wrote the manuscript. QSW supervised and performed data analysis for slide-seqV2 data, provided high level direction for data analysis, and wrote the manuscript. HC developed computational methods for both HCR and slide-seqV2, performed HCR image analysis, and wrote the manuscript. EM generated all slide-seqV2 arrays. KAV assisted with procurement of human samples, AS performed computational analysis of human and BTBR *wt/wt* mouse single cell data used for mapping cell types on slide-seqV2 data, and KAV and AS contributed to findings on *LYVE1+* and *Trem2+* macrophages from single cell analysis of human and mouse kidneys. KL designed immune cell interaction schematics. RRS performed experiments using the original slide-seq version 1 protocol. BM performed analysis of NMF-reg cell mappings and cell type interactions. RMH collected images of PAS stained tissue. KK and AW assisted with procurement of human samples. EZM provided Slide-seqV2 arrays for this study. FC provided Slide-seqV2 arrays for this study, assisted with study design and implementation, and provided input on results. AG supervised the study and wrote the manuscript. All authors reviewed and provided input on the manuscript.

