## Supplemental Figures and Tables for "High Resolution Slide-seqV2 Spatial Transcriptomics Enables Discovery of Disease-Specific Cell Neighborhoods and Pathways"

**This PDF file includes:**

Figs. S1 to S65

Tables S1 to S14

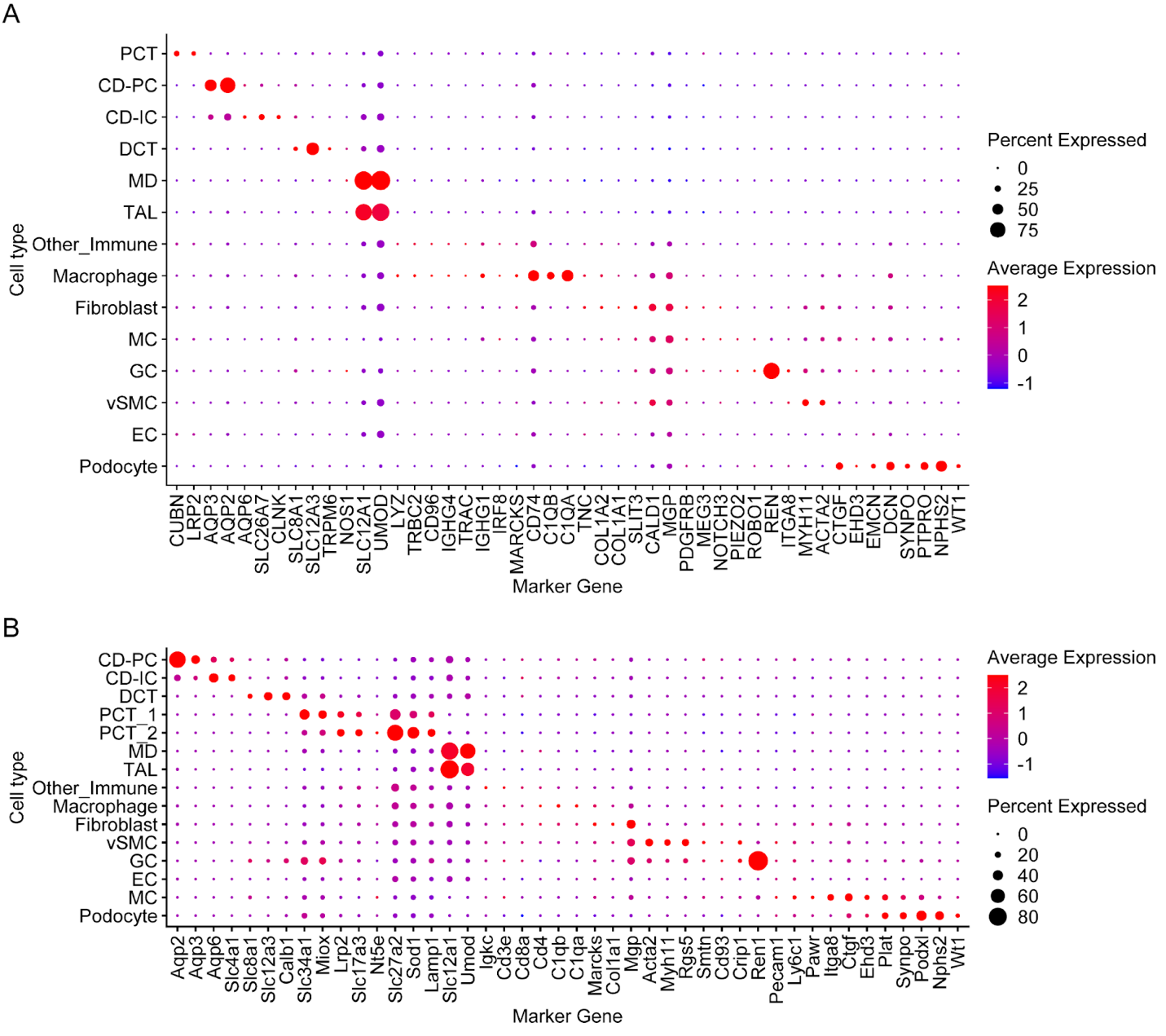

**Fig. S1. Cell type marker genes in human and mouse arrays.**

Dot plot showing marker gene expression in each cell assignment on slide-seq beads in (**A**) human and (**B**) mouse arrays.

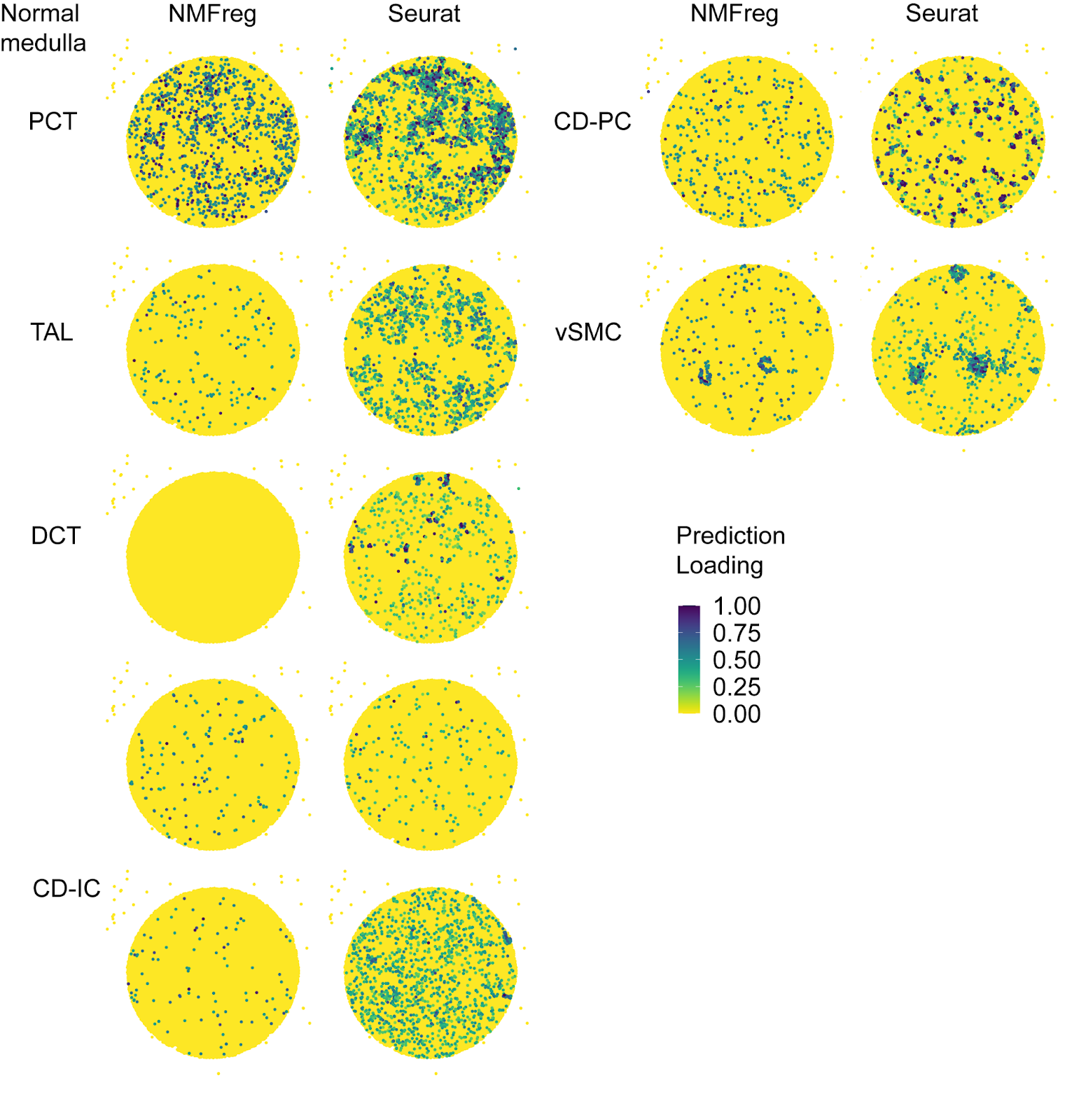

**Fig. S2: Comparison of NMFreg and Seurat cell mappings in human healthy medulla array.** Spatial distribution of cell type prediction loadings from NMFreg (left) and Seurat transfer learning (right) methods across a human healthy medulla array.

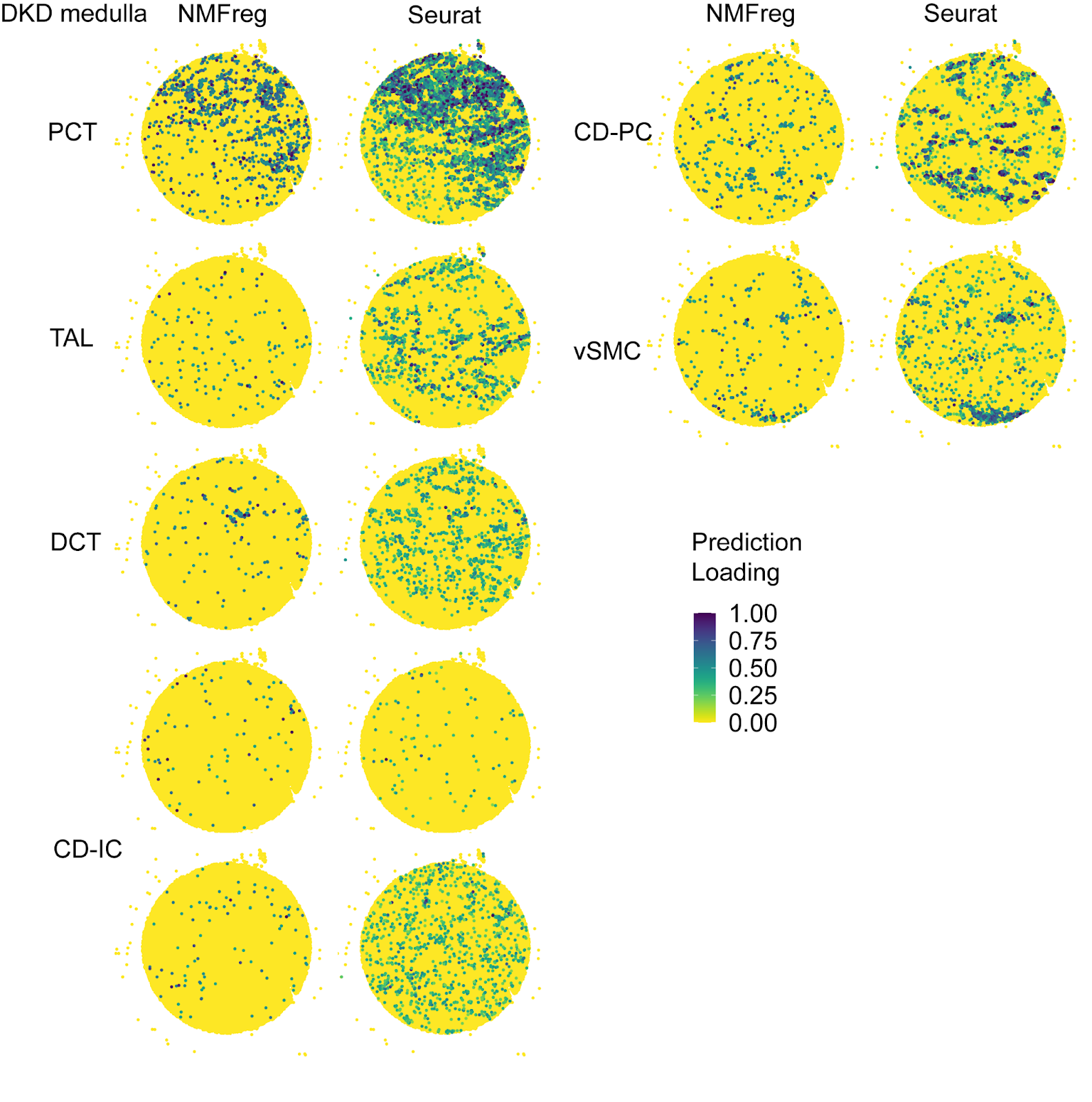

**Fig. S3. Comparison of NMFreg and Seurat cell mappings in human DKD medulla array.** Spatial distribution of cell type prediction loadings from NMFreg (left) and Seurat transfer learning (right) methods across a human DKD medulla array.

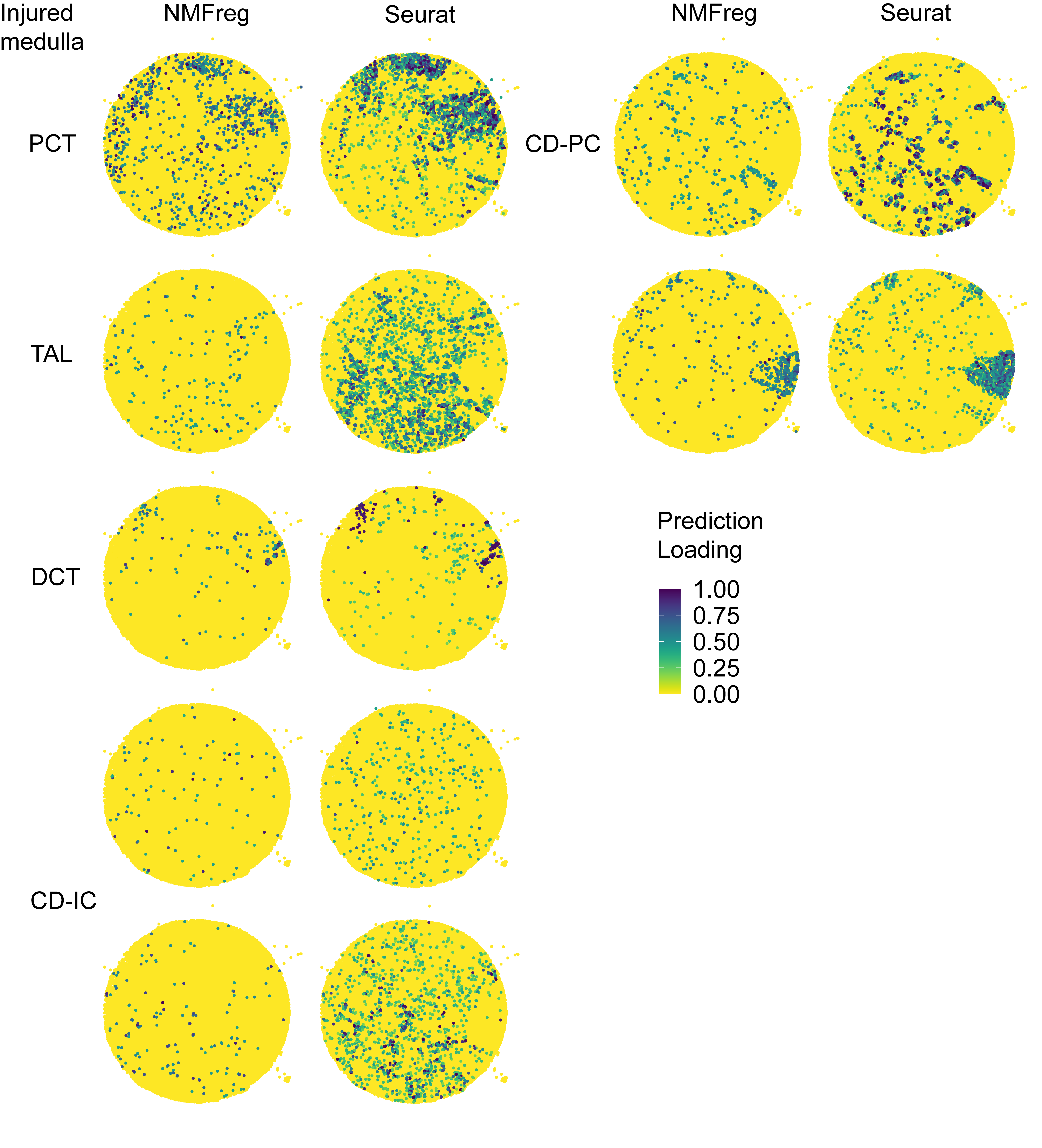

**Fig. S4. Comparison of NMFreg and Seurat cell mappings in human injured medulla array.** Spatial distribution of cell type prediction loadings from NMFreg (left) and Seurat transfer learning (right) methods across a human injured medulla array.

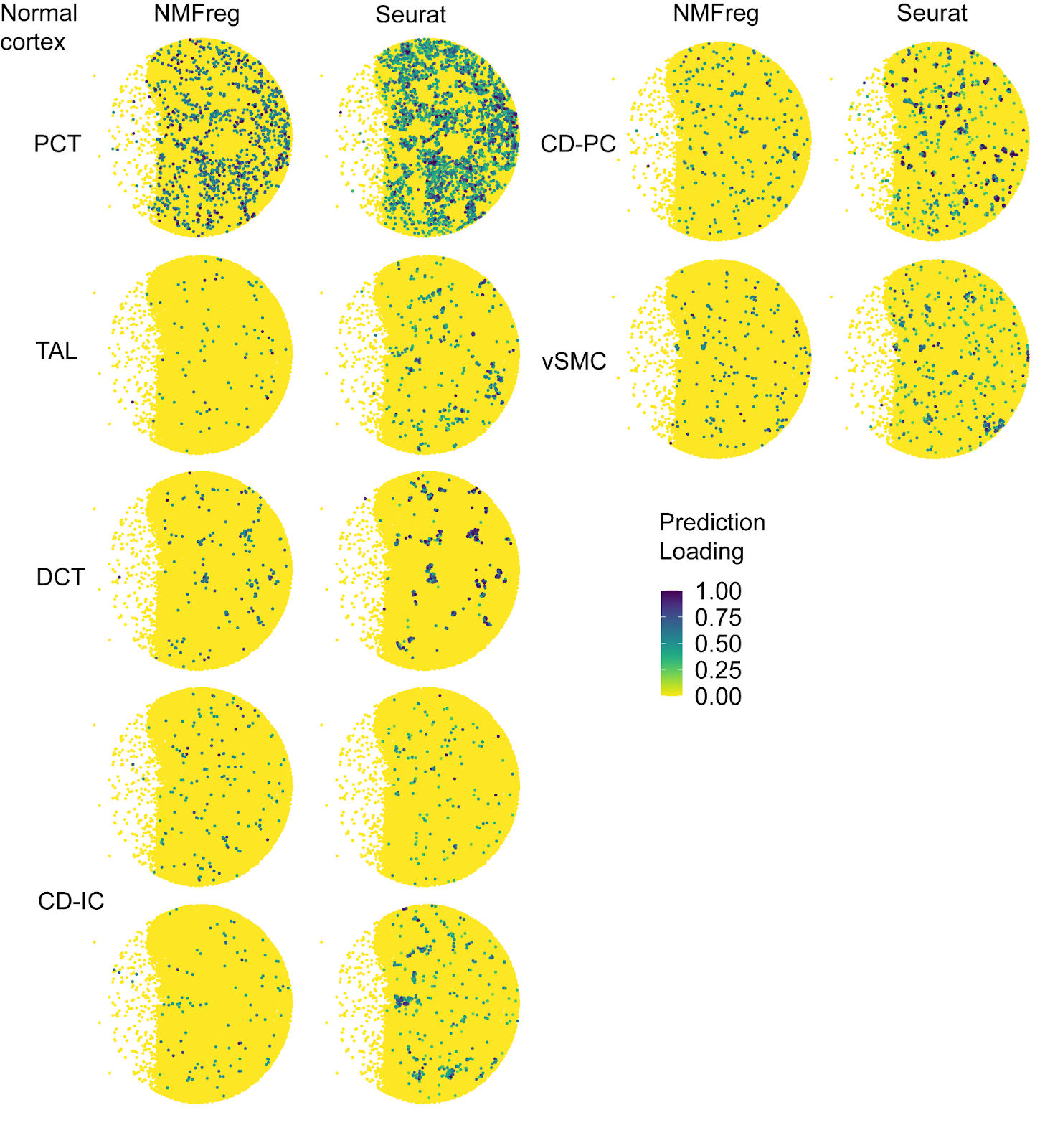

**Fig. S5. Comparison of NMFreg and Seurat cell mappings in human healthy cortex array.** Spatial distribution of cell type prediction loadings from NMFreg (left) and Seurat transfer learning (right) methods across a human healthy cortex array.

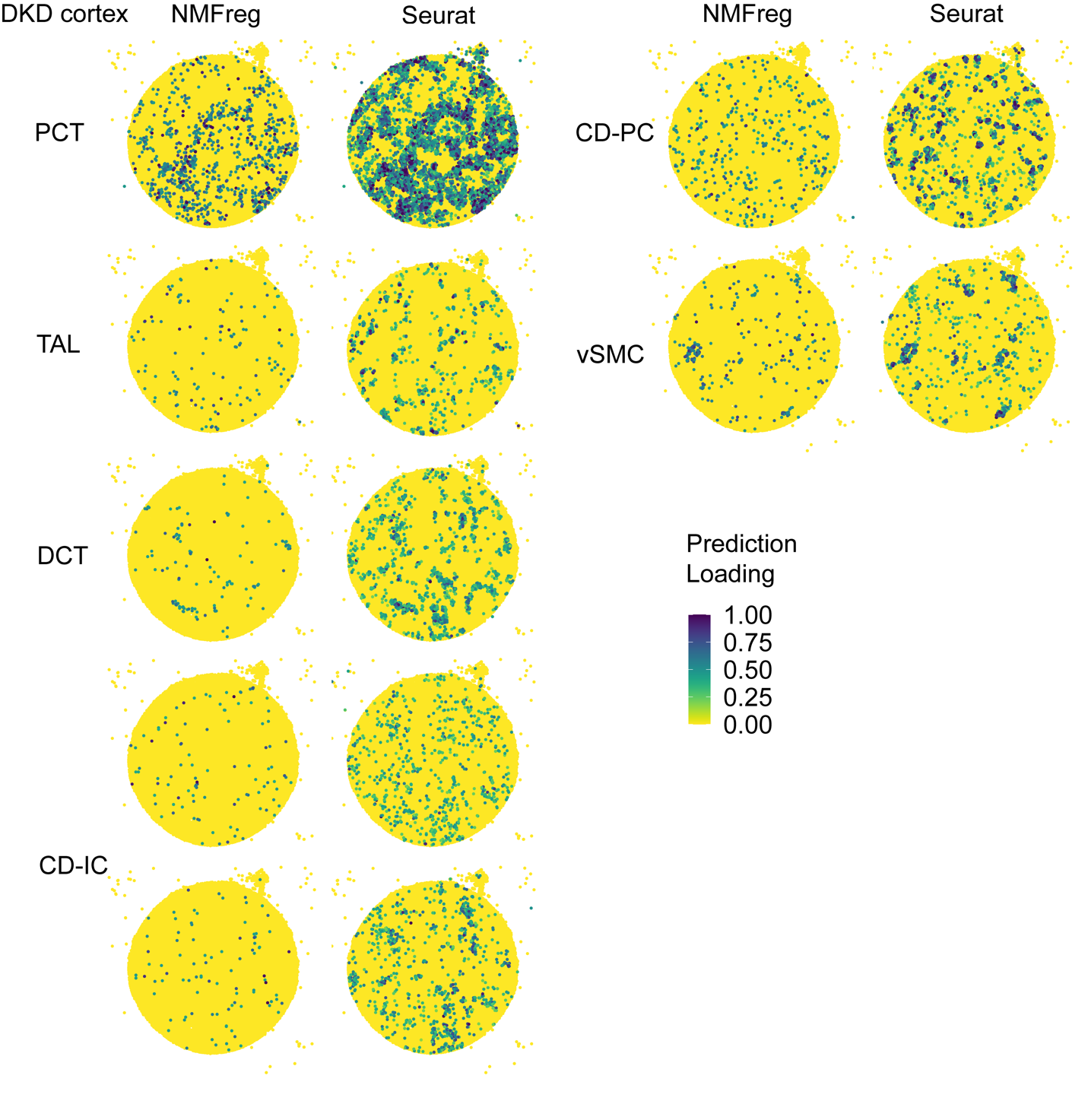

**Fig. S6. Comparison of NMFreg and Seurat cell mappings in human DKD cortex array.** Spatial distribution of cell type prediction loadings from NMFreg (left) and Seurat transfer learning (right) methods across a human DKD cortex array.

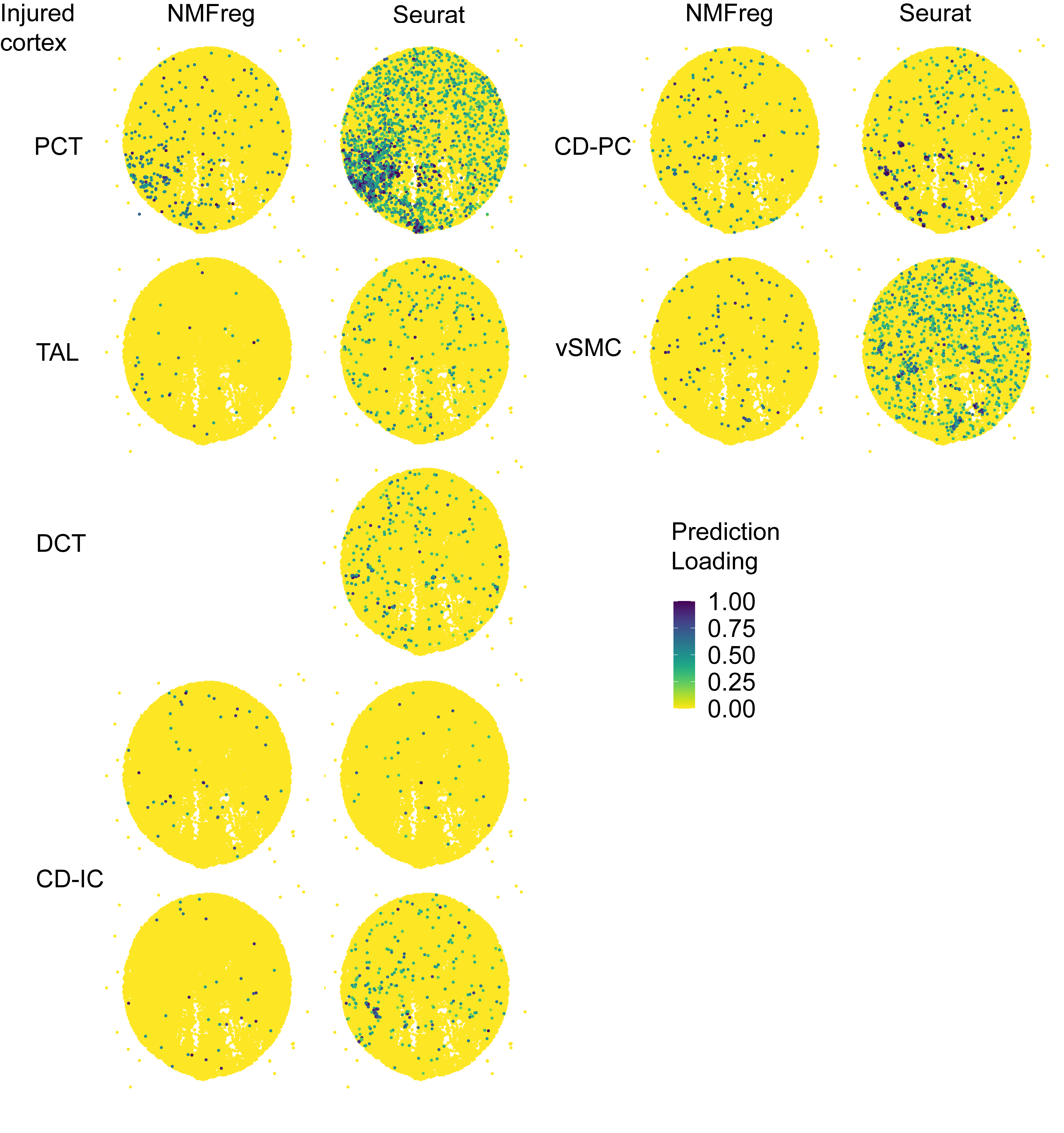

**Fig. S7. Comparison of NMFreg and Seurat cell mappings in human injured cortex array.** Spatial distribution of cell type prediction loadings from NMFreg (left) and Seurat transfer learning (right) methods across a human injured cortex array.

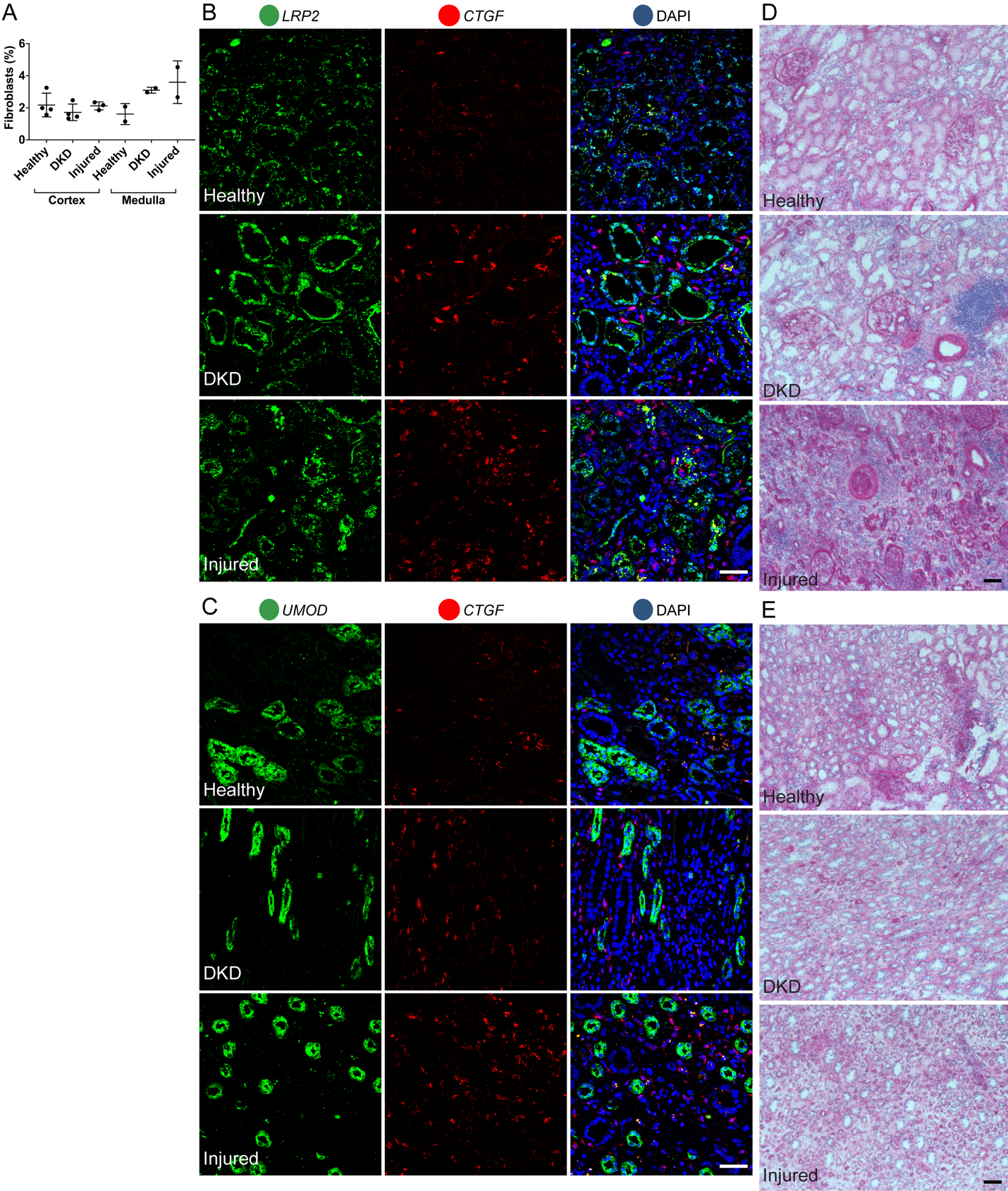

**Fig. S8. Ischemia results in an increase in fibroblasts in the medulla.** (**A**) Plot displaying the percentage of fibroblast beads in cortex and medulla arrays. Each point is an array from 1 human biopsy sample. All error bars represent standard deviation of the mean. HCR images of *CTGF*+ cells in the (**B**) cortex (*LRP2*) and (**C**) medulla (*UMOD*). Scale bar 50µm. DKD acts as a control in this experiment. PAS images from (**D**) cortex and (**E**) medulla. Scale bar 100µm.

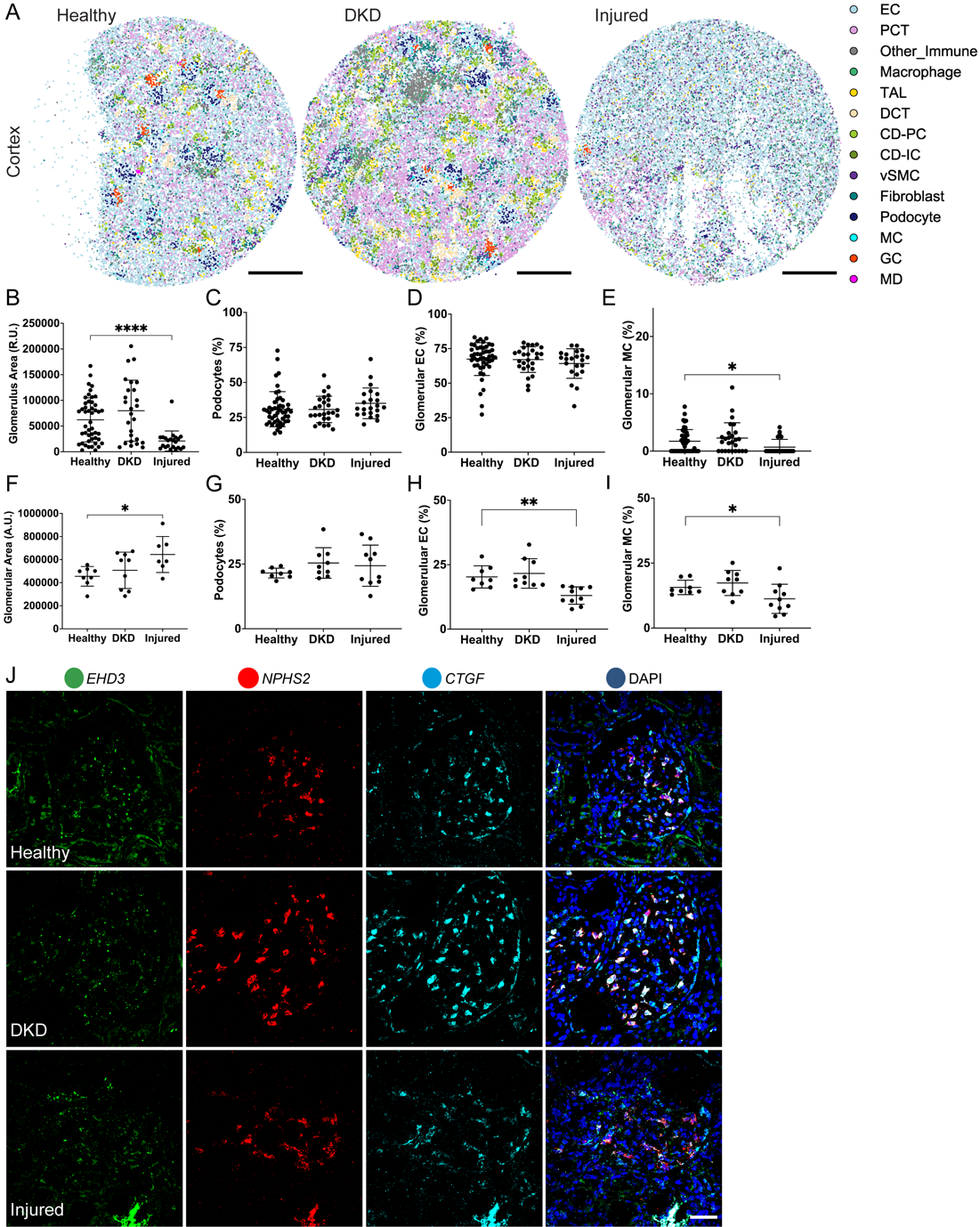

**Fig. S9. Glomeruli not affected in kidneys with early DKD or ischemia.** (**A**) Arrays displaying cell types in healthy, DKD, and Injured biopsies from the cortex. Images of individual cell populations are plotted in Fig S61-63. Scale bars, 500µm. Plots from slide-seq arrays showing (**B**) average area of individual glomeruli, (**C**) percentage of beads classified as podocytes, (**D**) percentage of beads classified as glomerular endothelial cells, and (**E**) percentage of beads classified as glomerular mesangial cells. Plots generated from HCR of (**F**) average area of individual glomeruli, (**G**) percentage of *NPHS2*+ podocytes within the glomerulus, H) percentage of *EHD3*+ glomerular endothelial cells, and (**I**) percentage of *CTGF*+ glomerular mesangial cells*.* Each data point is a structure from 1 human biopsy. (**J**) HCR images showing *EHD3*+ endothelial cells, *NPHS2*+ podocytes, *CTGF*+ mesangial cells, and all cells in DAPI. The absence of injured in early DKD led us to not pursue further analyses in this sample. Scale bar 50µm. All error bars represent standard deviation of the mean.

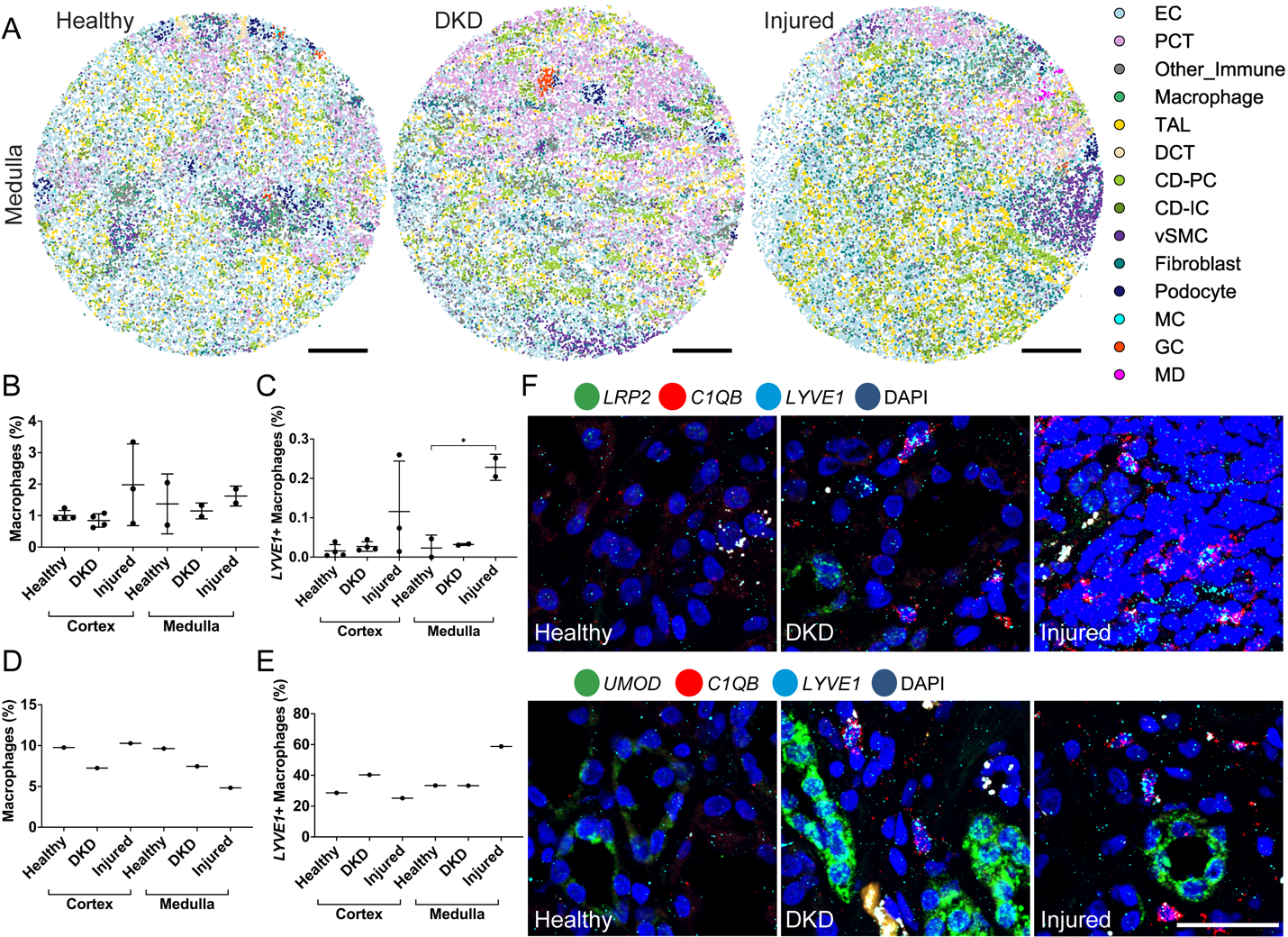

**Fig. S10. Spatial transcriptomics in injured human kidney reveals *LYVE1+* macrophage expansion that changes the composition of a cell neighborhood in the medulla**. (**A**) Medulla arrays from healthy (left), DKD (middle), and injured (right) tissue showing mapping of cell types. Images of individual cell populations are plotted in Fig S61-63. Scale bars, 500µm. (**B**) Plot displaying the percentage *C1QB*+ *C1QA*+ macrophage beads in cortex and medulla arrays. Each point is an array from one human kidney sample. (**C**) Plot displaying the percentage *LYVE1*+ macrophage beads in cortex and medulla arrays. *LYVE1*+ macrophages are increased in the medulla of Injured arrays (p-value 0.0249, Welch’s t-test). Each point is an array from one human kidney sample. (**D**) HCR validation of percentage *C1QB*+ cells in cortex and medulla arrays. One human kidney sample. (**E**) HCR validation of percentage *LYVE1*+ *C1QB*+ double positive cells in cortex and medulla arrays. *LYVE1*+ macrophages are increased in the medulla of injured arrays. One human kidney sample. (**F**) HCR images of *LYVE1*+ *C1QB*+ double positive cells in cortex (top, *LRP2*) and medulla (bottom, *UMOD*). Scale bar 50µm. All error bars represent standard deviation of the mean.

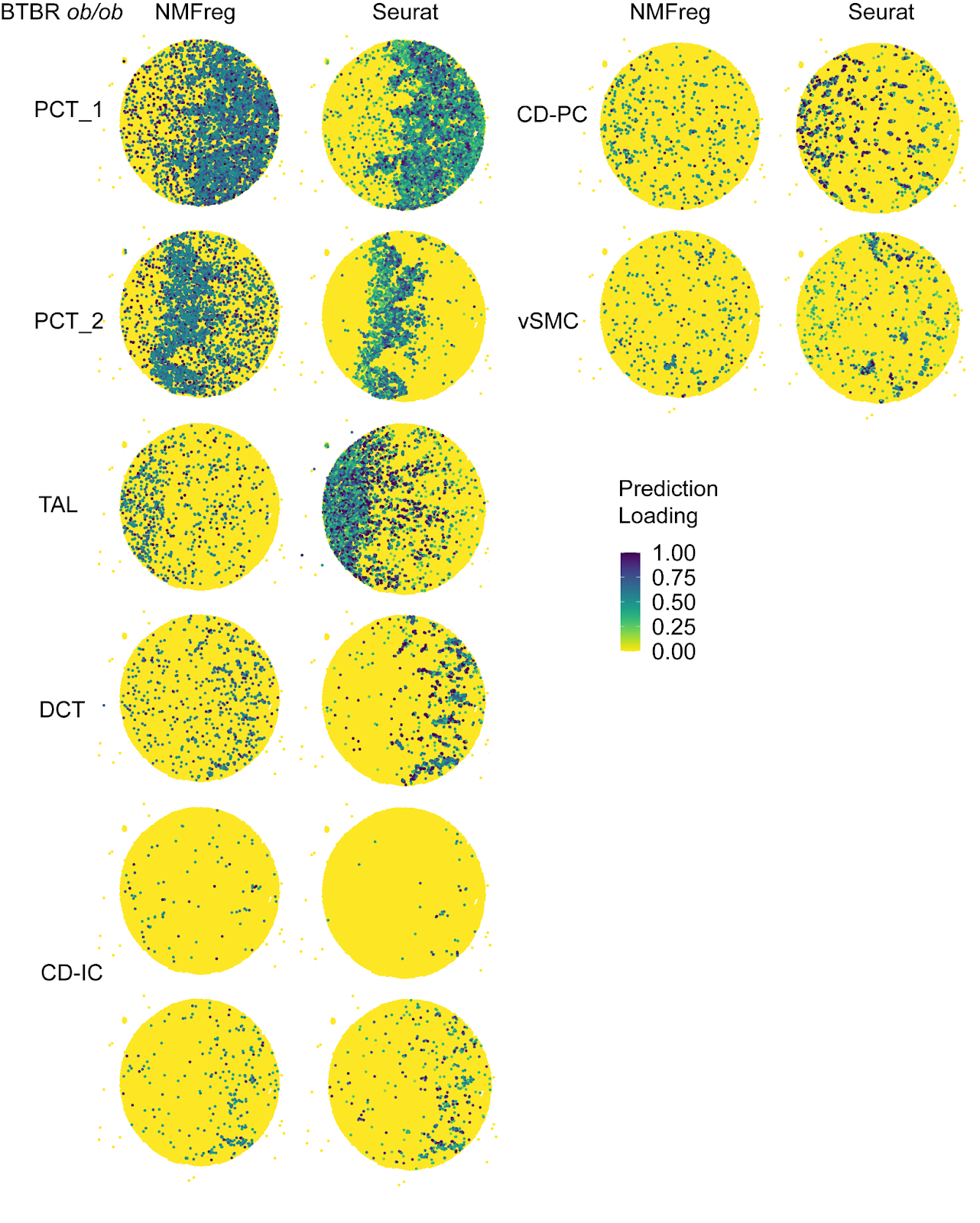

**Fig. S11. Comparison of NMFreg and Seurat cell mappings in BTBR *wt/wt* control mice.** Spatial distribution of cell type prediction loadings from NMFreg (left) and Seurat transfer learning (right) methods.

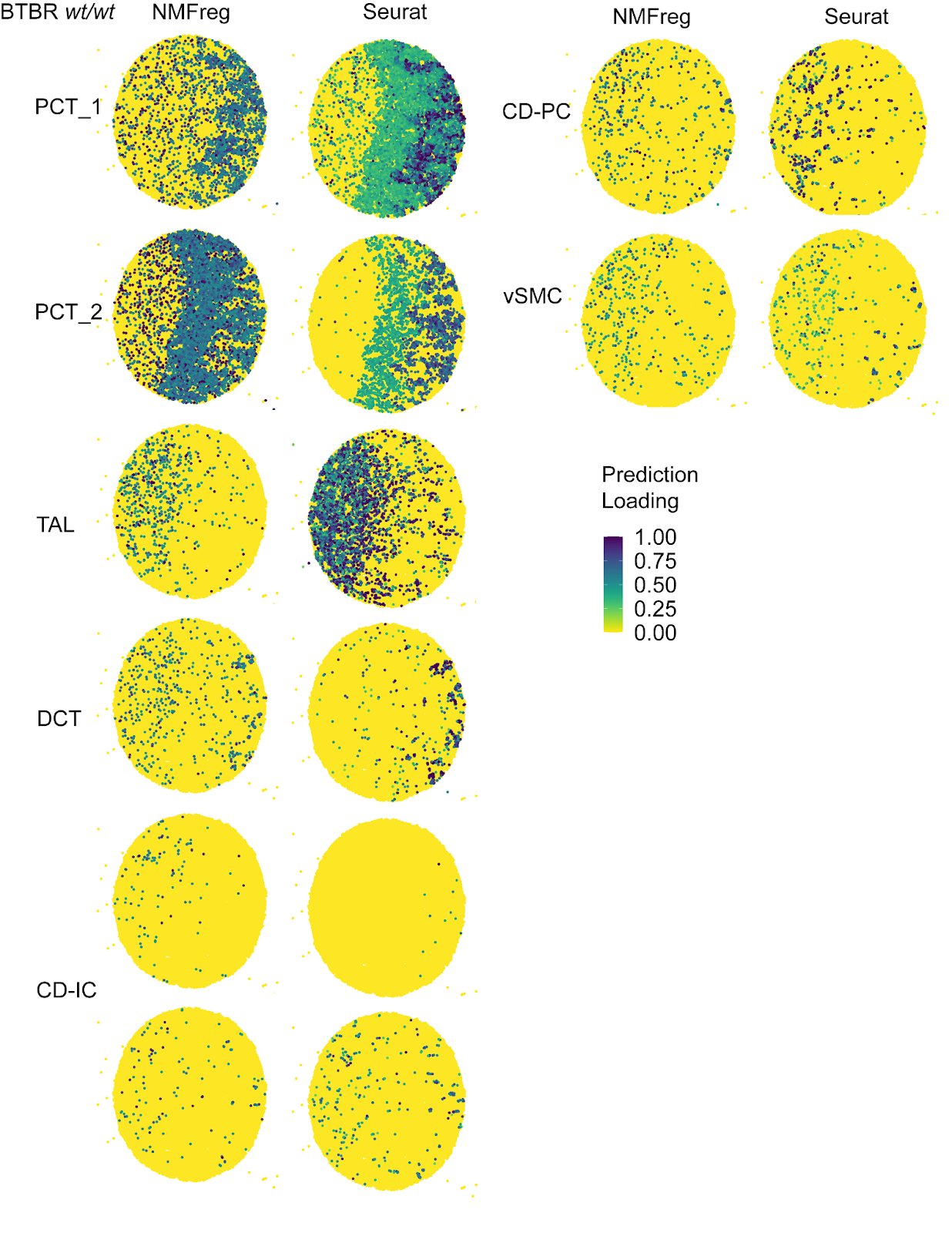
**Fig. S12. Comparison of NMFreg and Seurat cell mappings in BTBR *ob/ob* mice.** Spatial distribution of cell type prediction loadings from NMFreg (left) and Seurat transfer learning (right) methods.

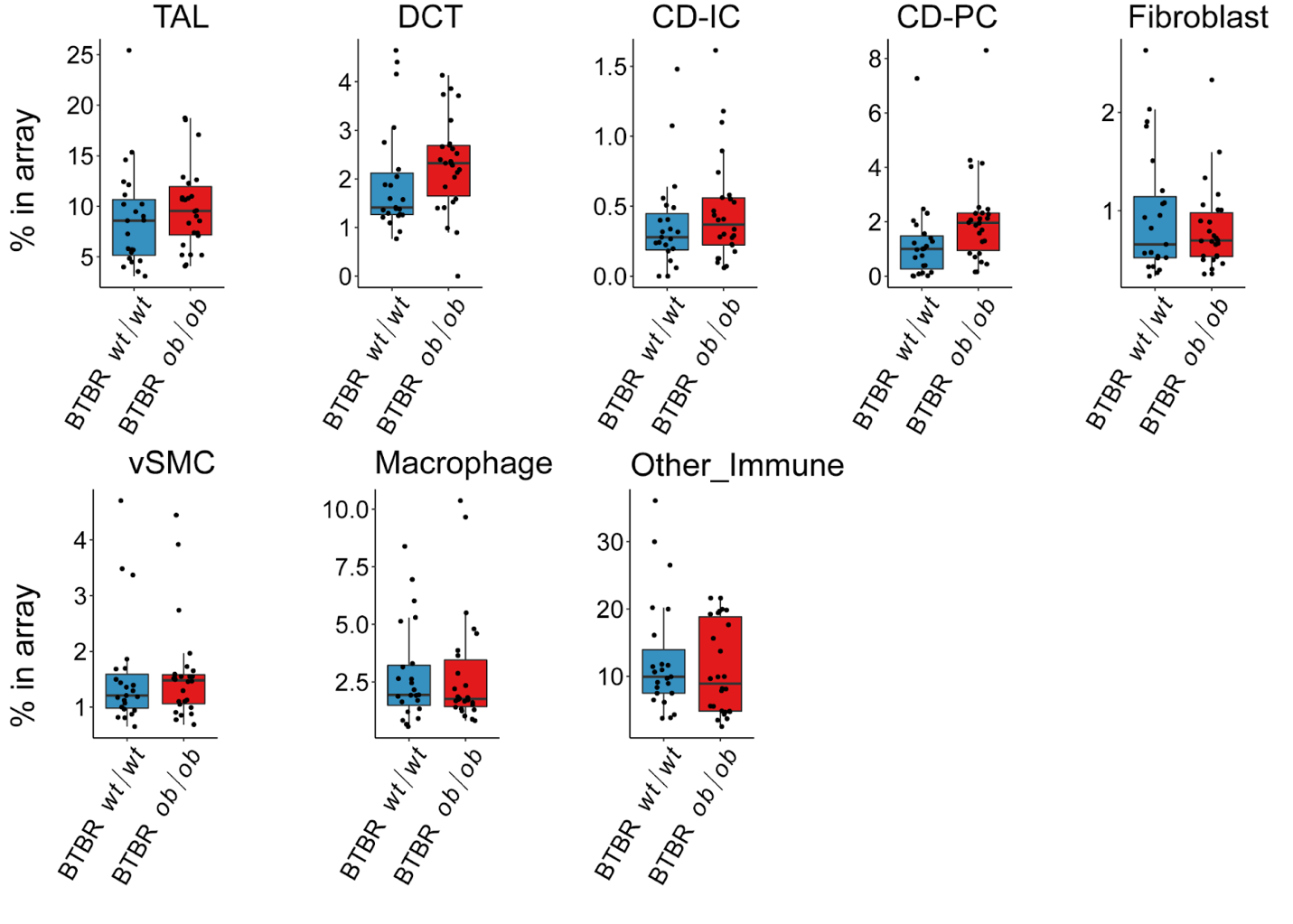

**Fig. S13. Cell type percentages across arrays from BTBR *wt/wt* and BTBR *ob/ob* mice.** Percentage of arrays classified as non-glomerular cell types. Lower, middle, and upper horizontal lines of the boxes represent the first quartile, median, and third quartile, respectively. Lower and upper whiskers represent 1.5*IQR + first quartile and 1.5*IQR + third quartile, respectively.

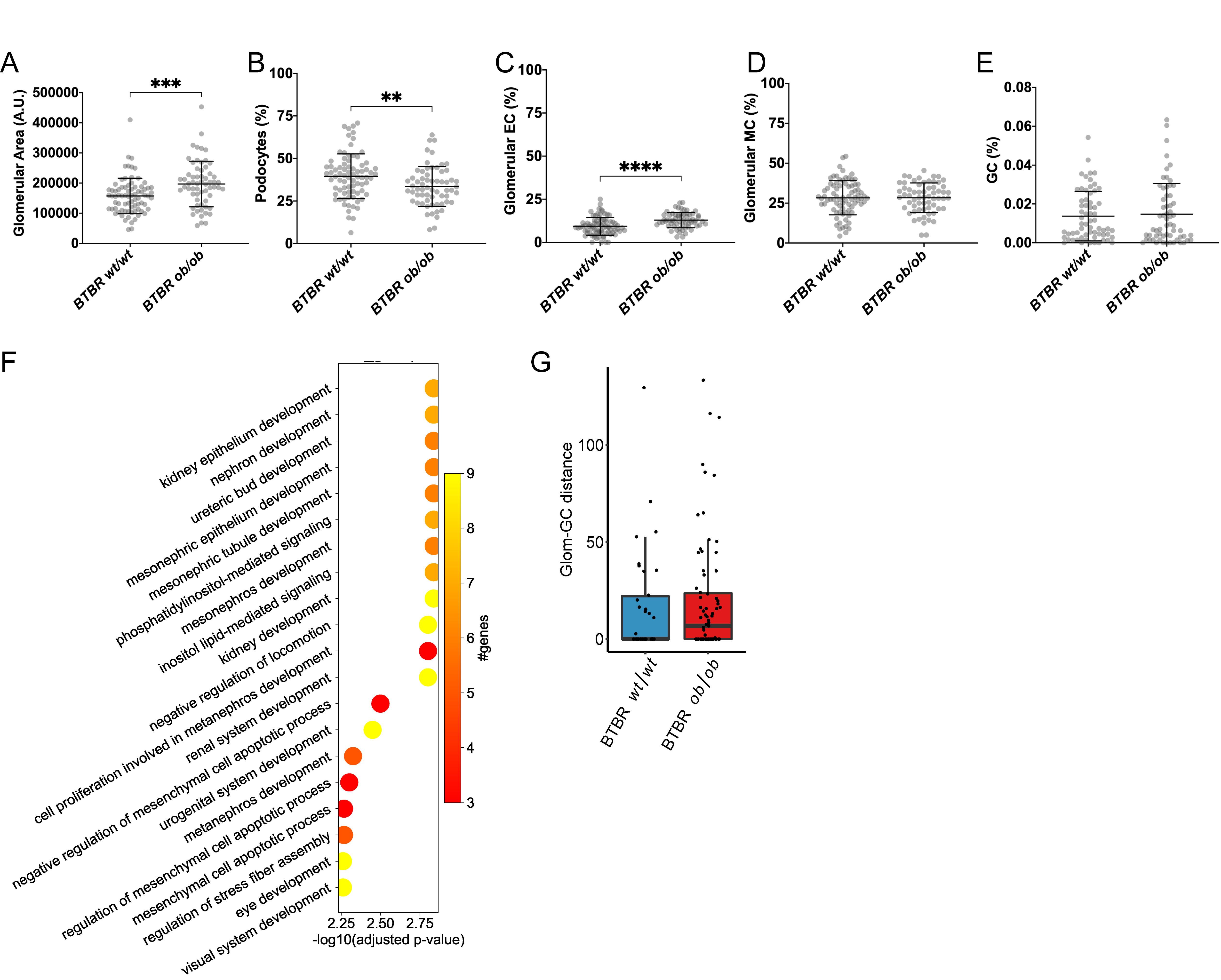

**Fig. S14. Spatial transcriptomics reveals injured podocytes and disorganized granular cells in kidney tissue from diabetic BTBR *ob/ob* mice.** Plots generated from HCR validation of (**A**) average area of glomeruli (p-value 0.0002), (**B**) percentage of *Nphs2*+ podocytes within the glomerulus (p-value 0.0034), (**C**) percentage of *Ehd3*+ glomerular endothelial cells (p-value <0.0001), (**D**) percentage of *Ctgf*+ glomerular mesangial cells, (**E**) percentage of *Ren1*+ granular cells immediately adjacent to glomeruli in BTBR *wt/wt* and BTBR *ob/ob* mice*.* Each data point is a structure from 4 mice per genotype. (**F**) Gene ontology plot showing top 20 statistically significant pathways enriched in differentially expressed genes from podocytes classified as diseased. Color scale shows the number of differentially expressed genes in diseased podocytes for each pathway. All error bars represent standard deviation of the mean. (**G**) Minimum distances between edges of glomerular and granular cell structures. Lower, middle, and upper horizontal lines of the boxes represent the first quartile, median, and third quartile, respectively. Lower and upper whiskers represent 1.5*IQR + third quartile, respectively.

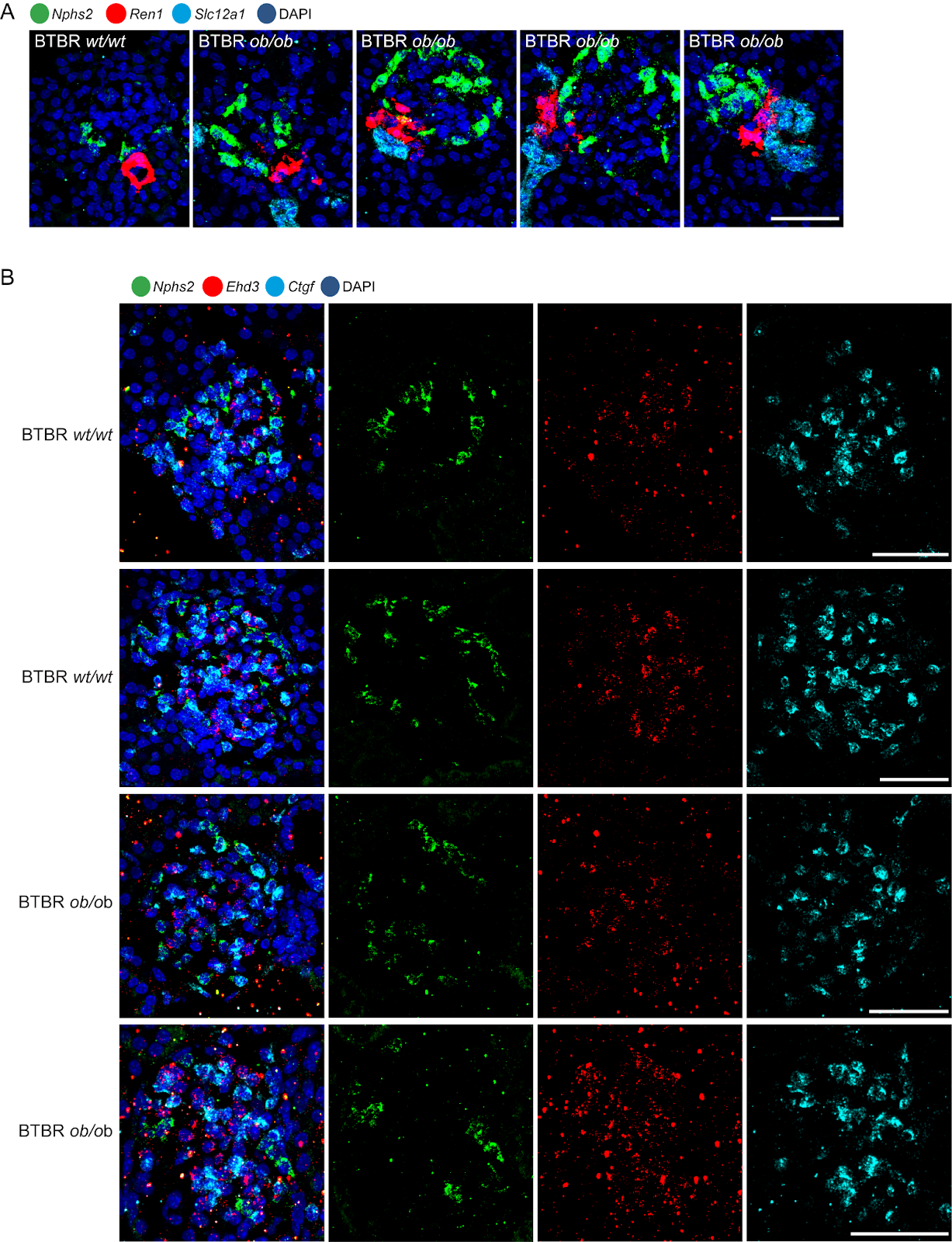

**Fig. S15. BTBR *ob/ob* kidneys have disorganized granular cells and injured podocytes.**

HCR images showing (**A**) *Nphs2*+ podocytes, *Ren1*+ granular cells, *Slc12a1*+ TAL, and all cells in DAPI. (**B**) *Nphs2*+ podocytes, Ehd3+ endothelial cells, *Ctgf*+ mesangial cells, *Nphs2*+ *Ctgf*+ double positive injured podocytes, and all cells in DAPI. Scale bar 50µm. Data obtained from cross sections of 4 mice per genotype.

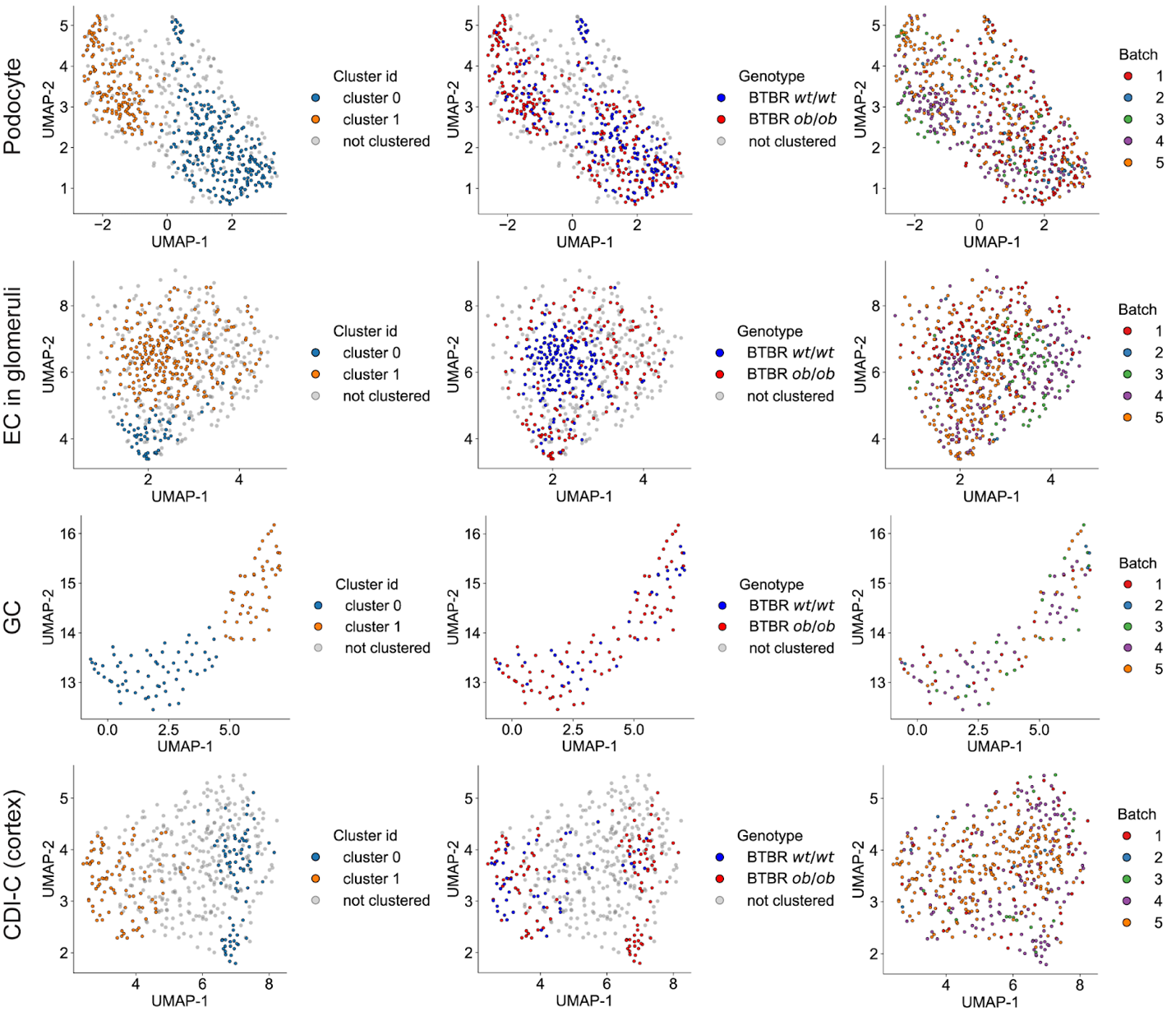

**Fig. S16. Clustering analysis of BTBR *wt/wt* and BTBR *ob/ob* podocytes, glomerular EC, GC, and CD-IC in the cortex.** UMAP projections of cluster labels (left), genotypes of arrays that data points were derived from (middle), and batch ids of arrays (right). Data points are averaged gene expression profiles of podocytes in glomeruli, EC in glomeruli, granular cell clusters, and CD-IC cortex partitions. Diseased clusters were defined by podocyte cluster 1 shows ~76% BTBR *ob/ob* and ~24% BTBR *wt/wt* glomeruli, EC cluster 0 shows 75% BTBR *ob/ob* and 25% BTBR *wt/wt* glomeruli, GC cluster 0 shows ~71% BTBR *ob/ob* and ~29% BTBR *wt/wt* structures, and CD-IC cluster 0 shows ~88% BTBR *ob/ob* and ~12% BTBR *wt/wt* structures.

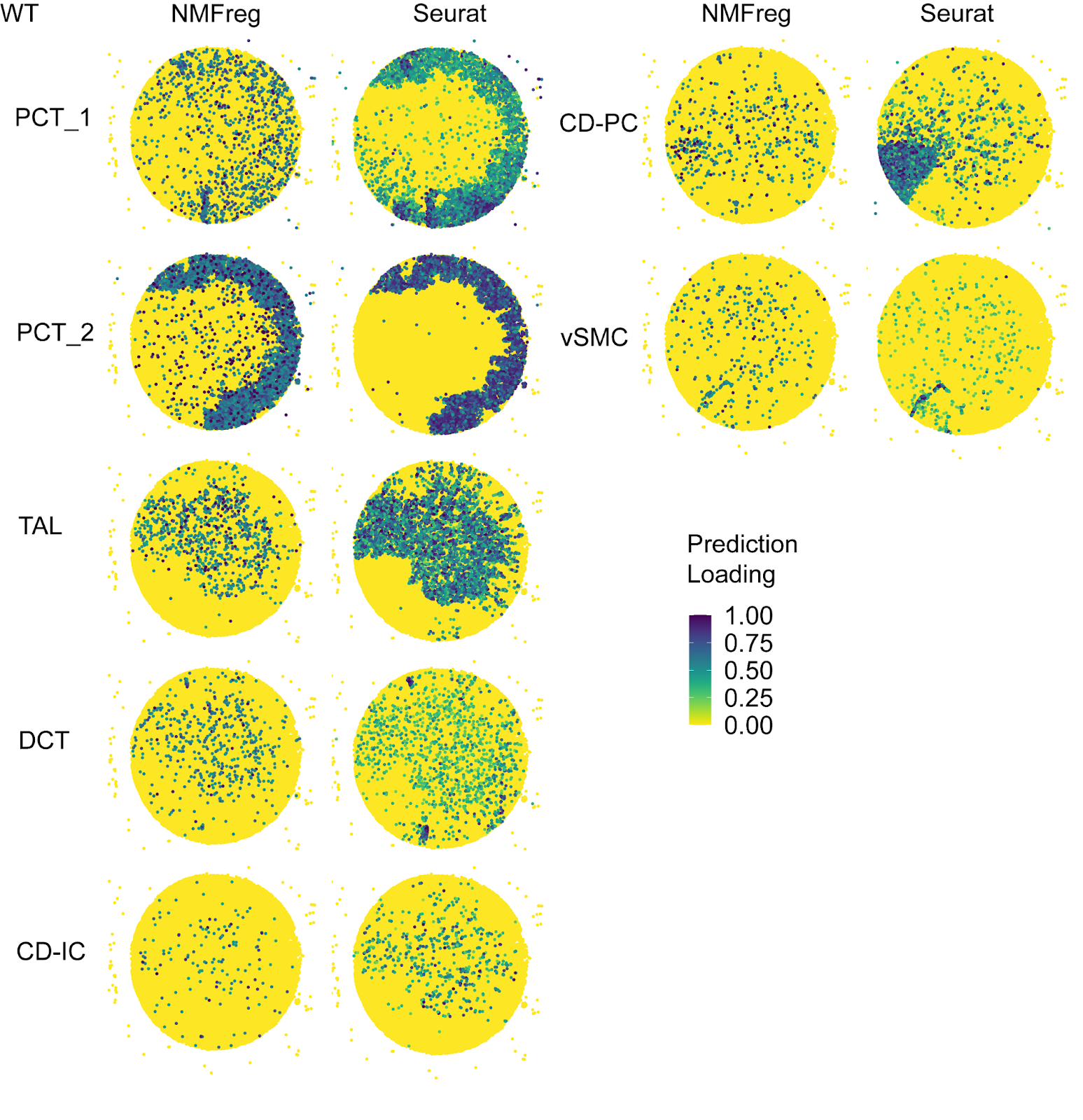

**Fig. S17. Comparison of NMFreg and Seurat cell mappings in WT controls of UMOD mice.** Spatial distribution of cell type prediction loadings from NMFreg (left) and Seurat transfer learning (right) methods.

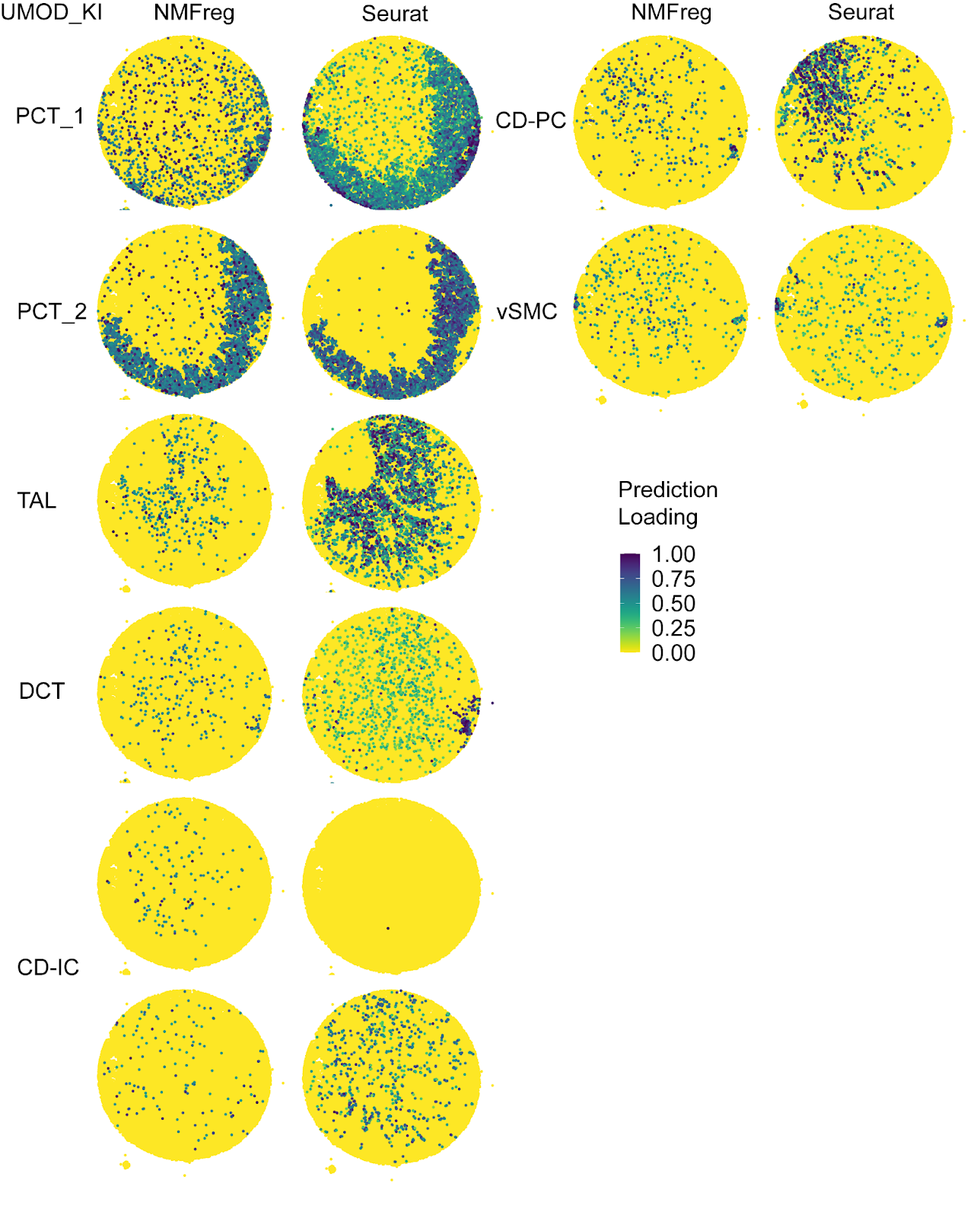

**Fig. S18. Comparison of NMFreg and Seurat cell mappings in homozygous UMOD-KI mice.** Spatial distribution of cell type prediction loadings from NMFreg (left) and Seurat transfer learning (right) methods.

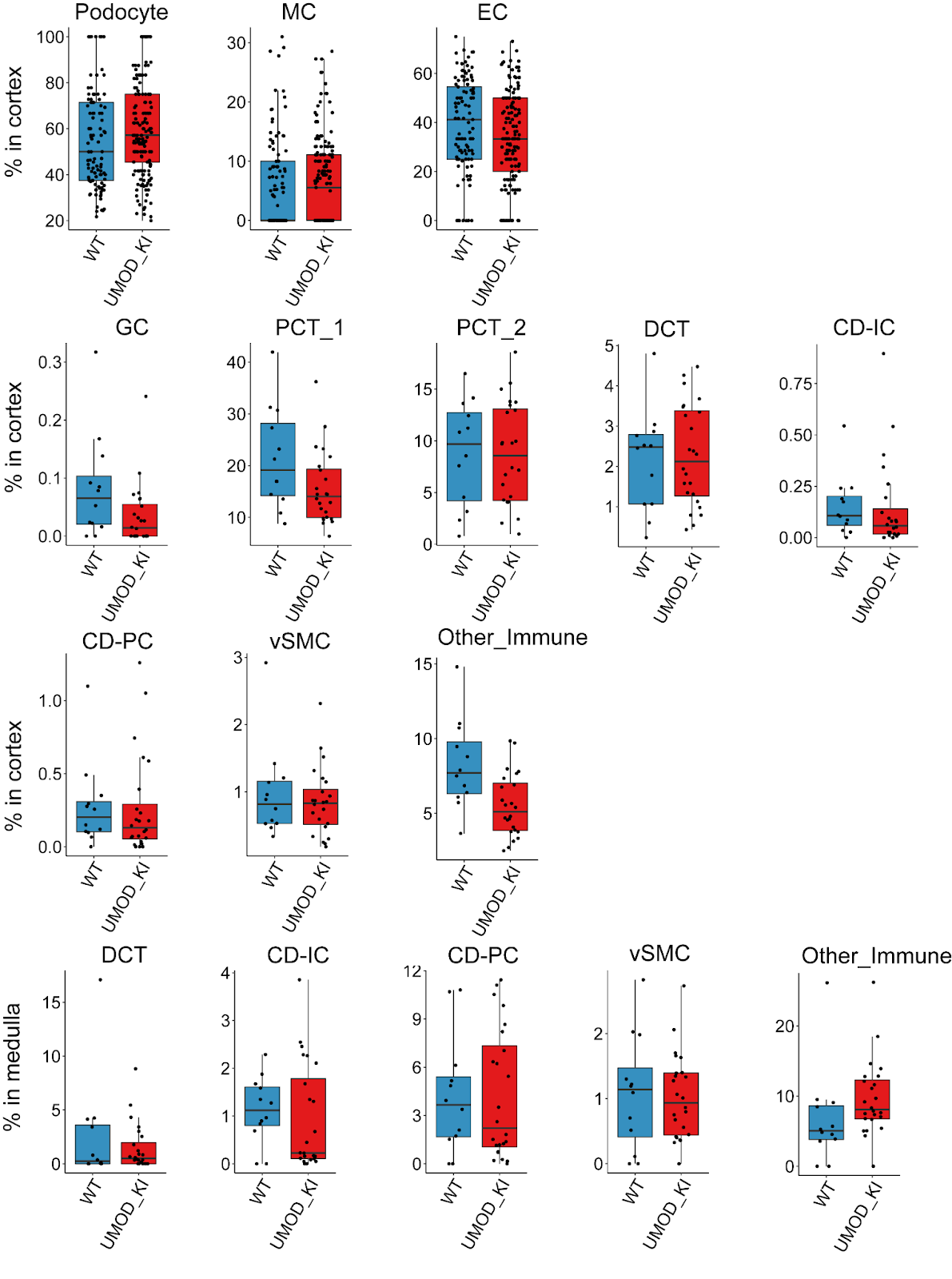

**Fig. S19. Cell type percentages across arrays in WT and UMOD-KI mice in cortex and medulla.** Percentage of glomerular cells classified as podocytes (top left), MC (top middle), and EC (top right), and percentage of cortex (middle) and medulla (bottom) beads of arrays

classified non-glomerular cell types. Lower, middle, and upper horizontal lines of the boxes represent the first quartile, median, and third quartile, respectively. Lower and upper whiskers represent 1.5*IQR + first quartile and 1.5*IQR + third quartile, respectively.

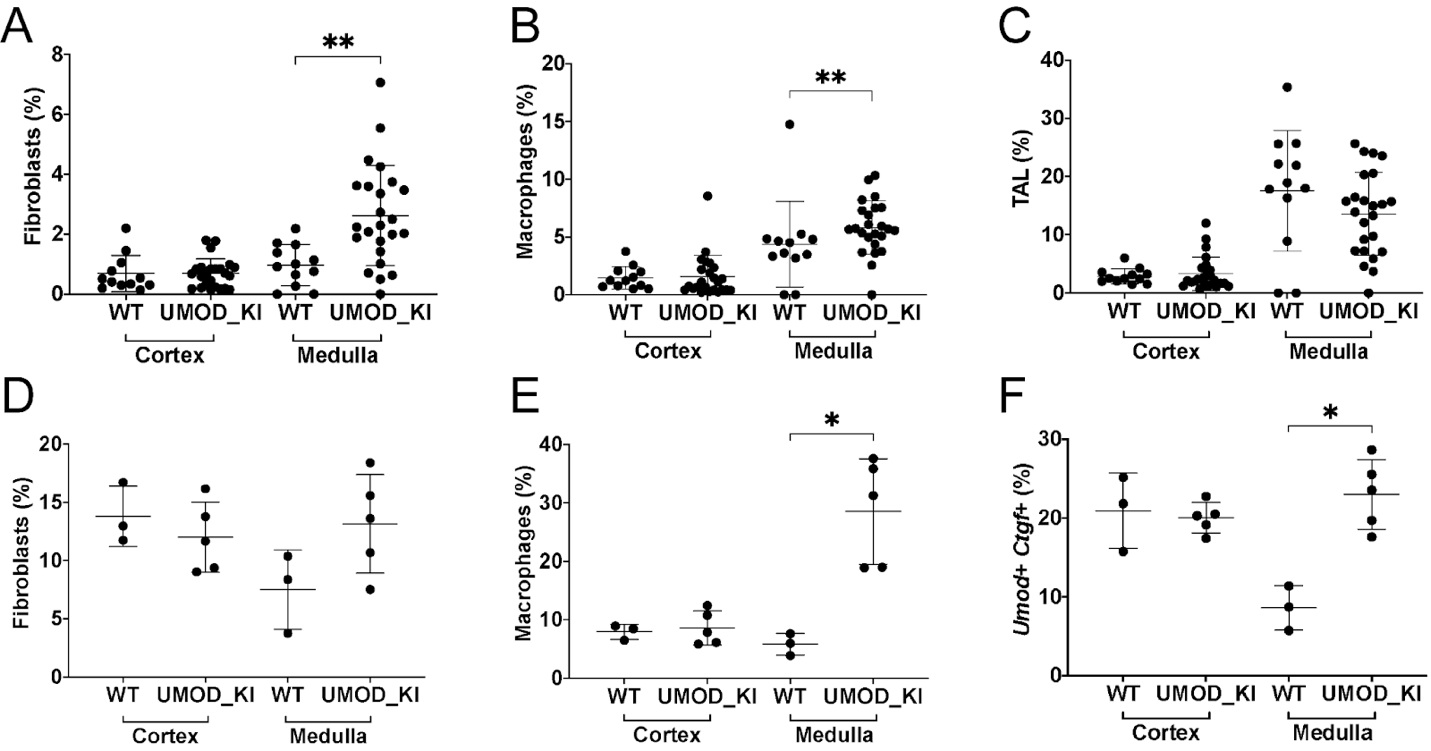

**Fig. S20. Spatial transcriptomics reveals that a monogenic mutation in *Umod* causes medulla-specific fibroblast and macrophage expansion, and promotes medullary TAL injured.** (**A**) fibroblasts (p-value 0.0011), (**B**) macrophages (p-value 0.0079), (**C**) TAL in cortex and medulla. Each point represents an array from 3 WT and 4 UMOD-KI mice. Plots generated from HCR validation showing percentage of cells that are (**D**) *Ctgf*+ fibroblasts, (**E**) *C1qb*+ macrophages (p-value 0.0357), (**F**) *Umod Ctgf* double positive injured TAL (p-value 0.0357). Each point represents a mouse. All error bars represent standard deviation of the mean.

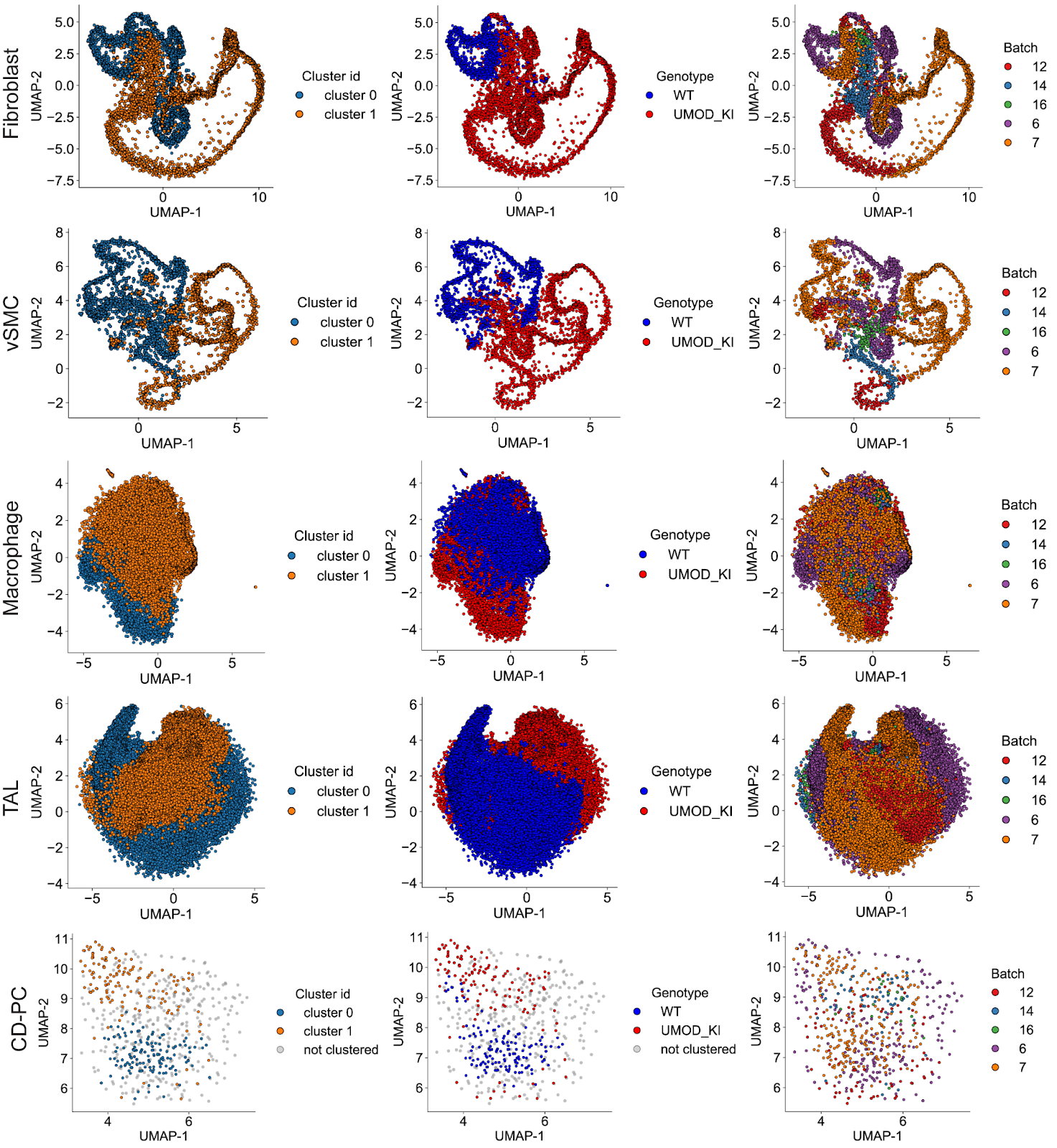

**Fig. S21. Clustering analysis of WT and UMOD-KI fibroblasts, vSMC, macrophages, TAL, and CD-PC in the medulla.** UMAP projections of cluster labels (left), genotypes of arrays that data points were derived from (middle), and batch ids of arrays (right). Data points are individual fibroblast, vSMC, macrophage, and TAL beads in the medulla, and averaged gene expression profiles of CD-PC in the medulla. Diseased clusters were defined by fibroblast cluster 1 shows ~99.5% UMOD_KI and ~0.5% WT beads, vSMC cluster 1 shows 100% UMOD_KI and 0% WT beads, macrophage cluster 0 shows ~99.7% UMOD_KI and ~0.3% WT beads, TAL cluster 1 shows ~100% UMOD_KI and ~0% WT beads, and CD-PC cluster 1 shows ~91% UMOD_KI and ~9% WT structures.

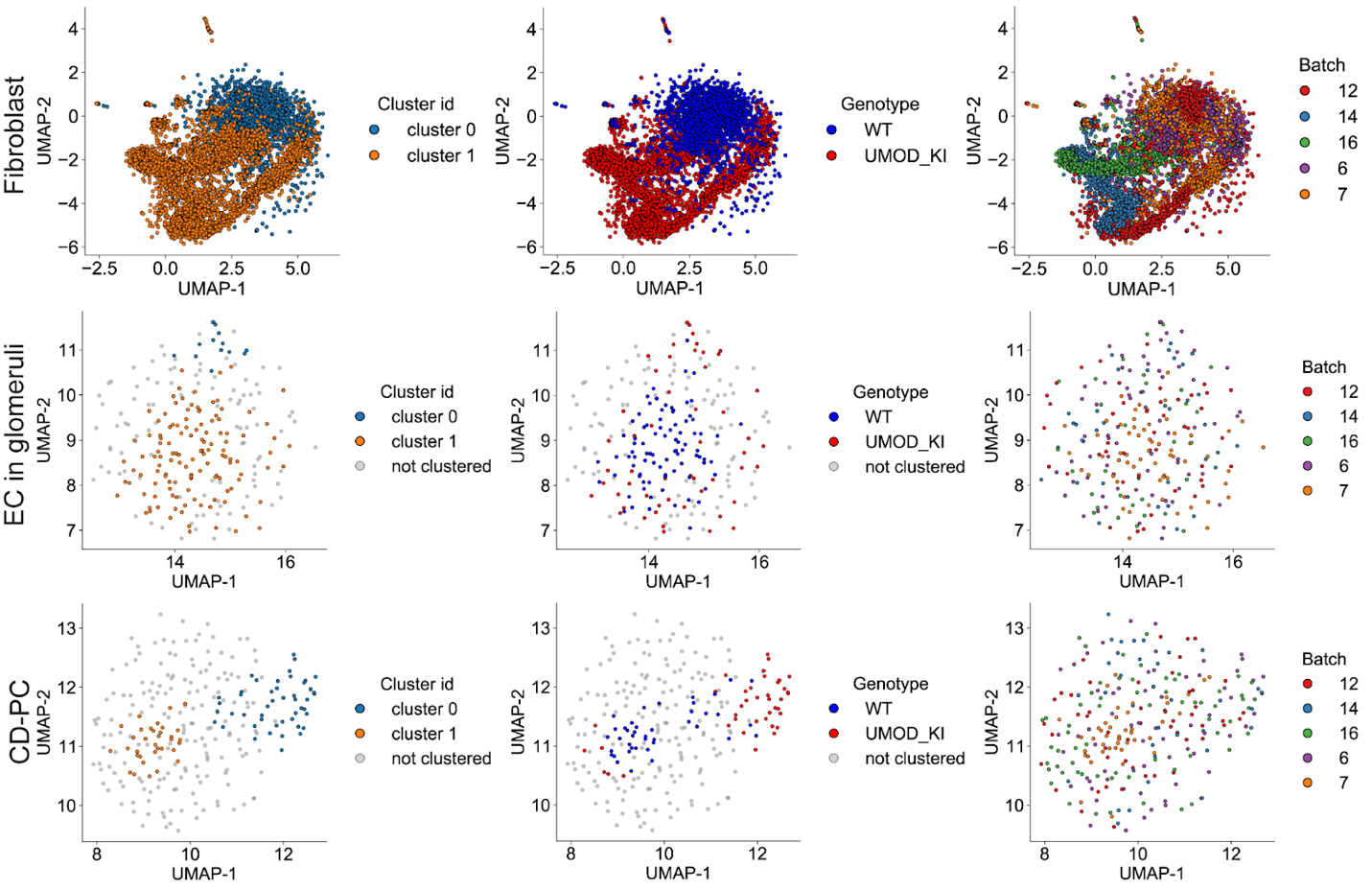

**Fig. S22. Clustering analysis of WT and UMOD-KI fibroblasts, glomerular EC, and CD-PC in the cortex.** UMAP projections of cluster labels (left), genotypes of arrays that data points were derived from (middle), and batch ids of arrays (right). Data points are individual fibroblast beads in the cortex, and averaged gene expression profiles of EC in glomeruli and CD-PC partitions in the cortex. Diseased clusters were defined by fibroblast cluster 1 shows ~95% UMOD_KI and ~5% WT beads, EC cluster 0 shows ~85% UMOD_KI and ~15% WT glomeruli, and CD-PC cluster 0 shows ~72% UMOD_KI and ~28% WT structures.

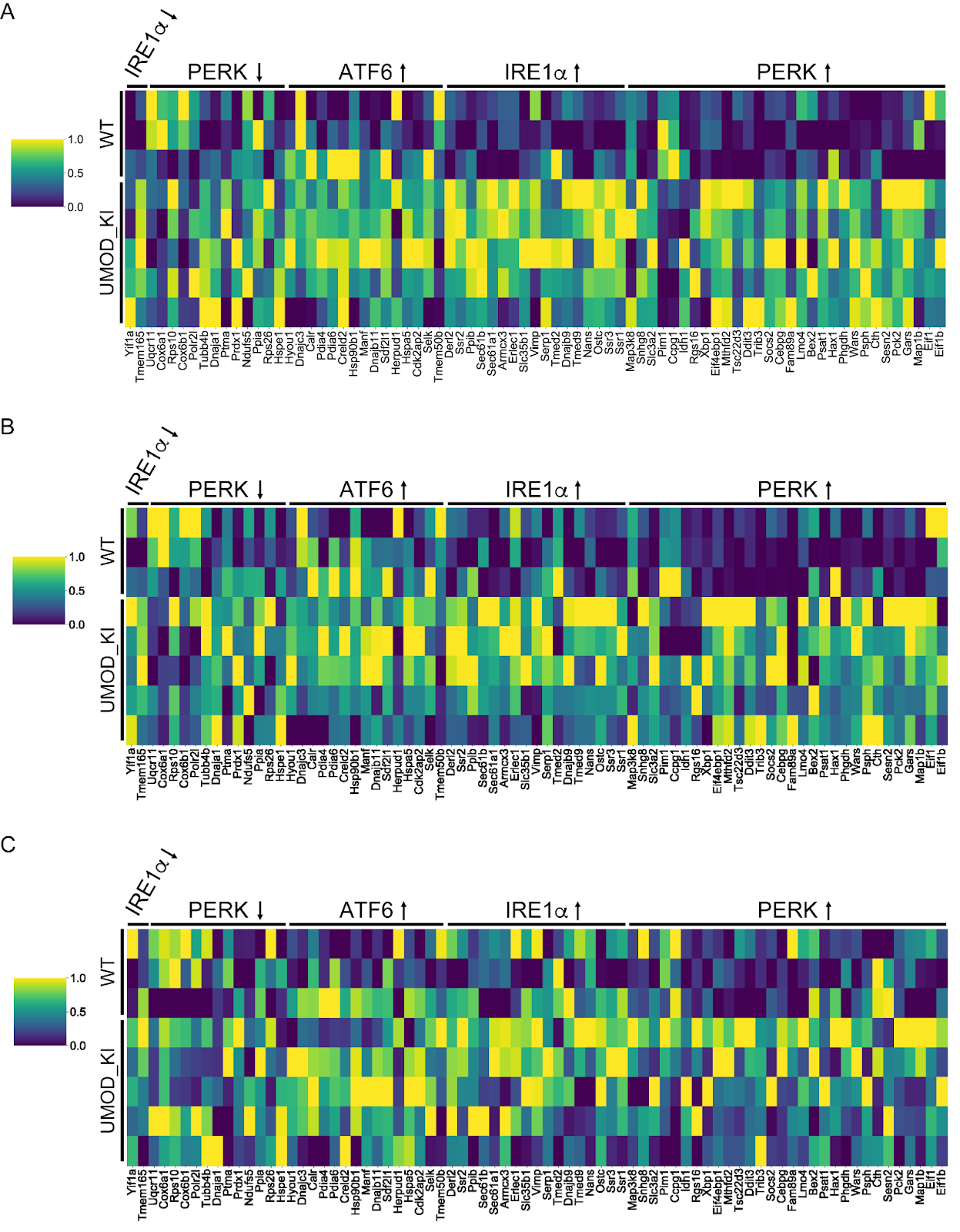

**Fig. S23. UPR unchanged in cortex TAL and PCT.** Heatmaps showing relative expression level of UPR genes averaged across arrays in WT and UMOD-KI mice in (**A**) all, cortical and medullary, TAL, (**B**) cortical TAL, and (**C**) cortical PCT_1 (cortex beads). No significant genes were identified in these cell types (Benjamani-Hochberg-corrected p<0.05). This is a negative control for the specific upregulation of UPR gene signatures in medullary TALs.

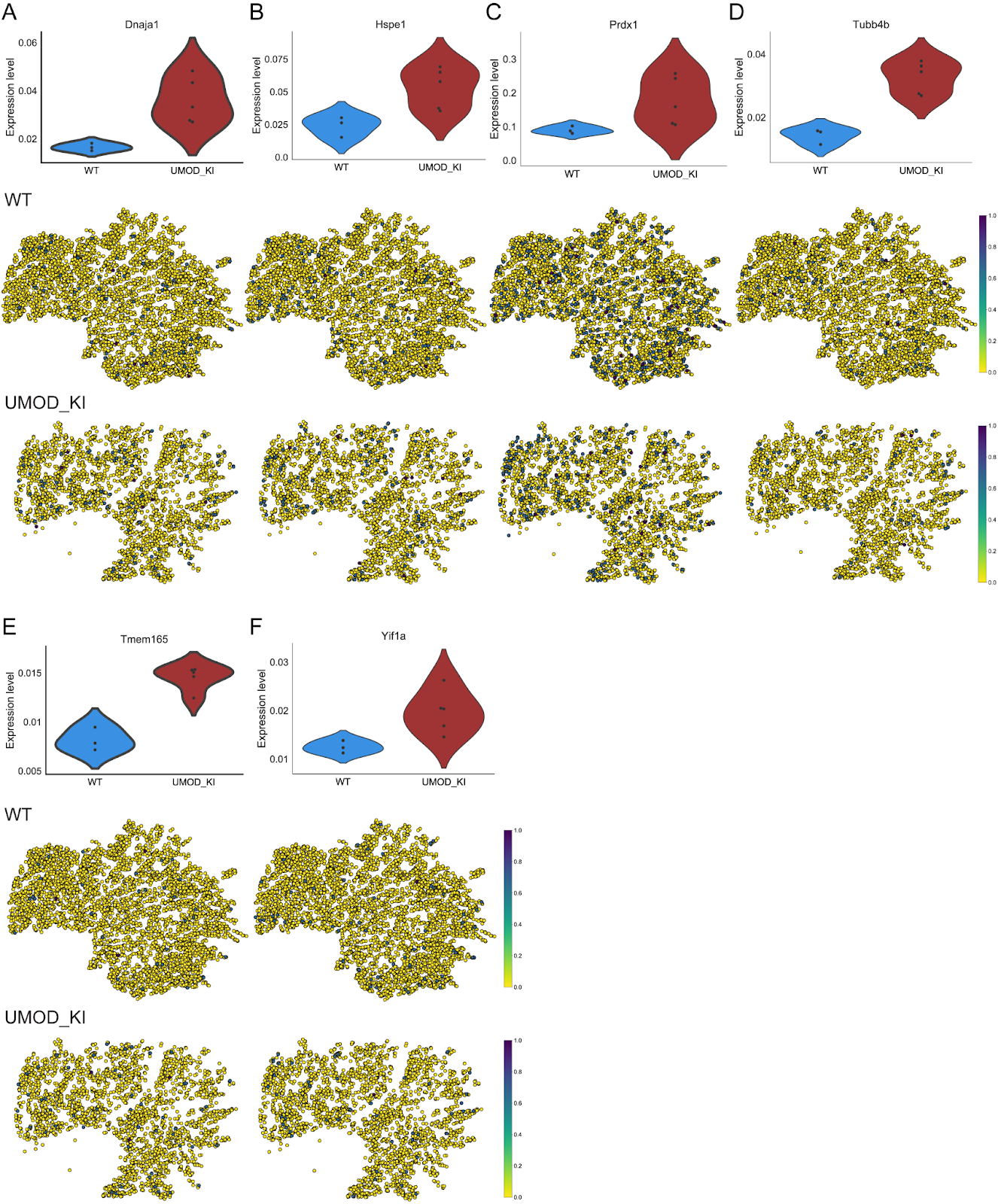

**Fig. S24. PERK and IRE1α downregulated UPR genes significantly changed in medulla TAL.** Violin plots (top) showing expression level in medulla TAL and all medulla TAL beads in arrays (bottom) are shown with expression level for significant genes in PERK downregulated pathway (**A**) *Dnaja1*, (**B**) *Hspe1*, (**C**) *Prdx1*, and (**D**) *Tubb4b* and IRE1ɑ downregulated pathway (**E**) *Tmem165,* and (**F**) *Yif1a*.

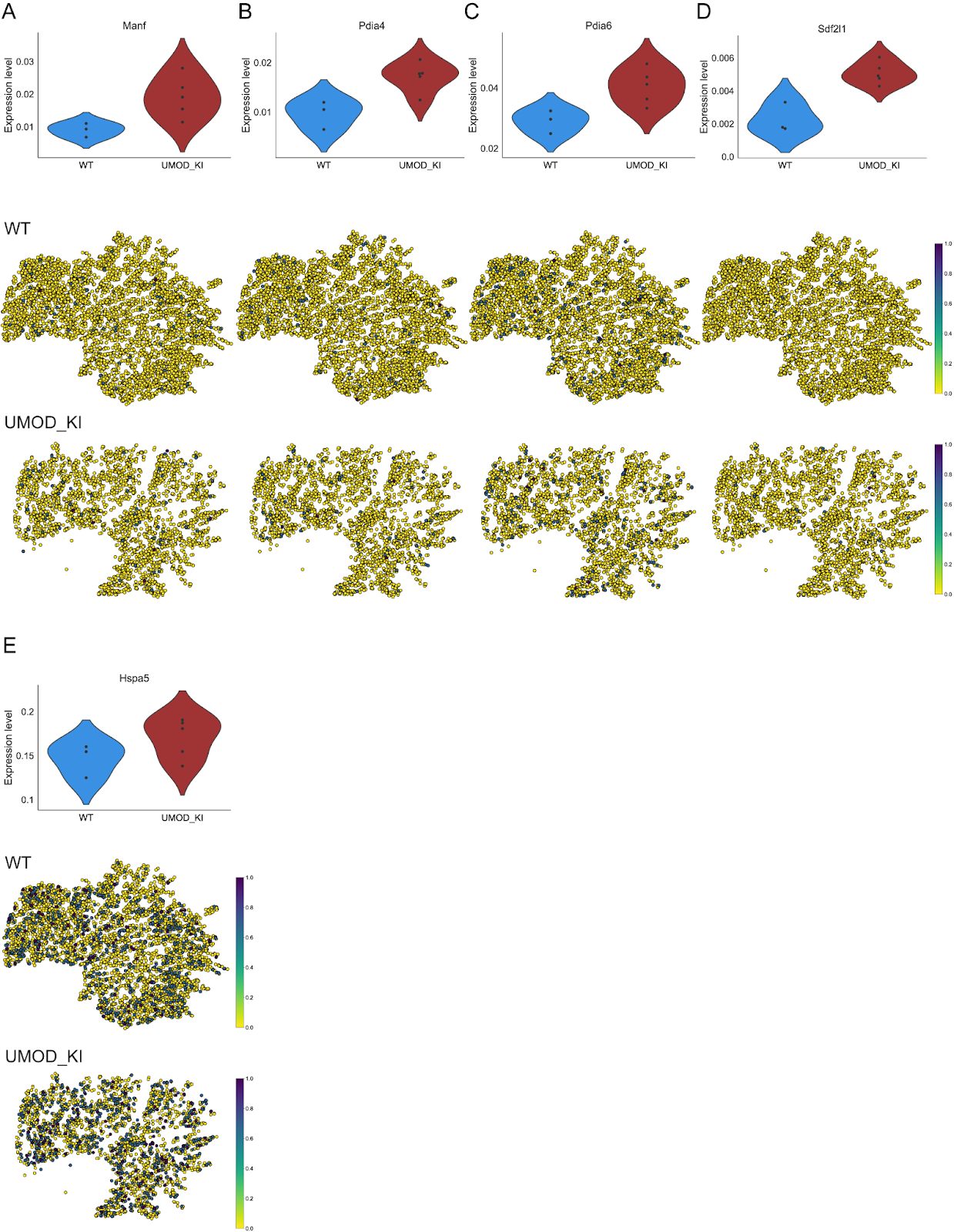

**Fig. S25. ATF6 upregulated UPR genes significantly changed in medulla TAL.** Violin plots (top) showing expression level in medulla TAL and all medulla TAL beads in arrays (bottom) are shown with expression level for significant genes in ATF6 upregulated pathway (**A**) *Manf*, (**B**) *Pdia4*, (**C**) *Pdia6,* (**D**) *Sdf2l1*, and (**E**) *Hspa5*.

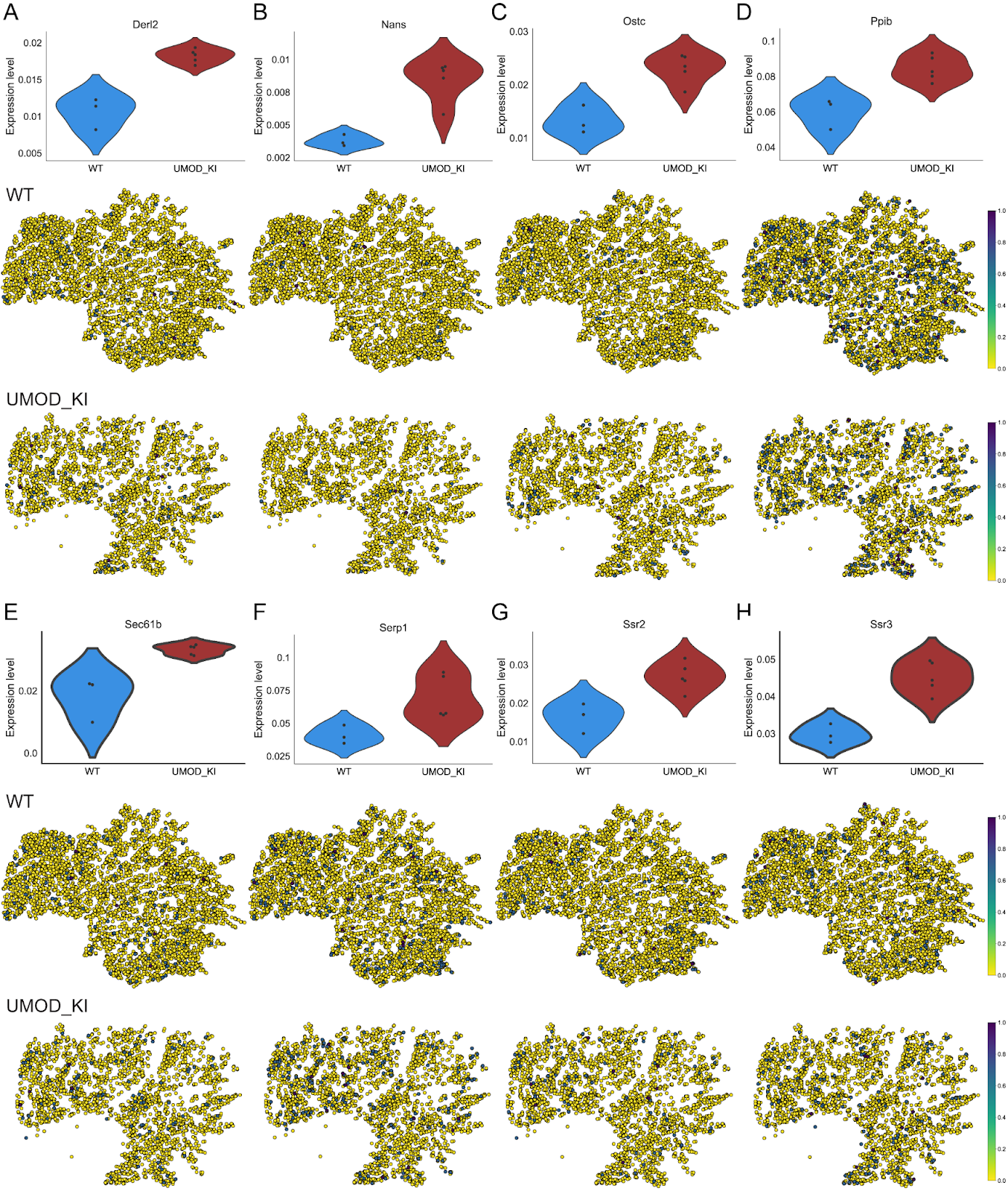

**Fig. S26. IRE1α upregulated UPR genes significantly changed in medulla TAL.** Violin plots (top) showing expression level in medulla TAL and all medulla TAL beads in arrays (bottom) are shown with expression level for significant genes in IRE1ɑ upregulated pathway (**A**) *Derl2,* (**B**) *Nans*, (**C**) *Ostc,* (**D**) *Ppib*, (**E**) *Sec61b*, (**F**) *Serp1*, (**G**) *Ssr2,* and (**H**) *Ssr3*.

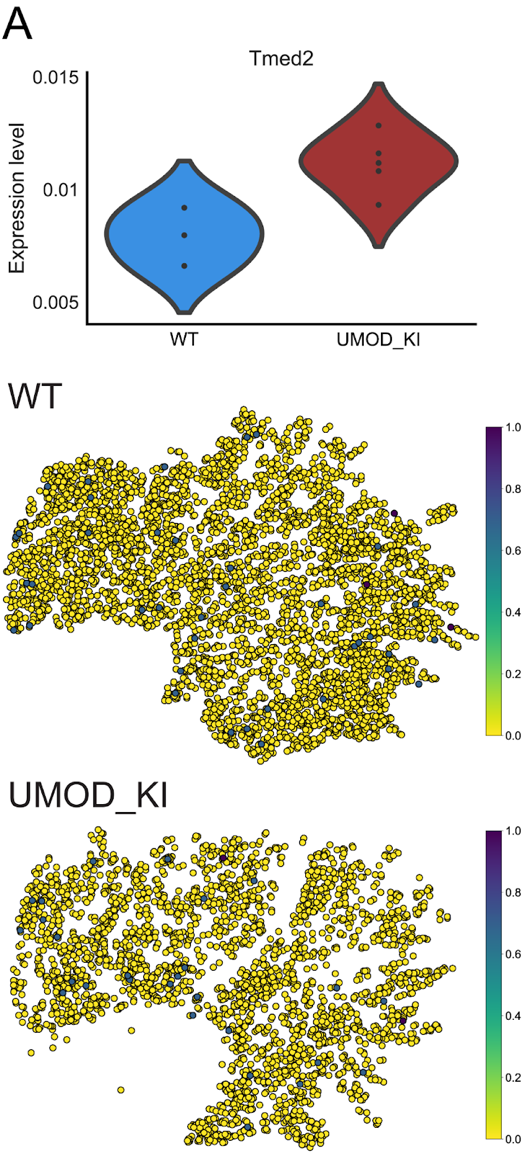

**Fig. S27. IRE1α upregulated UPR genes significantly changed in medulla TAL.** Violin plots (top) showing expression level in medulla TAL and all medulla TAL beads in arrays (bottom) are shown with expression level for significant genes in IRE1ɑ upregulated pathway (**A**) *Tmed2*.

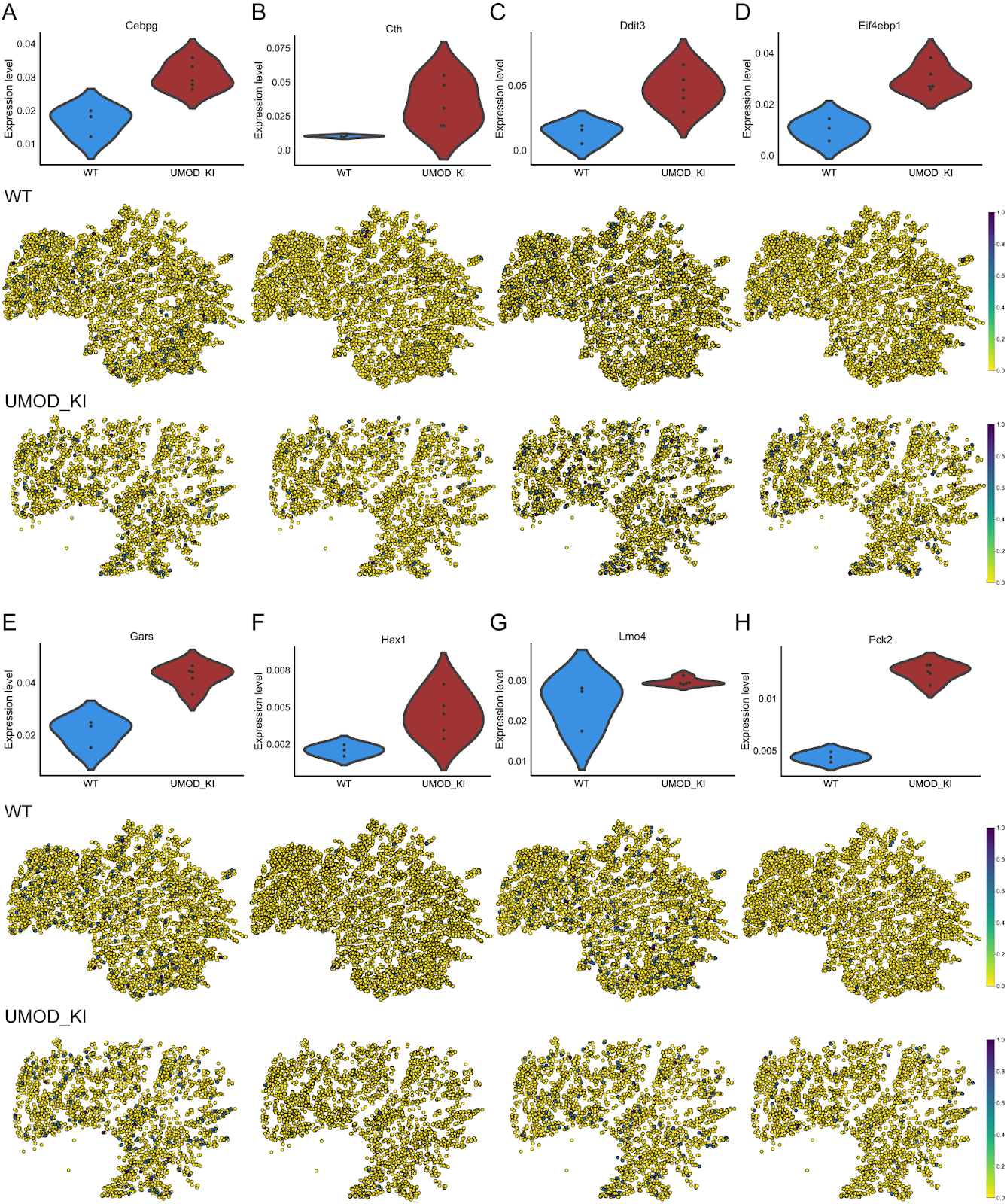

**Fig. S28. PERK upregulated UPR genes significantly changed  in medulla TAL.** Violin plots (top) showing expression level in medulla TAL and all medulla TAL beads in arrays (bottom) are shown with expression level for significant genes in PERK upregulated pathway (**A**) *Cebpg*, (**B**) *Cth*, (**C**) *Ddit3*, (**D**) *Eif4ebp1,* (**E**) *Gars*, (**F**) *Hax1*, (**G**) *Lmo4*, and (**H**) *Pck2*.

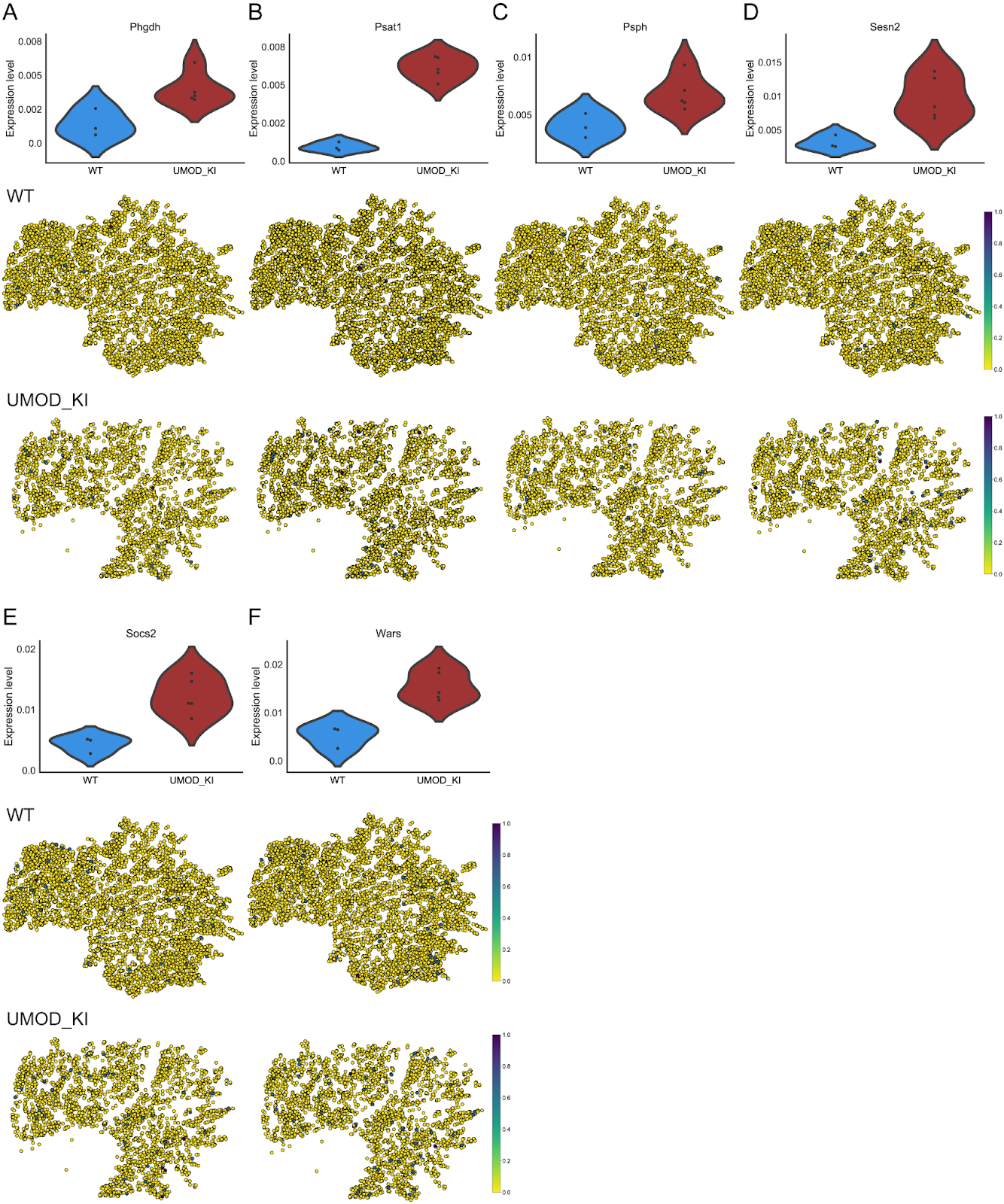

**Fig. S29. PERK upregulated UPR genes significantly changed  in medulla TAL.** Violin plots (top) showing expression level in medulla TAL and all medulla TAL beads in arrays (bottom) are shown with expression level for significant genes in PERK upregulated pathway (**A**) *Phgdh*, (**B**) *Psat1*, (**C**) *Psph*, (**D**) *Sesn2,* (**E**) *Socs2*, and (**F**) *Wars*.

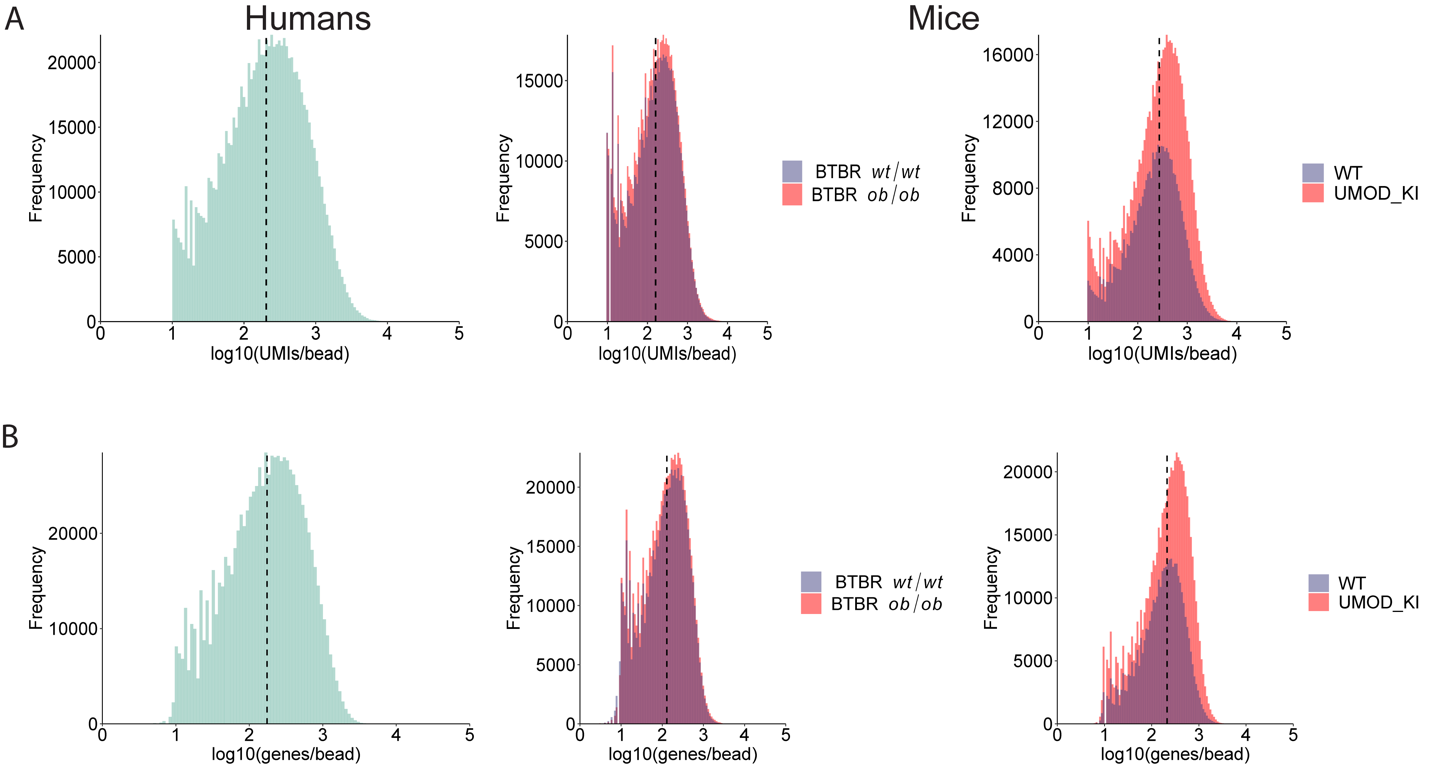

**Fig. S30. Frequency of UMIs per bead and genes per bead from humans and mice.** Histograms of A) log10 UMIs per bead and B) log10 genes per bead for human (left) and mouse (right) arrays. Dashed line represents the median value.

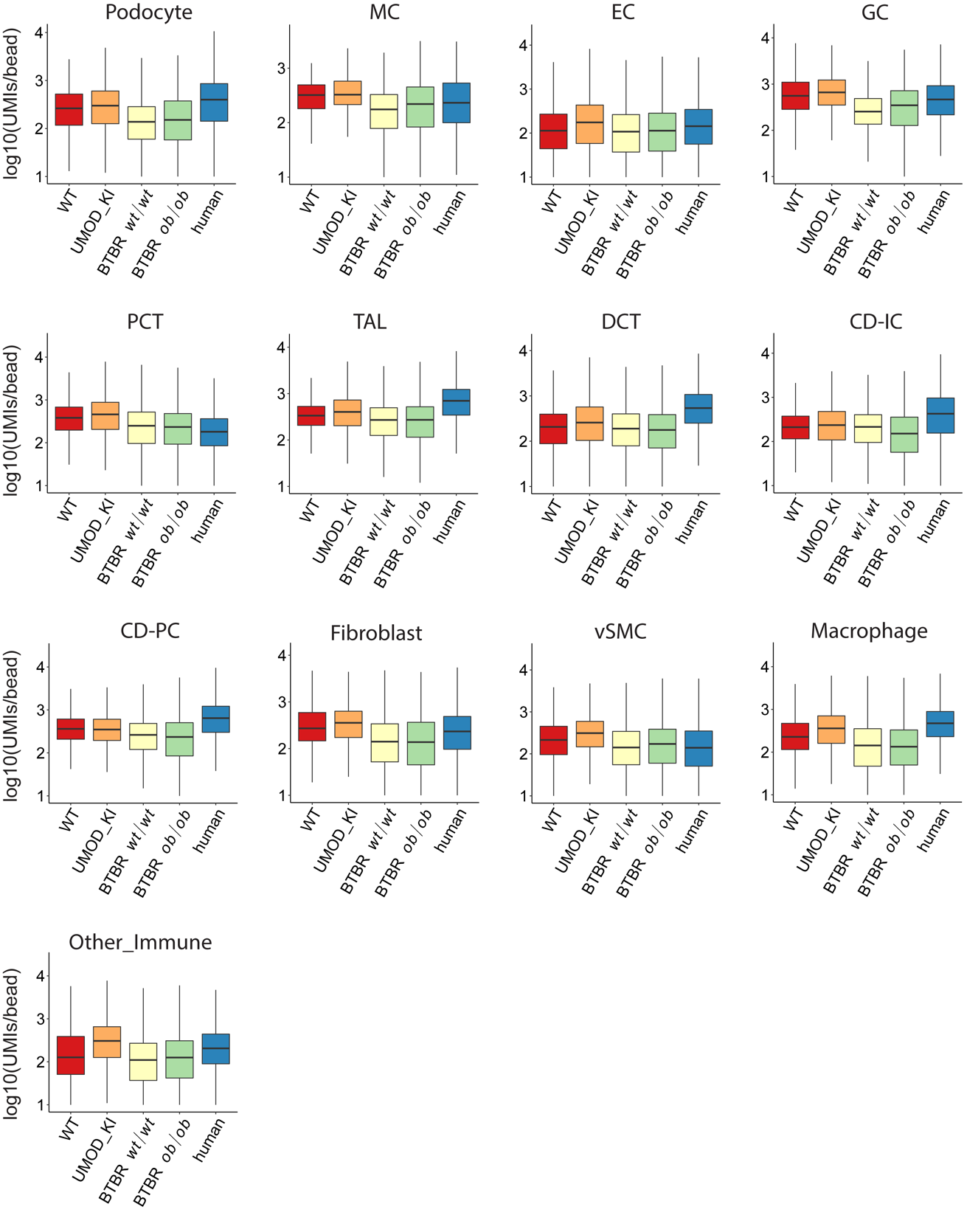

**Fig. S31. UMIs per bead plotted for humans and mice across all cell types.** Box plots of log10(UMIs/bead) for all beads classified as each cell type, grouped by genotype. Lower, middle, and upper horizontal lines of the boxes represent the first quartile, median, and third quartile, respectively. Lower and upper whiskers represent 1.5*IQR + first quartile and 1.5*IQR + third quartile, respectively.

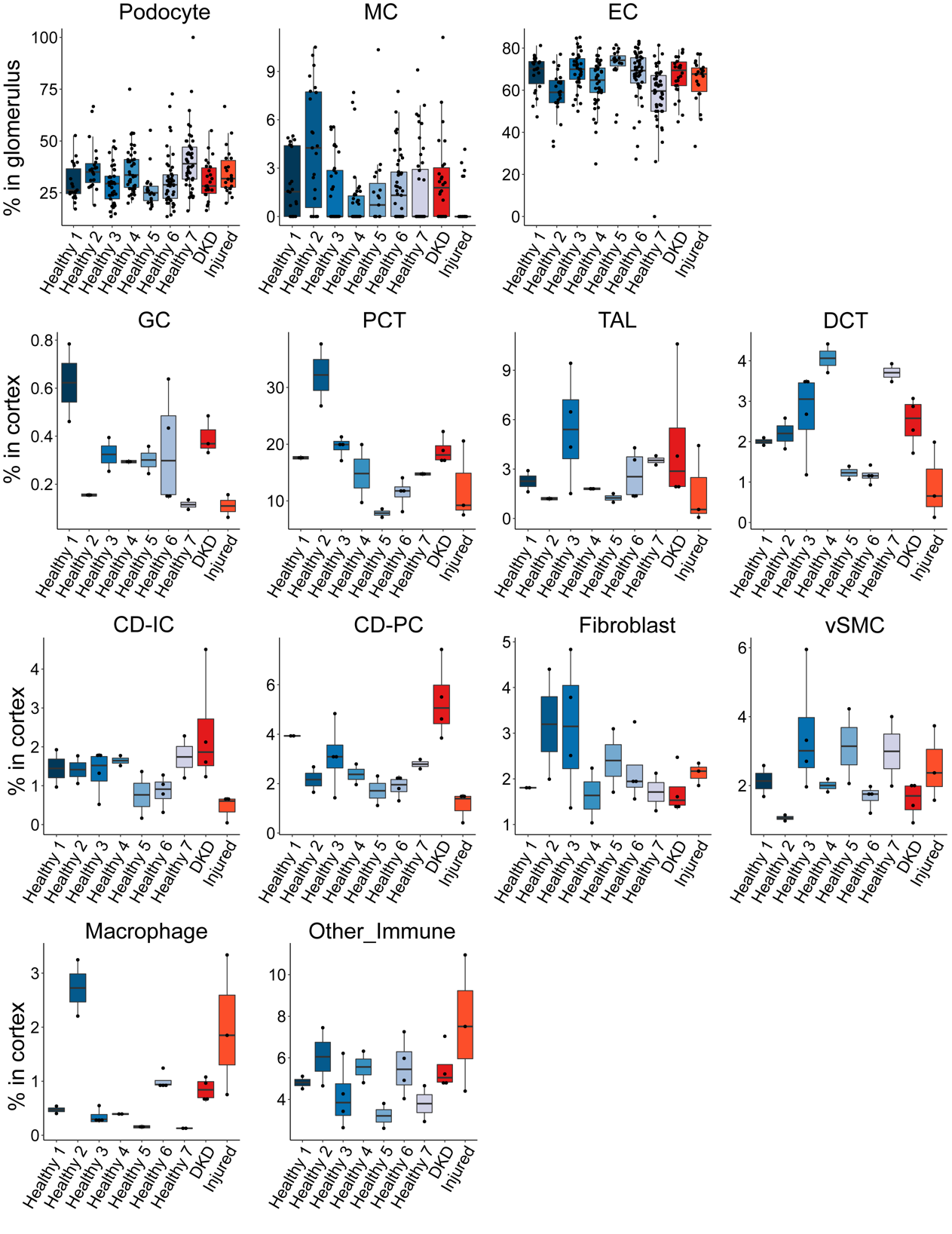

**Fig. S32. Cell type percentages from cortex regions across human arrays.** Percentage of glomerular cells classified as podocytes (top left), MC (top middle), and EC (top right), and percentage of cortex region of human arrays classified as all remaining cell types (bottom). Lower, middle, and upper horizontal lines of the boxes represent the first quartile, median, and third quartile, respectively. Lower and upper whiskers represent 1.5*IQR + first quartile and 1.5*IQR + third quartile, respectively.

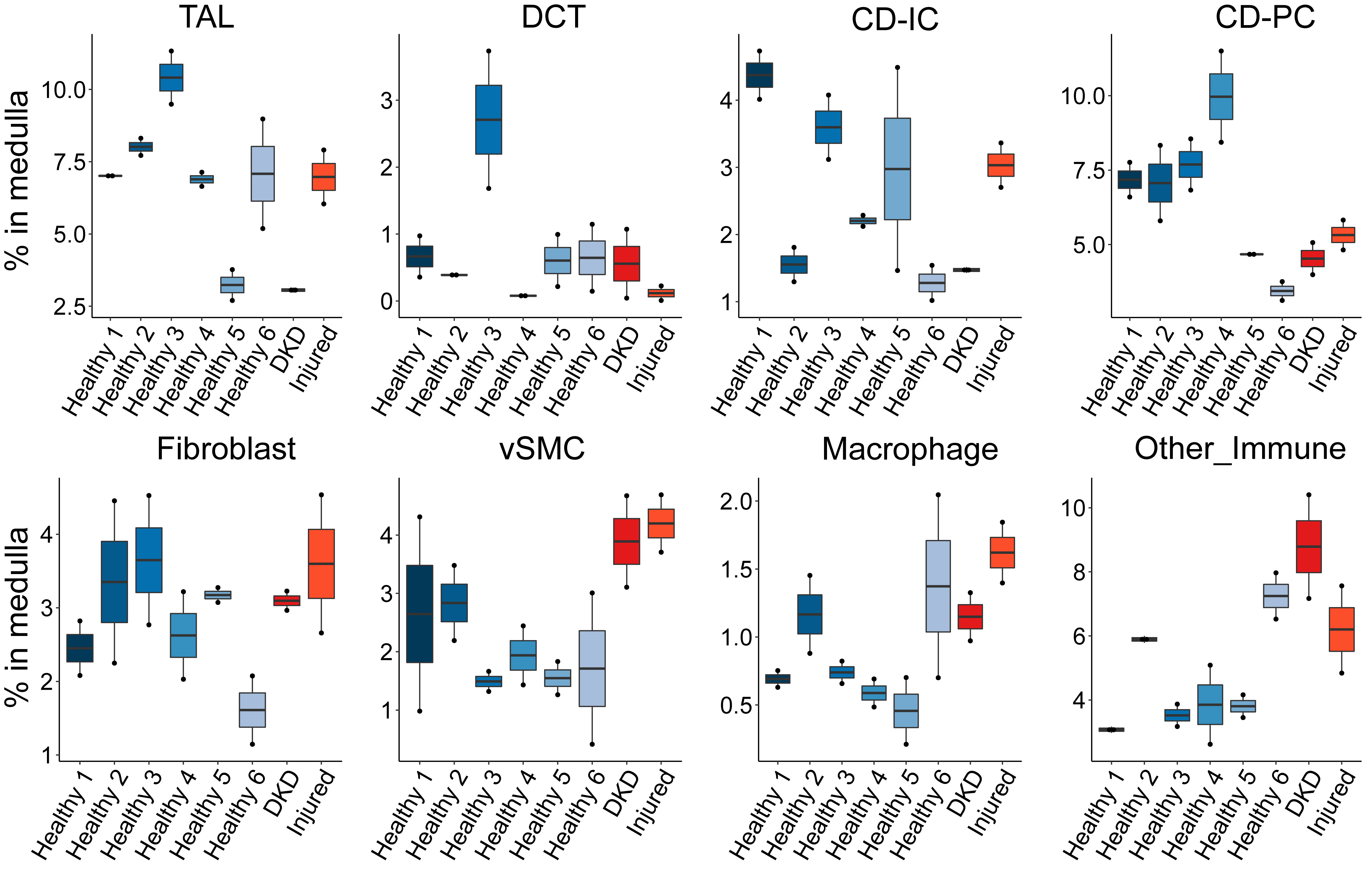

**Fig. S33. Cell type percentages from medulla regions across human arrays.** Percentage of medulla region of human arrays classified as all non-glomerular cell types. Lower, middle, and upper horizontal lines of the boxes represent the first quartile, median, and third quartile, respectively. Lower and upper whiskers represent 1.5*IQR + first quartile and 1.5*IQR + third quartile, respectively.

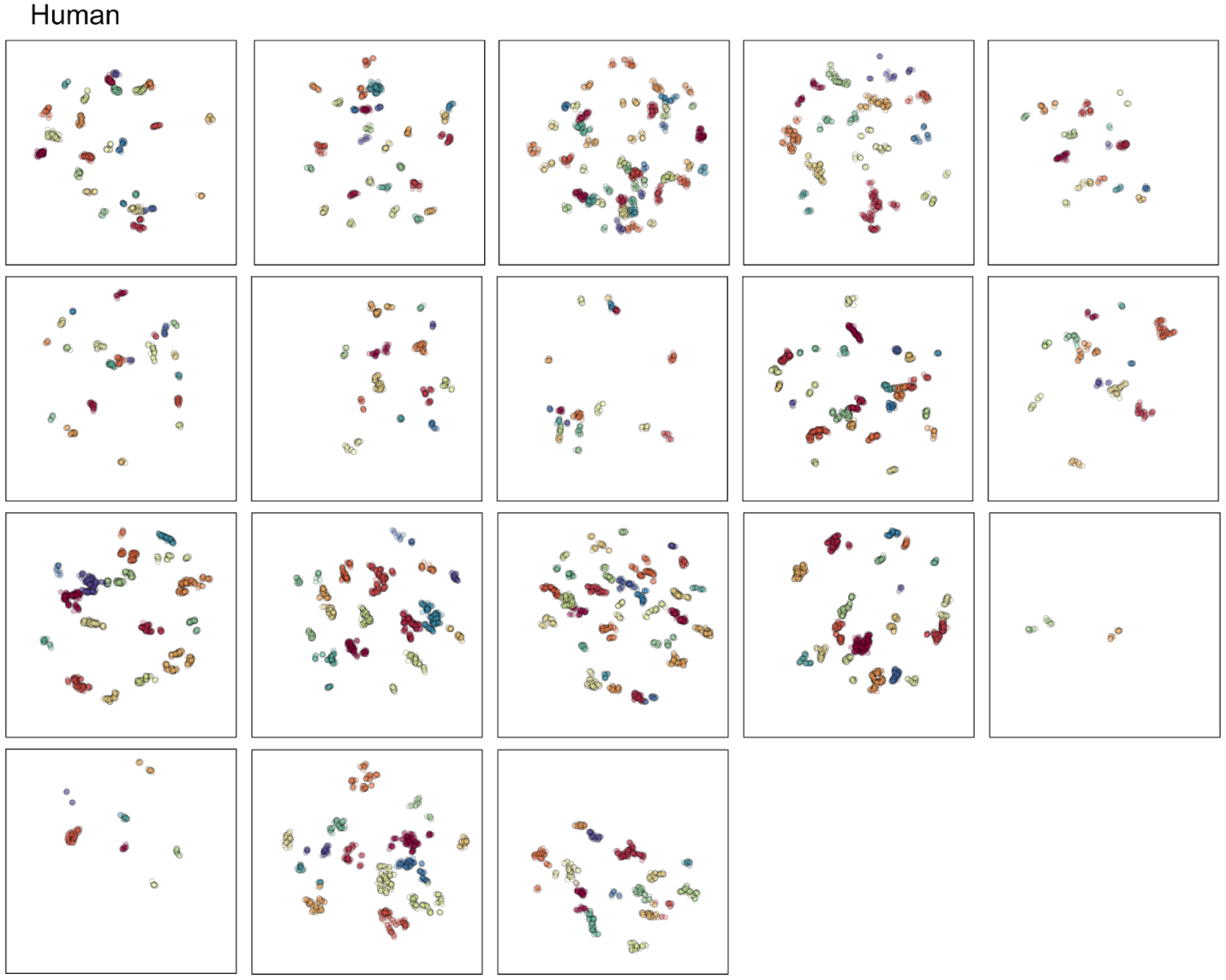

**Fig. S34. Human DCT tubule partitions in the cortex.** Spatial distribution of KMeans-derived partitions of DCT bead coordinates in the cortex, represented by unique colors across human arrays.

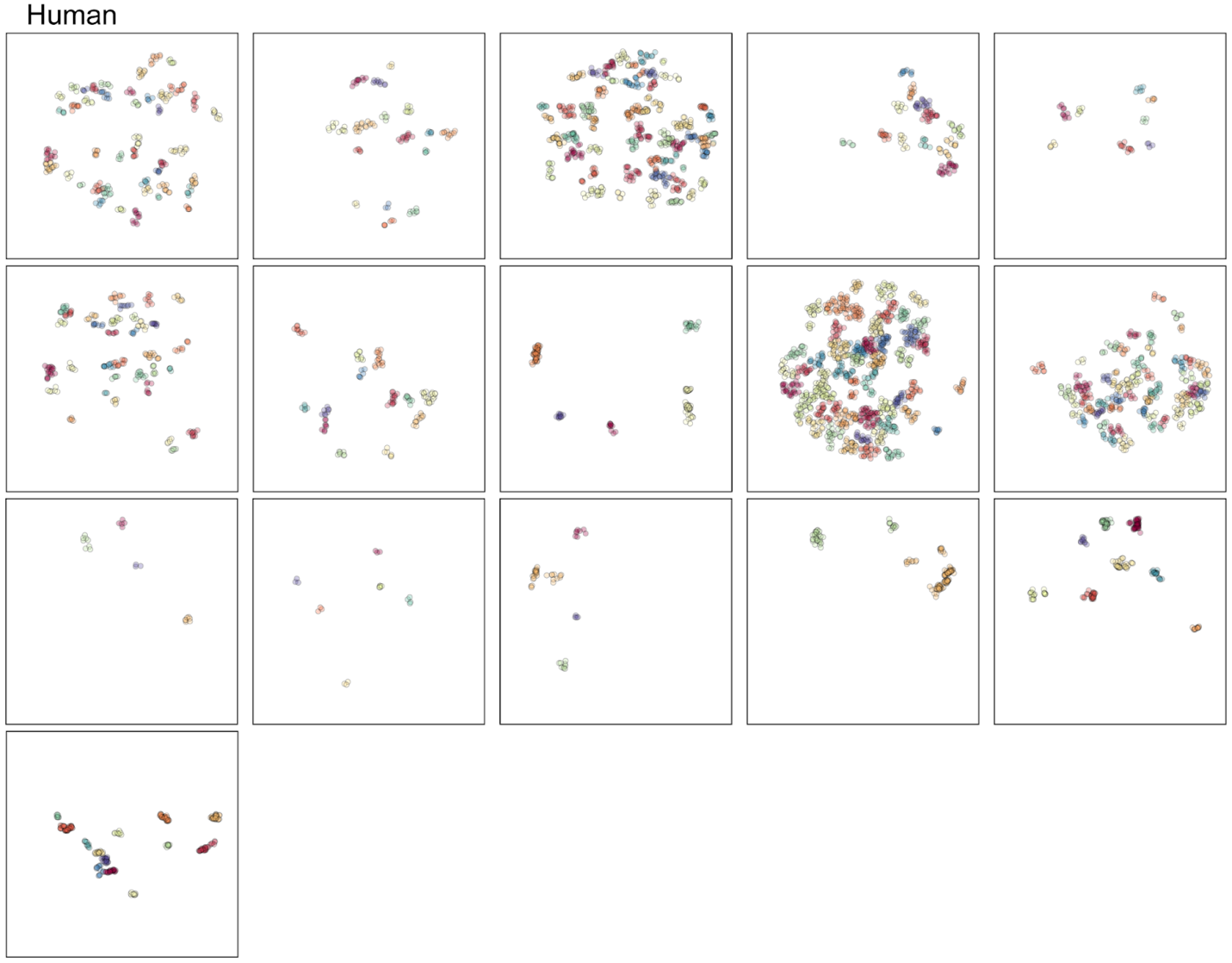

**Fig. S35. Human DCT tubule partitions in the medulla.** Spatial distribution of KMeans-derived partitions of DCT bead coordinates in the medulla, represented by unique colors across human arrays.

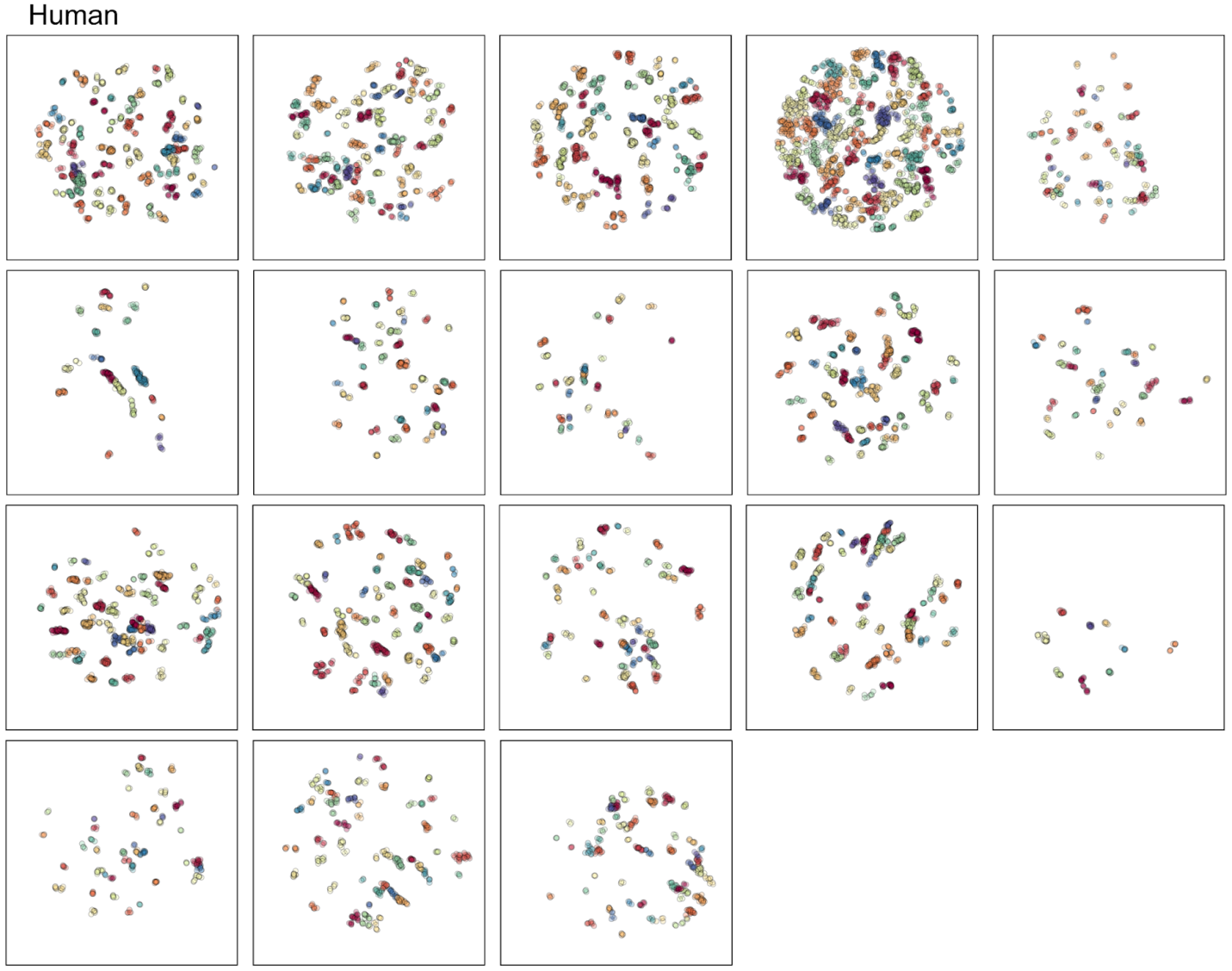

**Fig. S36. Human CD-PC tubule partitions in the cortex.** Spatial distribution of KMeans-derived partitions of CD-PC bead coordinates in the cortex, represented by unique colors across human arrays.

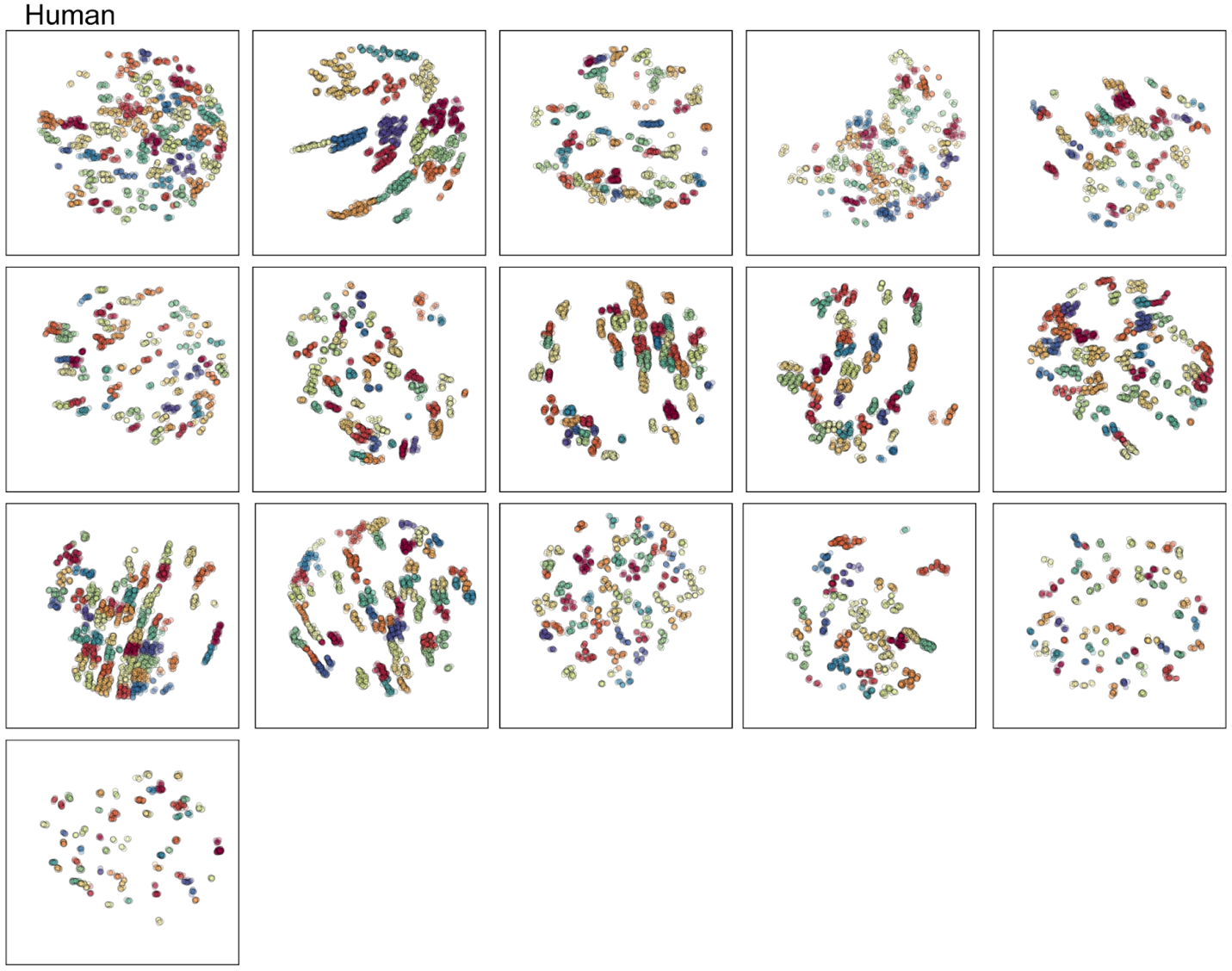

**Fig. S37. Human CD-PC tubule partitions in the medulla.** Spatial distribution of KMeans-derived partitions of CD-PC bead coordinates in the medulla, represented by unique colors across human arrays.

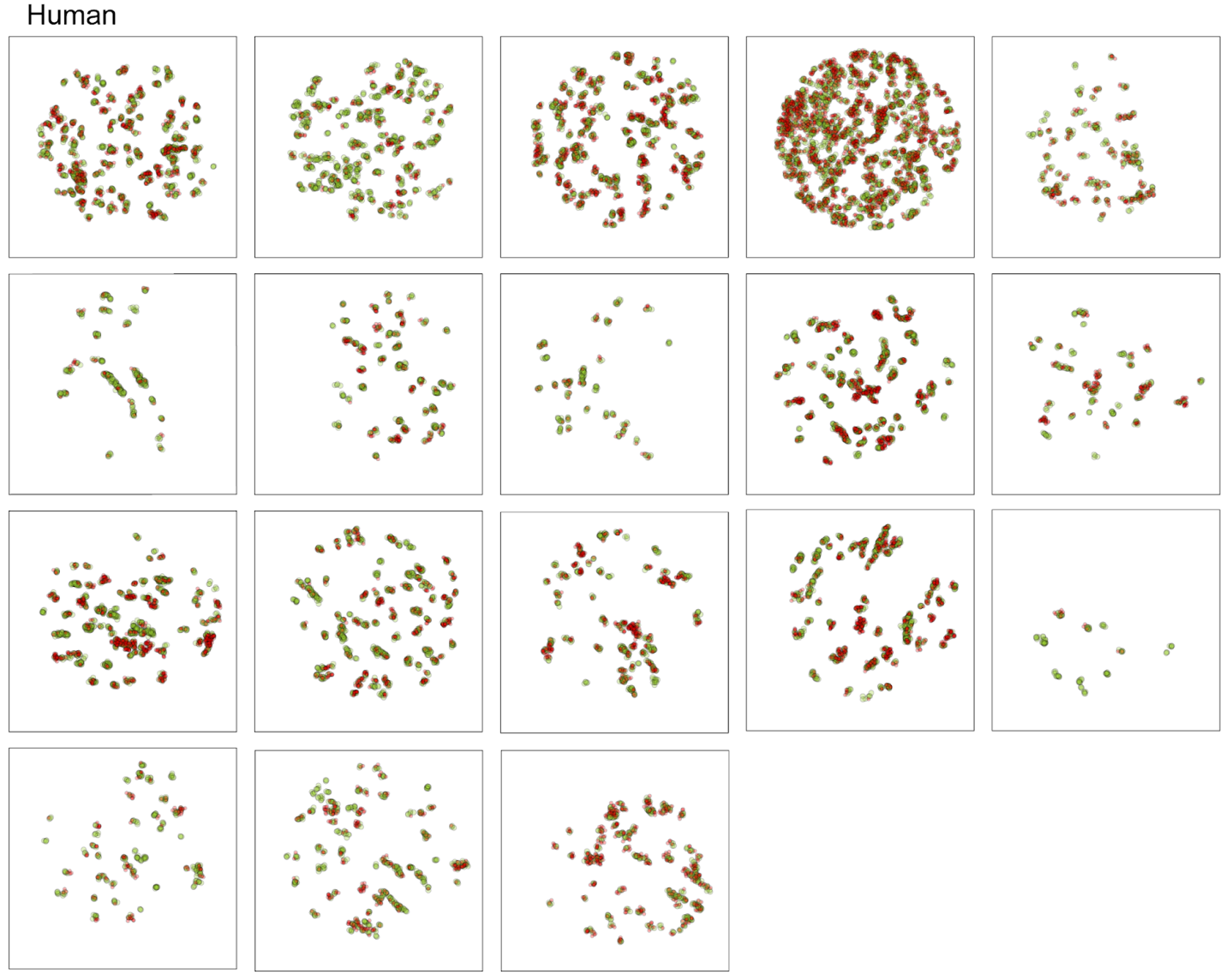

**Fig. S38. Human CD-IC tubule partitions in the cortex.** Spatial distribution of CD-IC partition assignments in the cortex (red), based on their nearest CD-PC tubule (green), across human arrays.

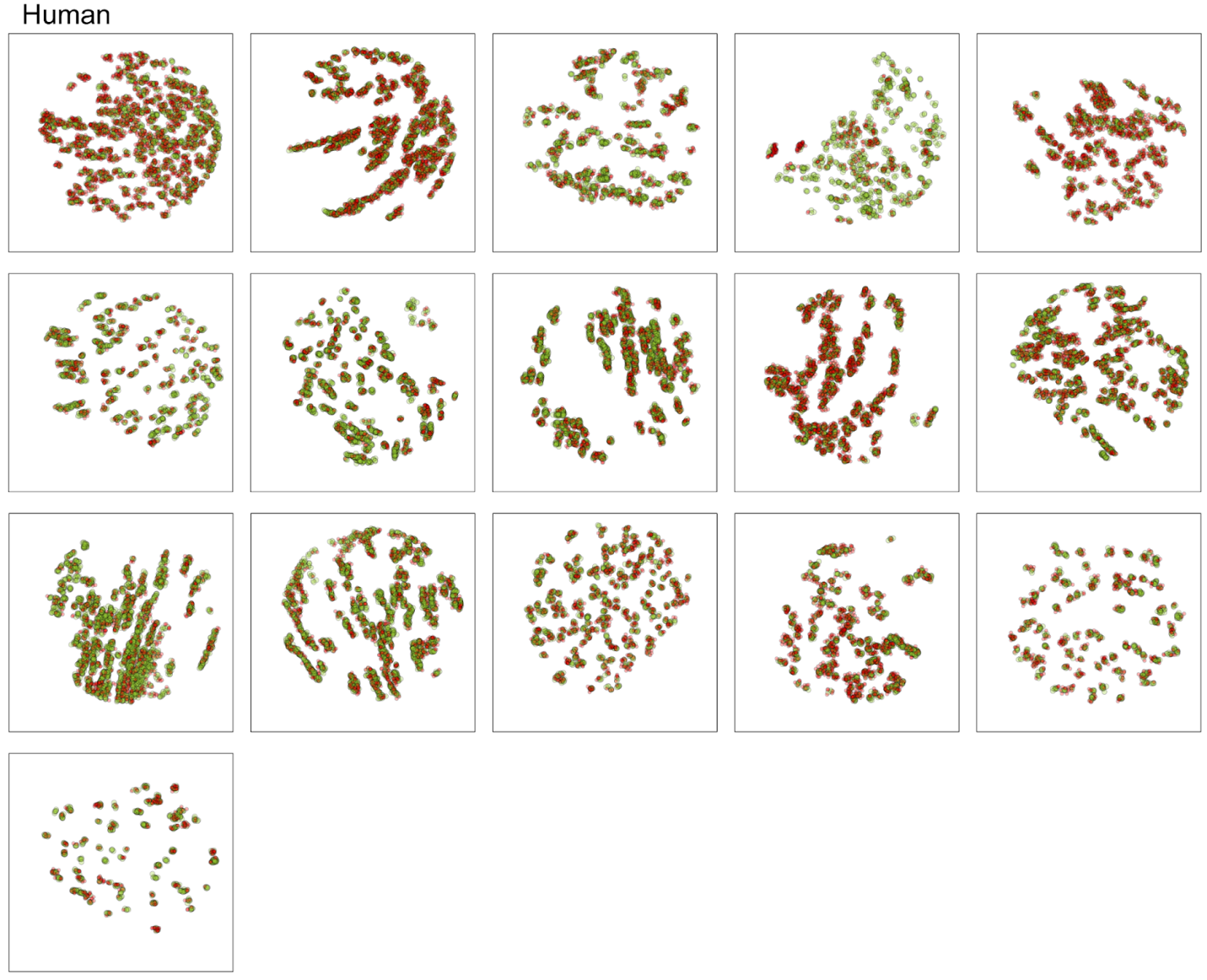

**Fig. S39. Human CD-IC tubule partitions in the medulla.** Spatial distribution of CD-IC partition assignments in the medulla (red), based on their nearest CD-PC tubule (green), across human arrays.

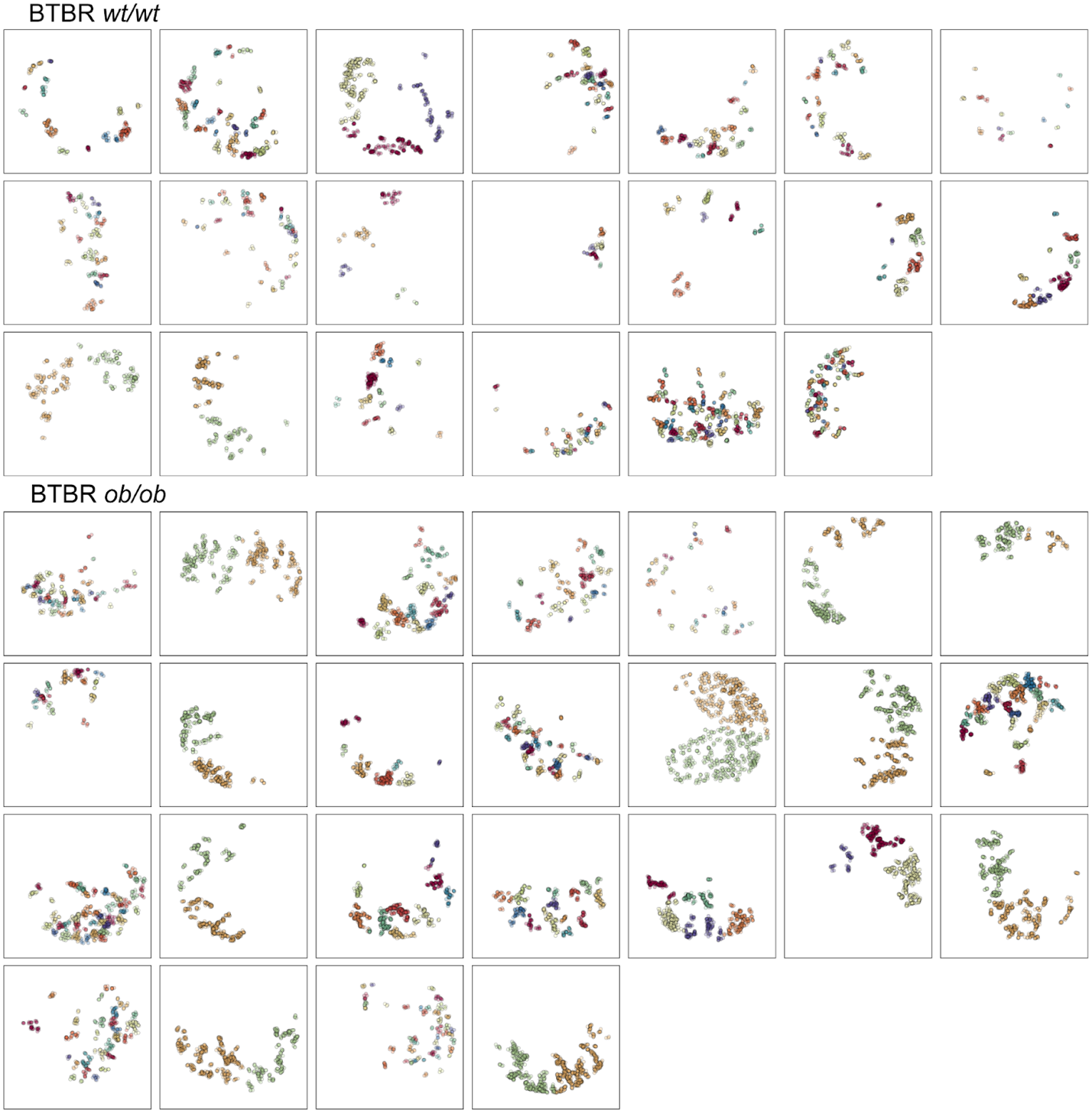

**Fig. S40. BTBR *wt/wt* and BTBR *ob/ob* DCT tubule partitions in the cortex.** Spatial distribution of KMeans-derived partitions of DCT bead coordinates in the cortex, represented by unique colors across BTBR *wt/wt* (top) and BTBR *ob/ob* (bottom) arrays.

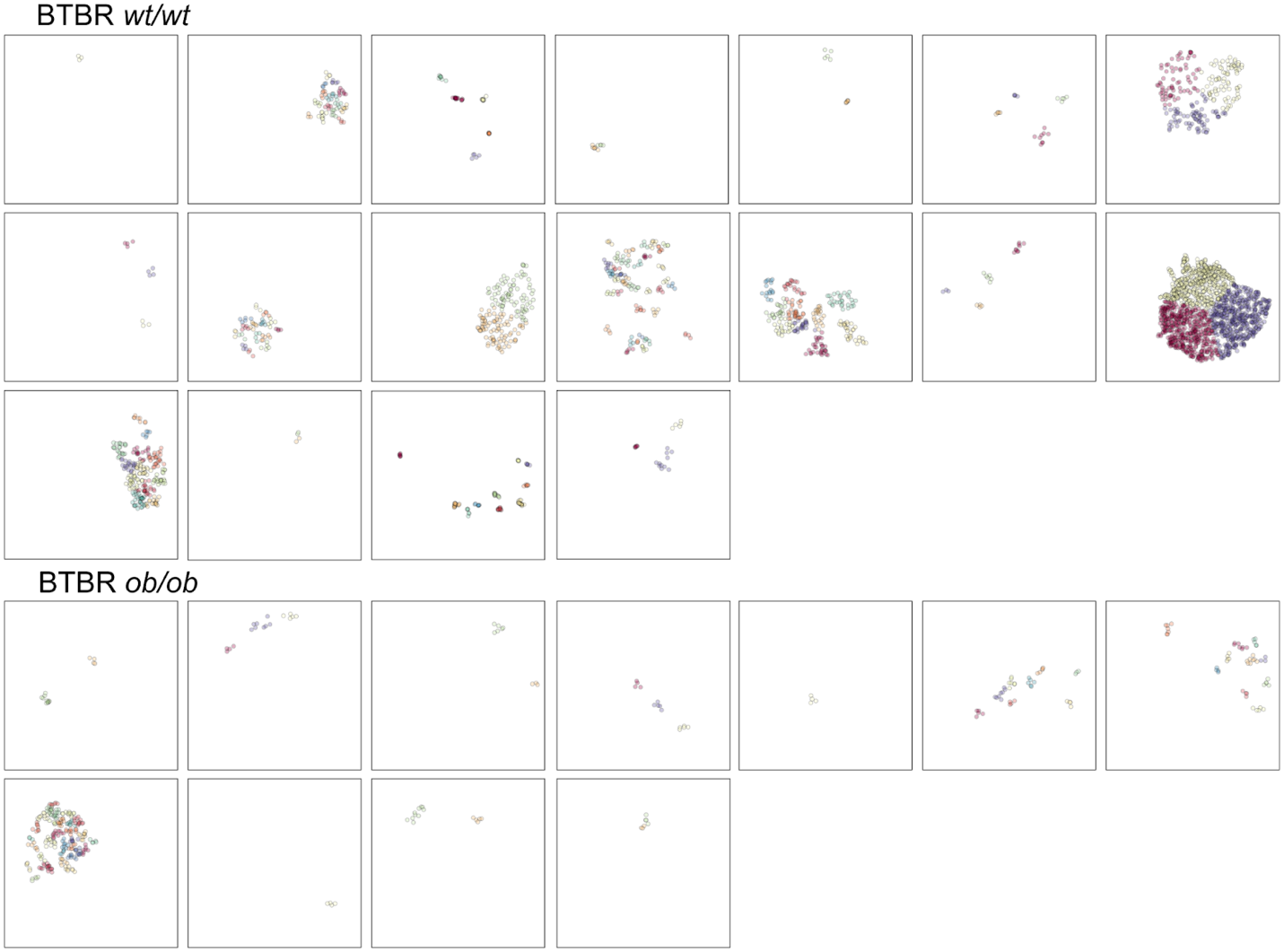

**Fig. S41. BTBR *wt/wt* and BTBR *ob/ob* DCT tubule partitions in the medulla.** Spatial distribution of KMeans-derived partitions of DCT bead coordinates in the medulla, represented by unique colors across BTBR *wt/wt* (top) and BTBR *ob/ob* (bottom) arrays.

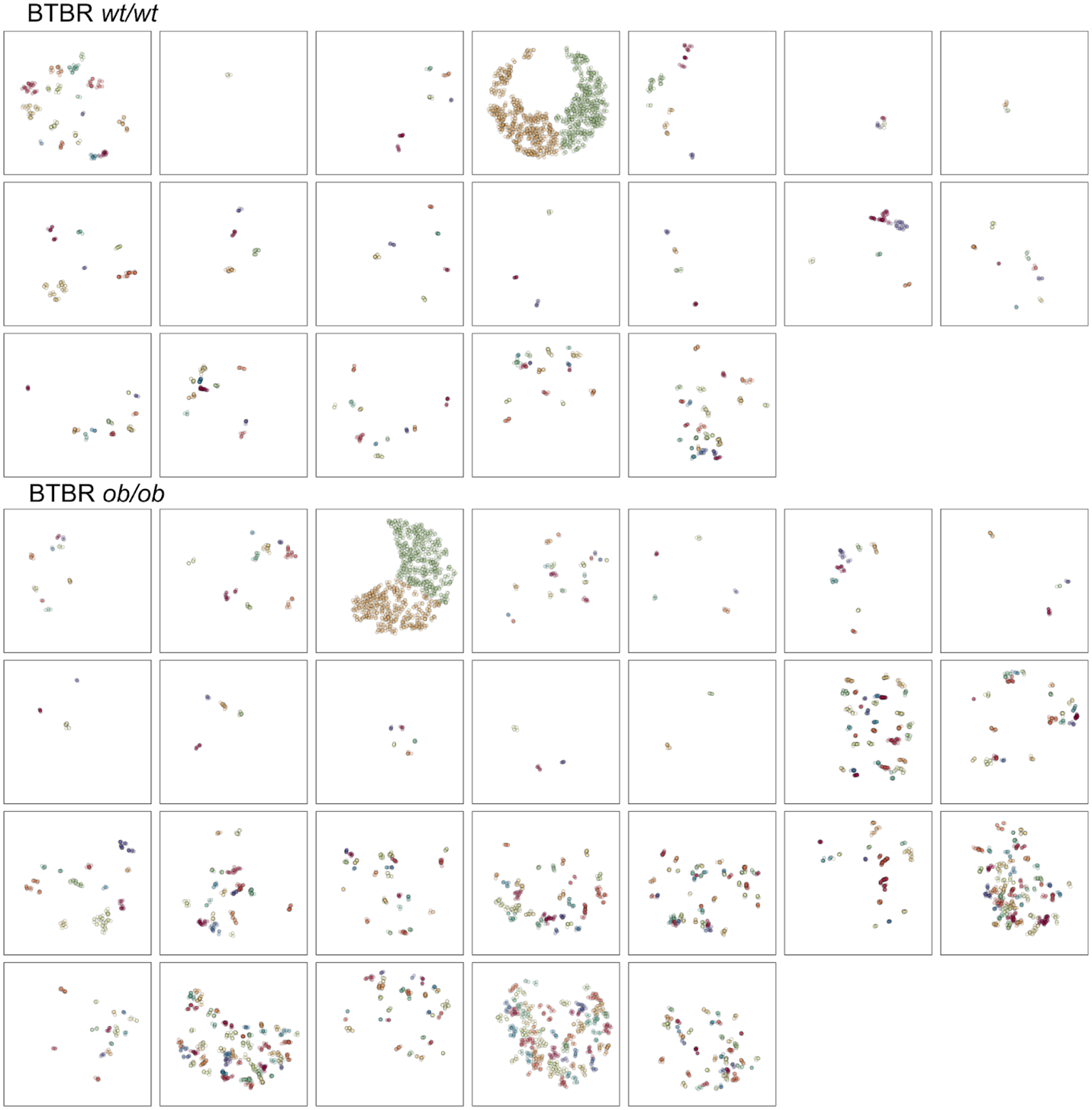

**Fig. S42. BTBR *wt/wt* and BTBR *ob/ob* CD-PC tubule partitions in the cortex.** Spatial distribution of KMeans-derived partitions of CD-PC bead coordinates in the cortex, represented by unique colors across BTBR *wt/wt* (top) and BTBR *ob/ob* (bottom) arrays.

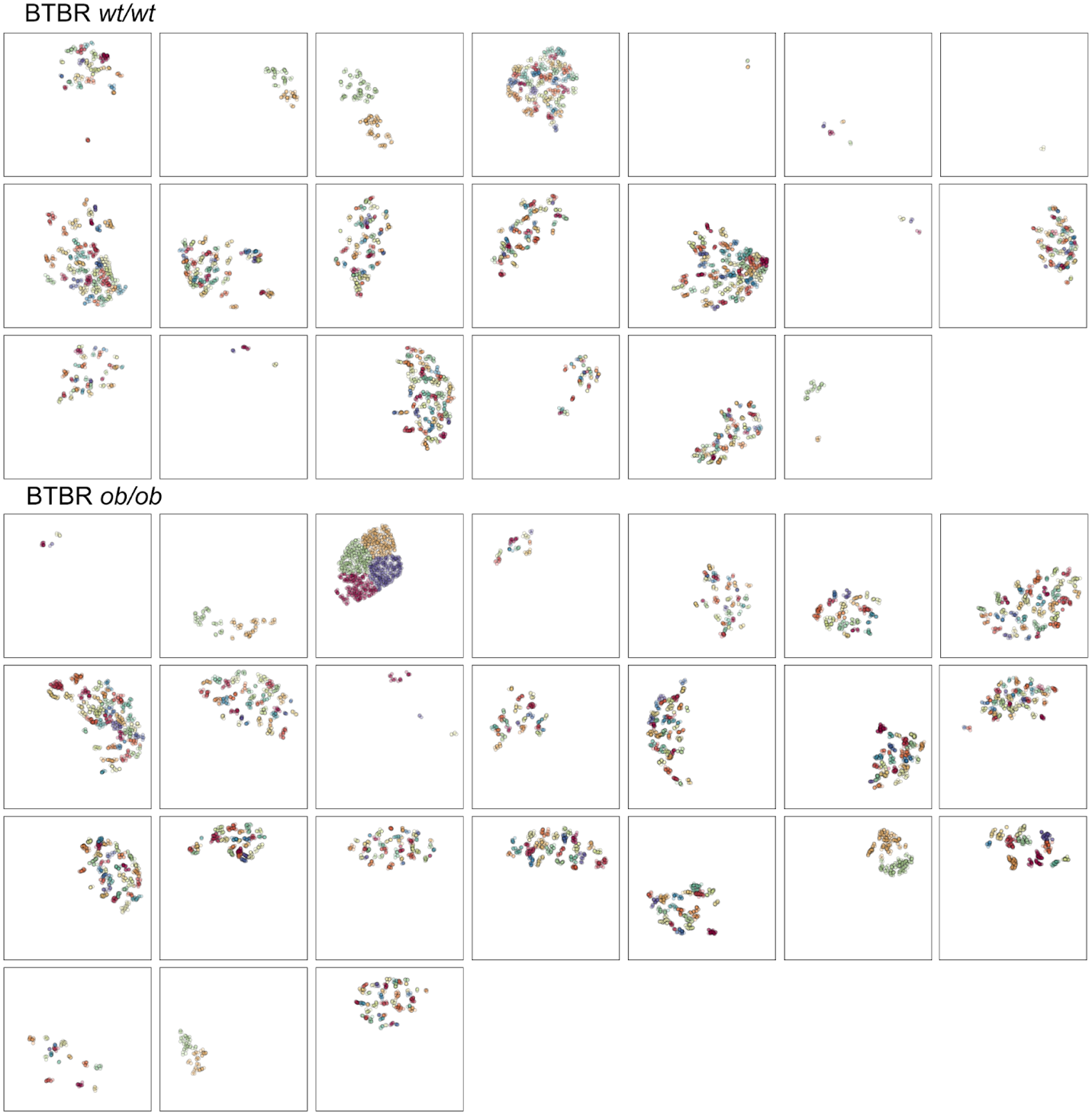

**Fig. S43. BTBR *wt/wt* and BTBR *ob/ob* CD-PC tubule partitions in the medulla.** Spatial distribution of KMeans-derived partitions of CD-PC bead coordinates in the medulla, represented by unique colors across BTBR *wt/wt* (top) and BTBR *ob/ob* (bottom) arrays.

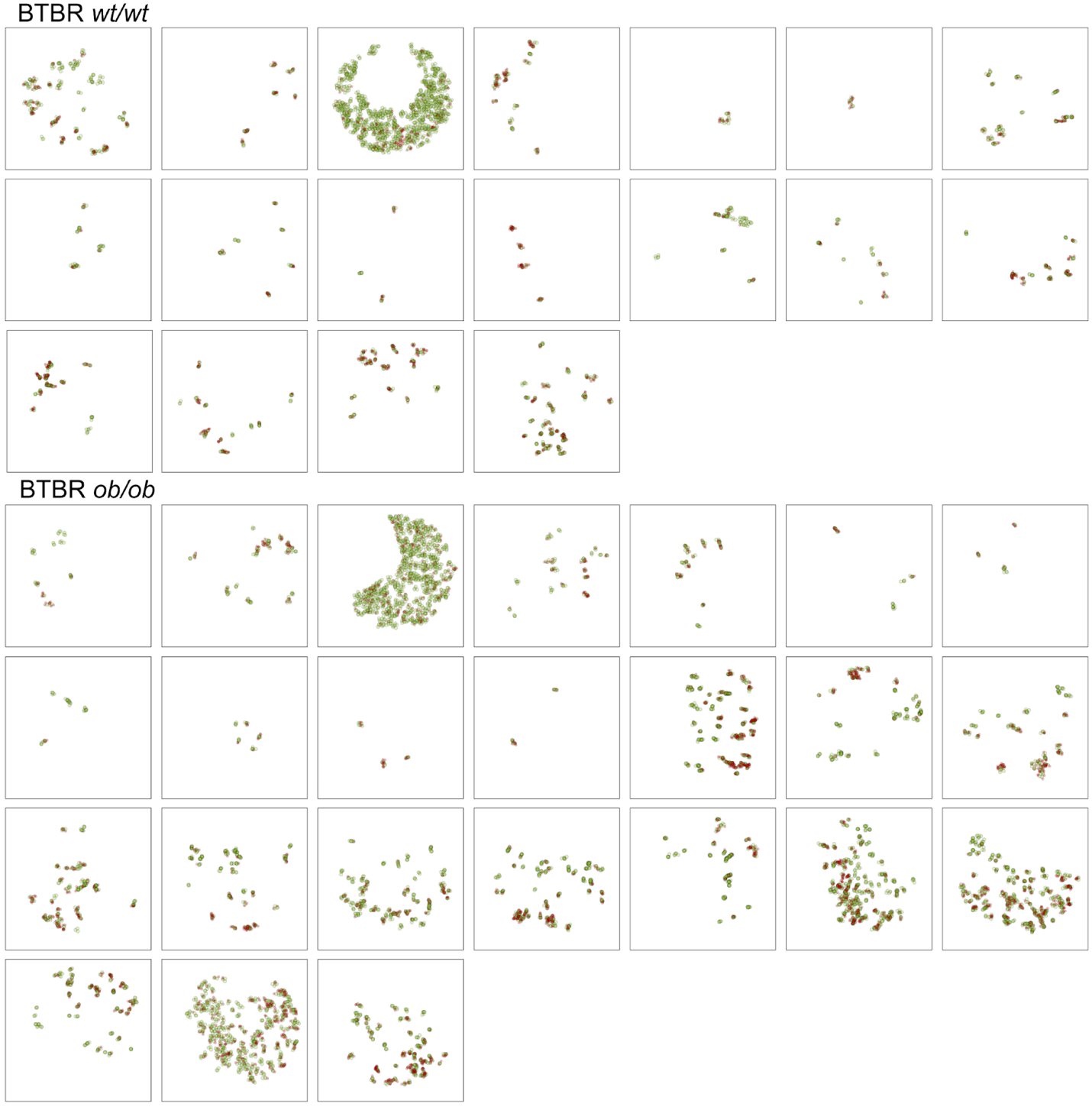

**Fig. S44. BTBR *wt/wt* and BTBR *ob/ob* CD-IC tubule partitions in the cortex.** Spatial distribution of CD-IC partition assignments in the cortex (red), based on their nearest CD-PC tubule (green), across BTBT *wt/wt* (top) and BTBR *ob/ob* (bottom) arrays.

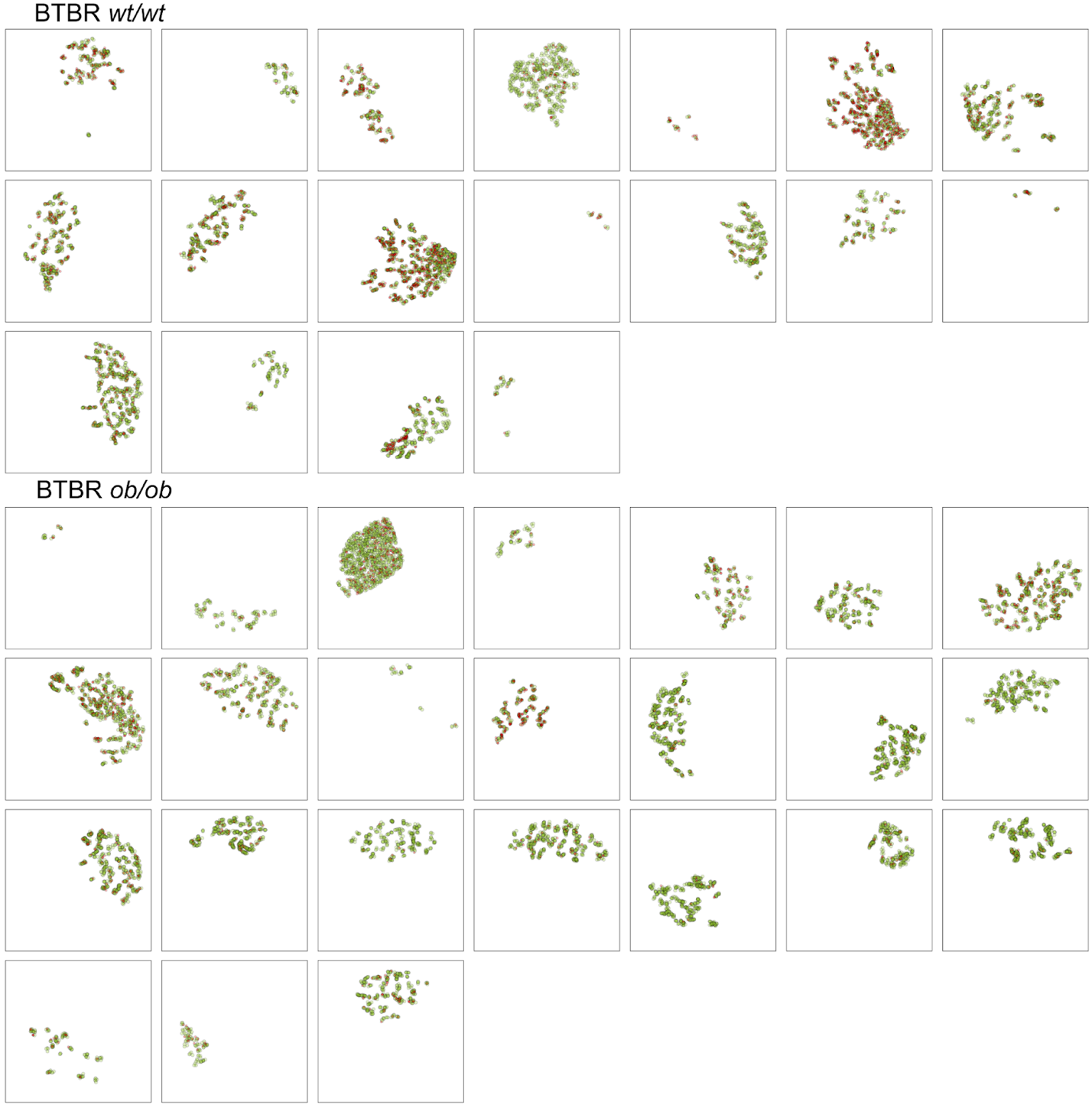

**Fig. S45. BTBR *wt/wt* and BTBR *ob/ob* CD-IC tubule partitions in the medulla.** Spatial distribution of CD-IC partition assignments in the medulla (red), based on their nearest CD-PC tubule (green), across BTBT *wt/wt* (top) and BTBR *ob/ob* (bottom) arrays.

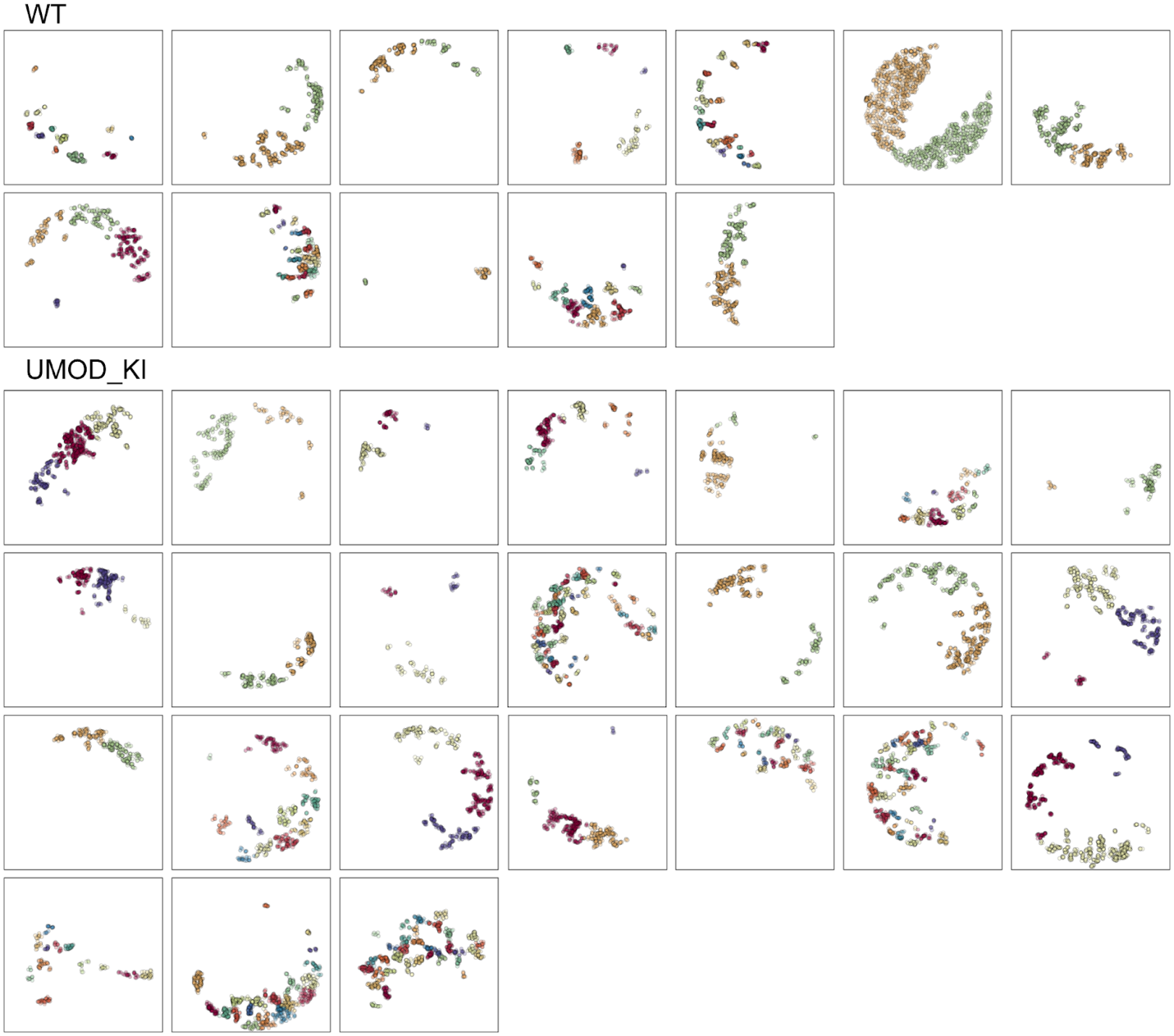

**Fig. S46. WT and UMOD-KI DCT tubule partitions in the cortex.** Spatial distribution of KMeans-derived partitions of DCT bead coordinates in the cortex, represented by unique colors across WT (top) and UMOD-KI (bottom) arrays.

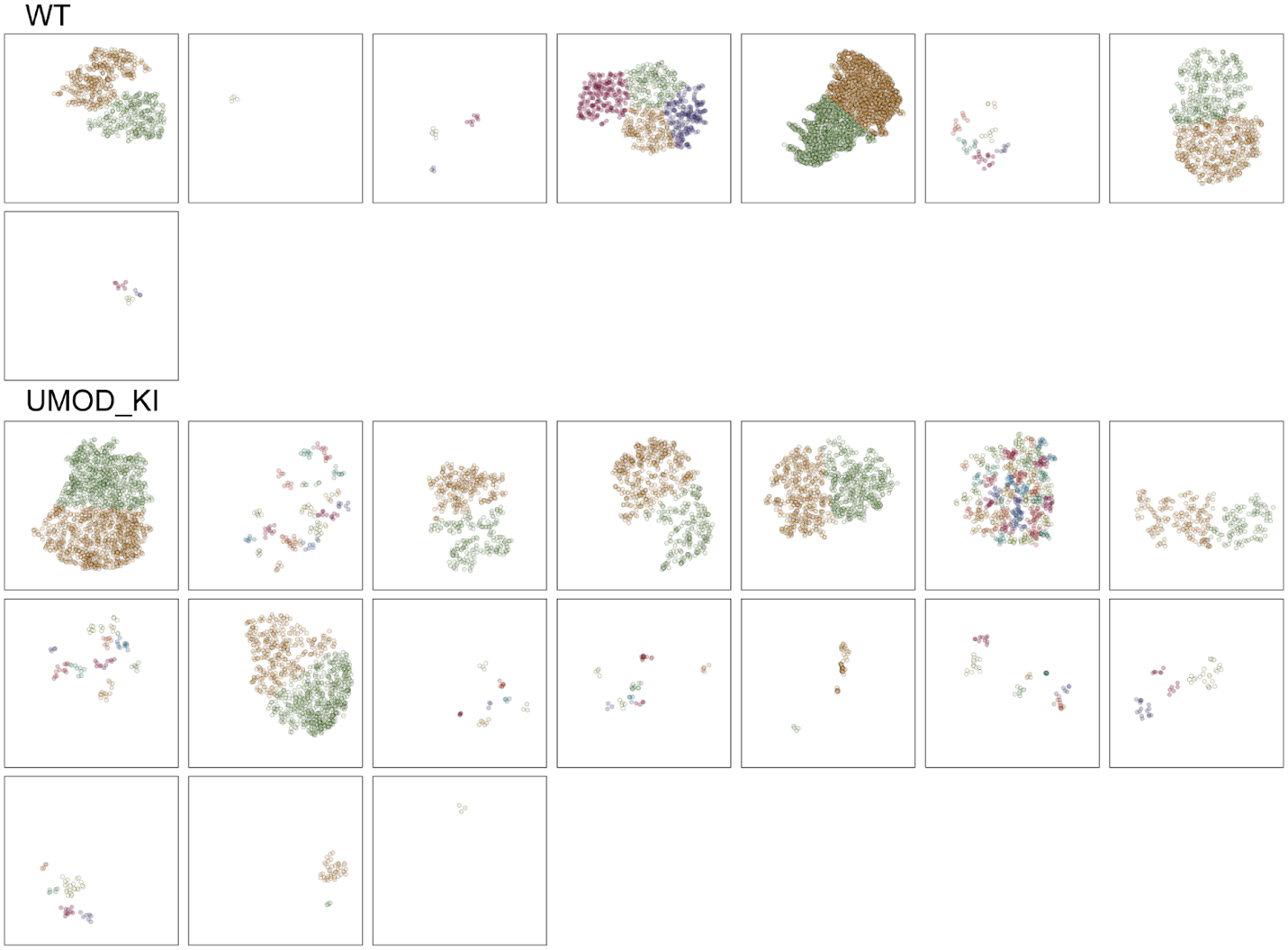

**Fig. S47. WT and UMOD-KI DCT tubule partitions in the medulla.** Spatial distribution of KMeans-derived partitions of DCT bead coordinates in the medulla, represented by unique colors across WT (top) and UMOD-KI (bottom) arrays.

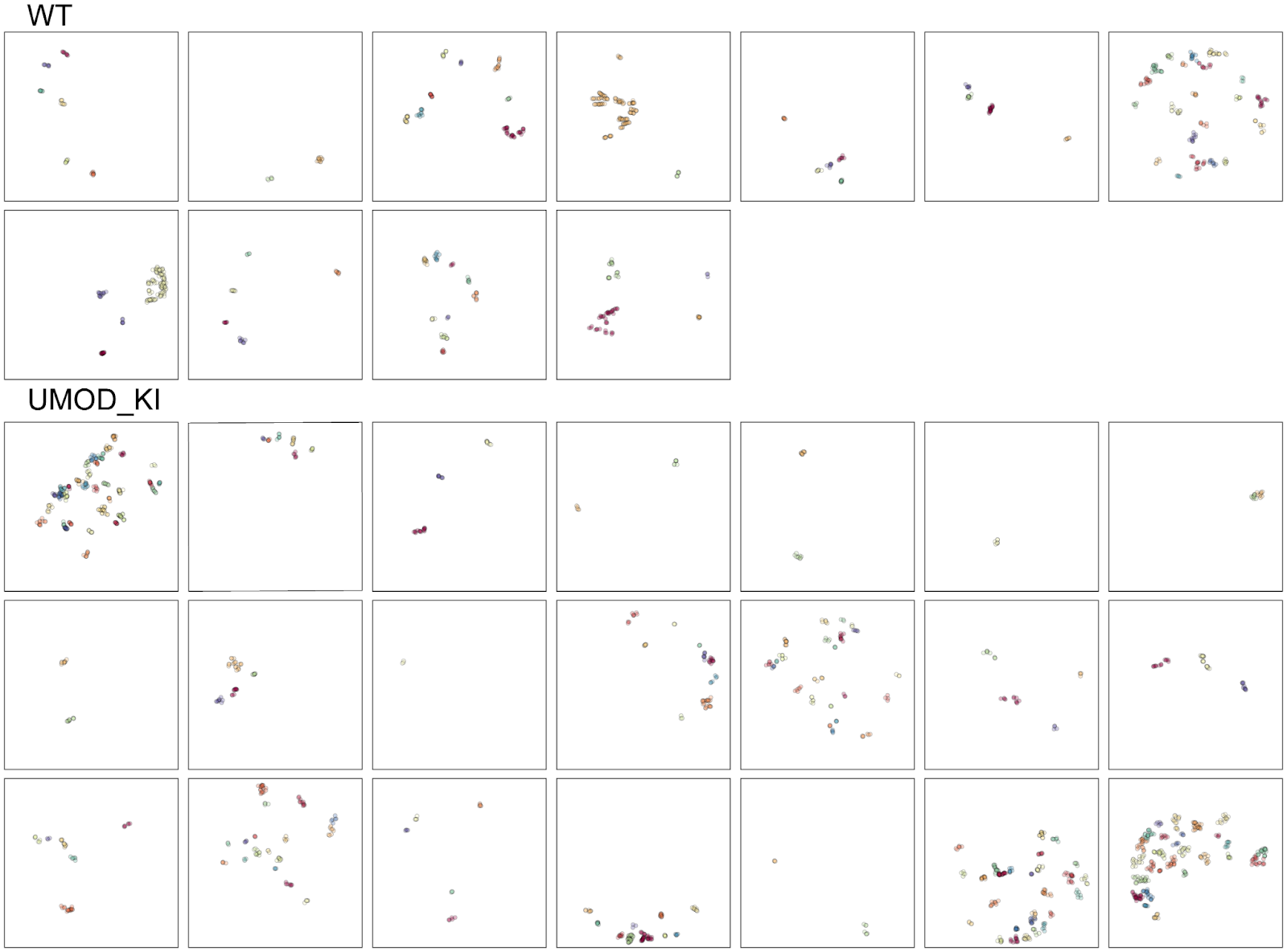

**Fig. S48. WT and UMOD-KI CD-PC tubule partitions in the cortex.** Spatial distribution of KMeans-derived partitions of CD-PC bead coordinates in the cortex, represented by unique colors across WT (top) and UMOD-KI (bottom) arrays.

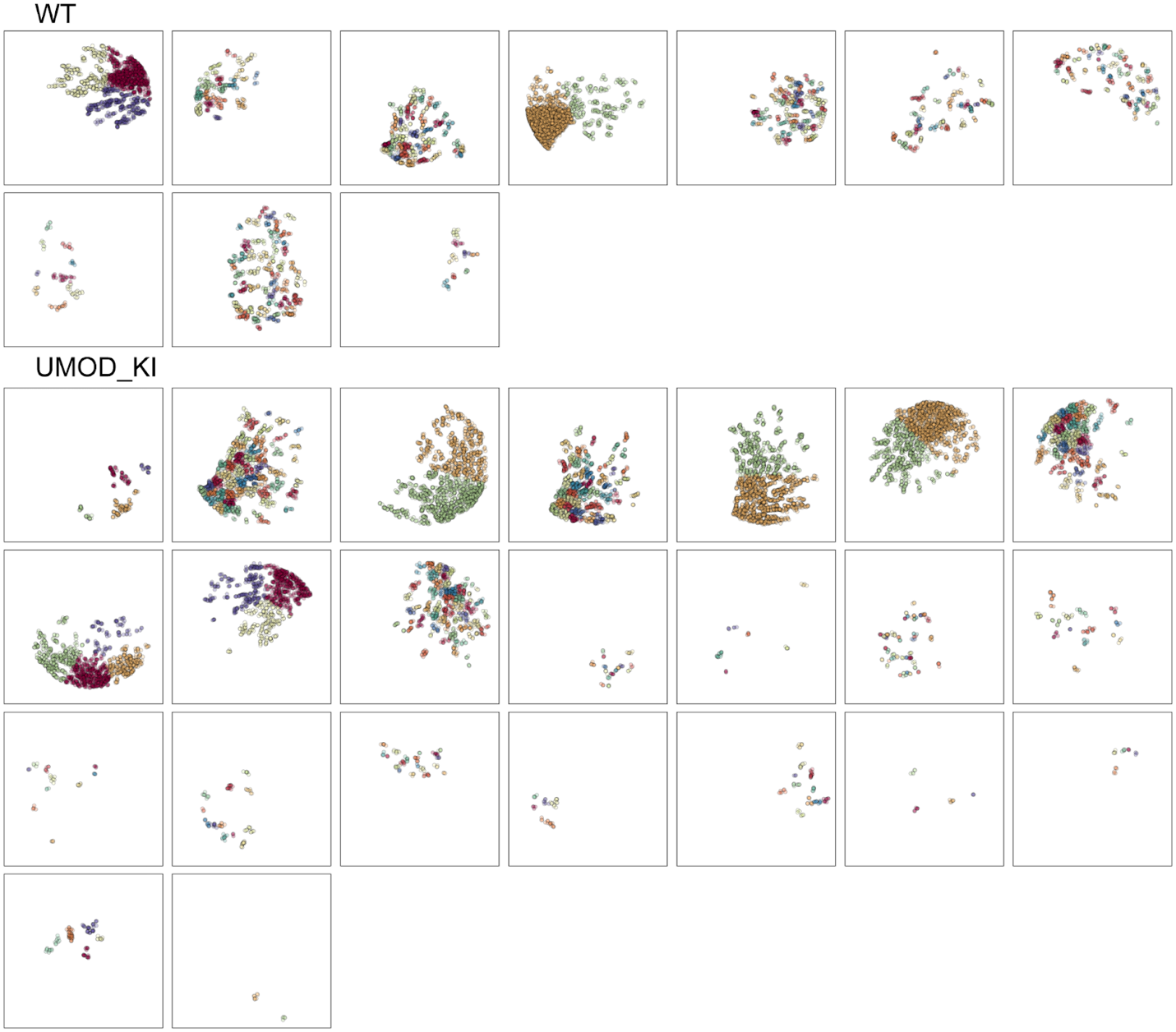

**Fig. S49. WT and UMOD-KI CD-PC tubule partitions in the medulla.** Spatial distribution of KMeans-derived partitions of CD-PC bead coordinates in the medulla, represented by unique colors across WT (top) and UMOD-KI (bottom) arrays.

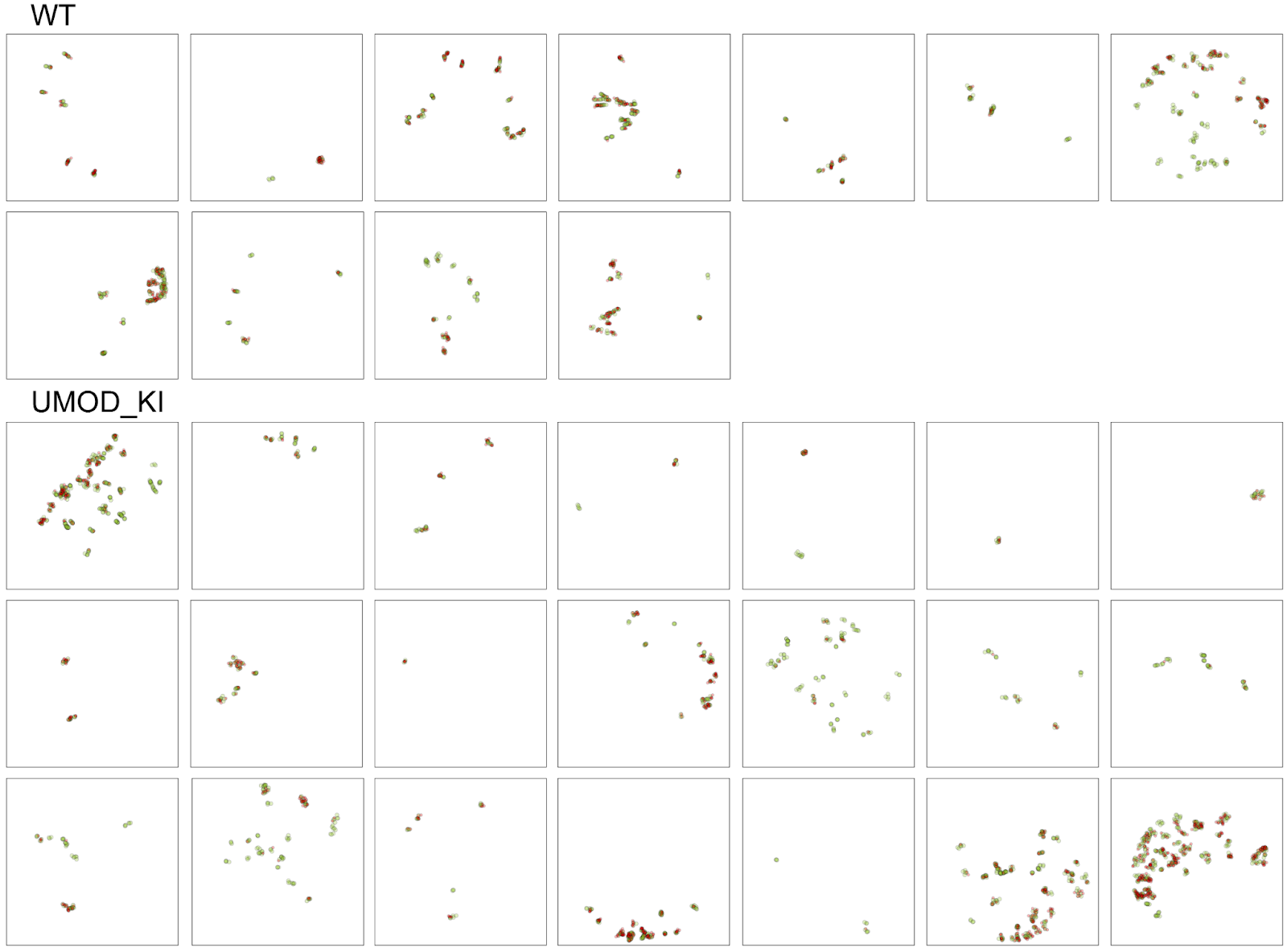

**Fig. S50. WT and UMOD-KI CD-IC tubule partitions in the cortex.** Spatial distribution of CD-IC partition assignments in the cortex (red), based on their nearest CD-PC tubule (green), across WT (top) and UMOD-KI (bottom) arrays.

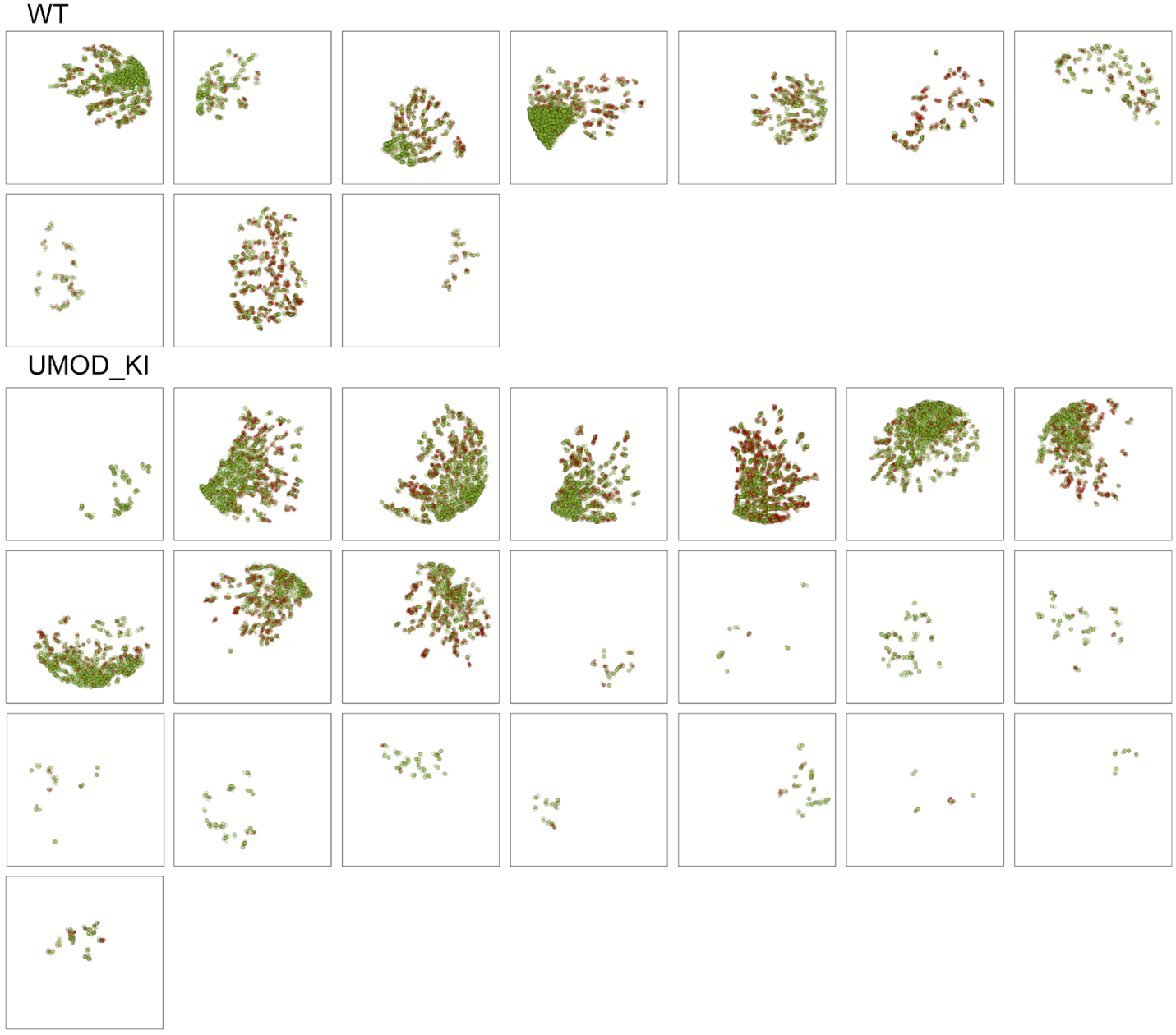

**Fig. S51. WT and UMOD-KI CD-IC tubule partitions in the medulla.** Spatial distribution of CD-IC partition assignments in the medulla (red), based on their nearest CD-PC tubule (green), across WT (top) and UMOD-KI (bottom) arrays.

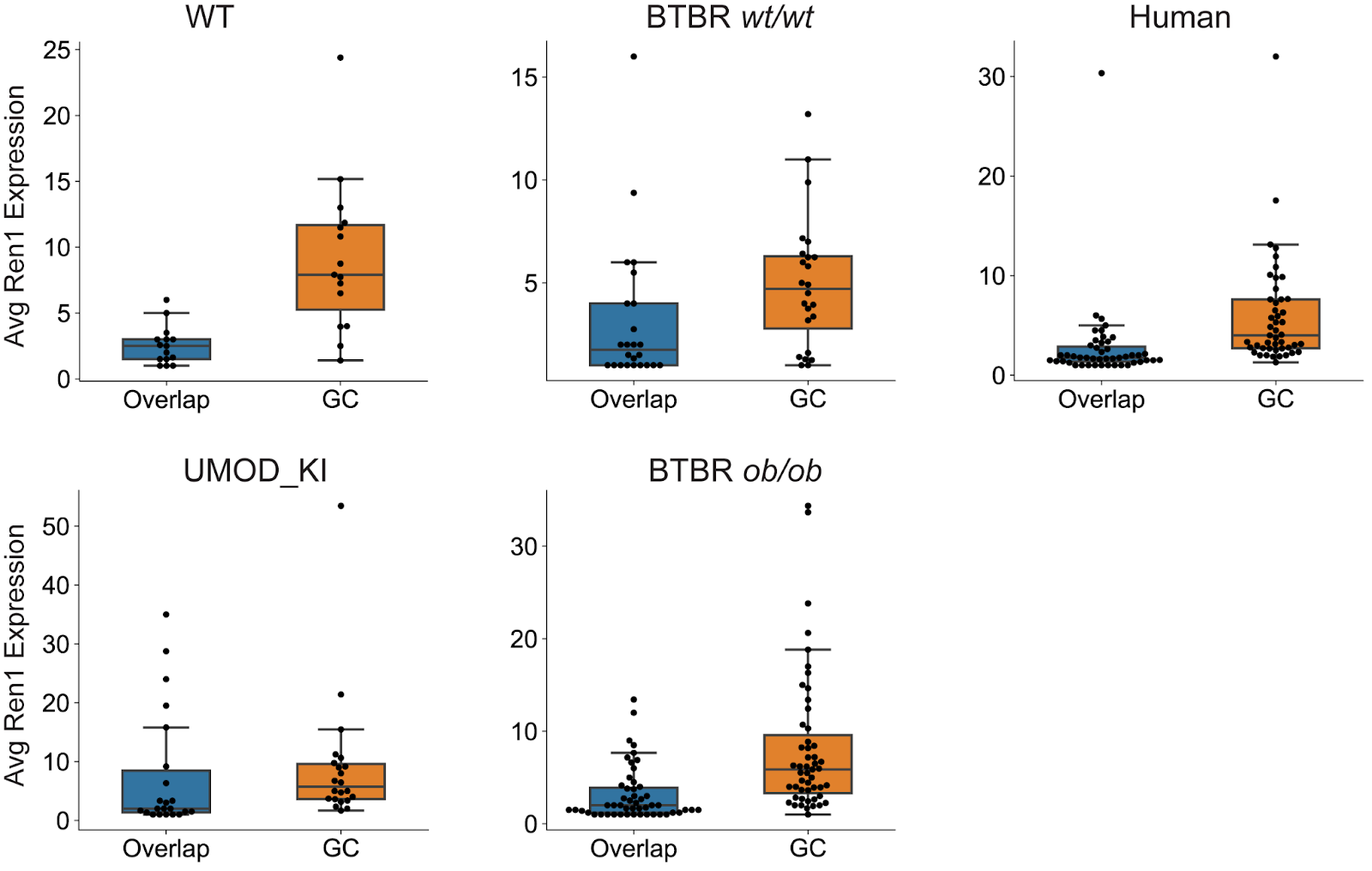

**Fig. S52. Average Ren1/REN expression was higher in beads uniquely identified in GC clusters than in GC beads overlapping with glomerular cell calls.** Boxplots of average expression of GC beads per cluster overlapping with glomerular cell calls (left) and uniquely identified as GC  (right). Lower, middle, and upper horizontal lines of the boxes represent the first quartile, median, and third quartile, respectively. Lower and upper whiskers represent 1.5*IQR + first quartile and 1.5*IQR + third quartile, respectively.

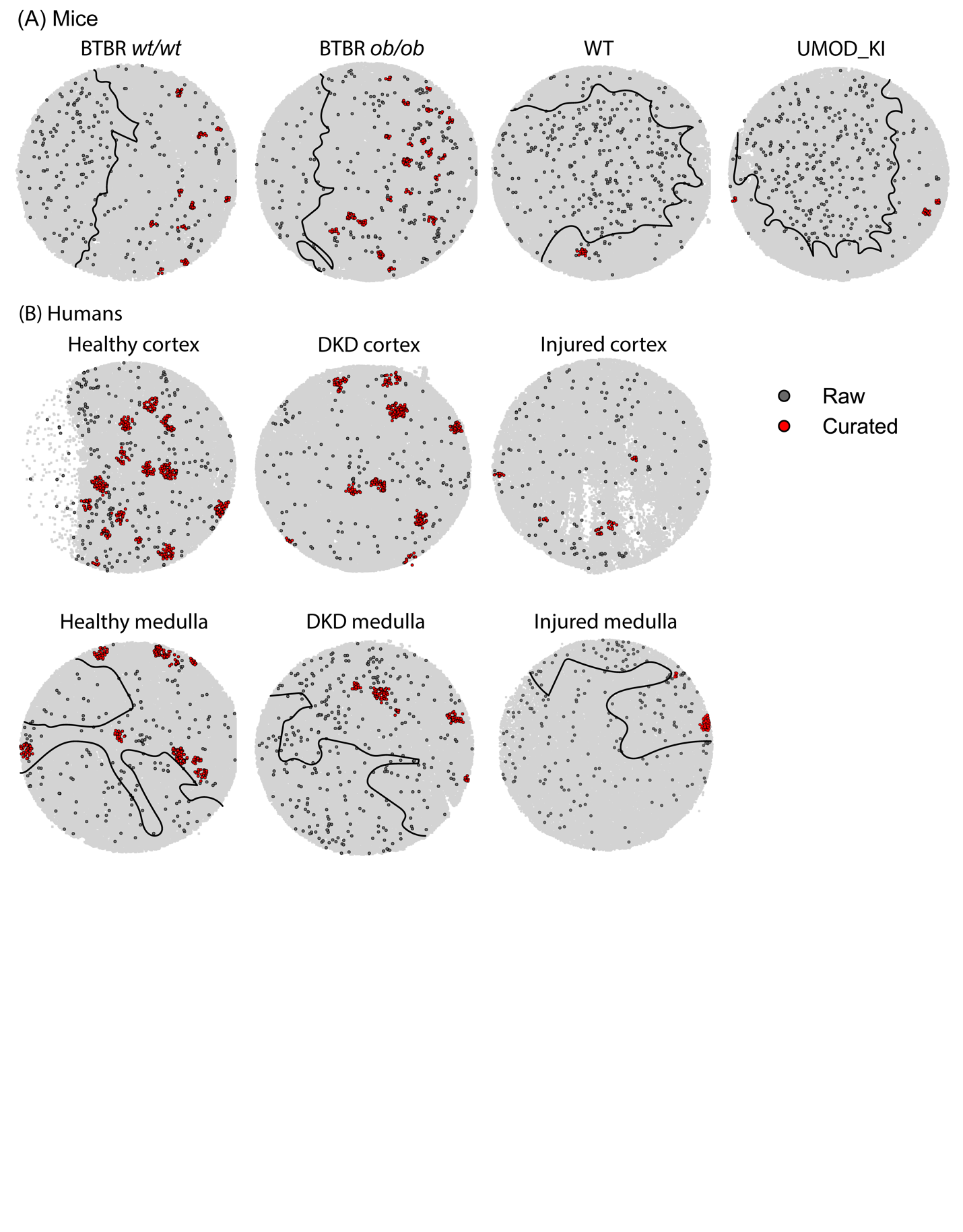

**Fig. S53. Raw versus KNN-curated podocyte calls in mice and humans.** Spatial distribution of raw NMFreg podocyte calls (dark grey) (with spatial outliers removed), and curated podocyte calls from KNN filtration (red) for (**A**) mouse and (**B**) human arrays. Black outline shows approximate delineation of cortex and medulla regions of each array.

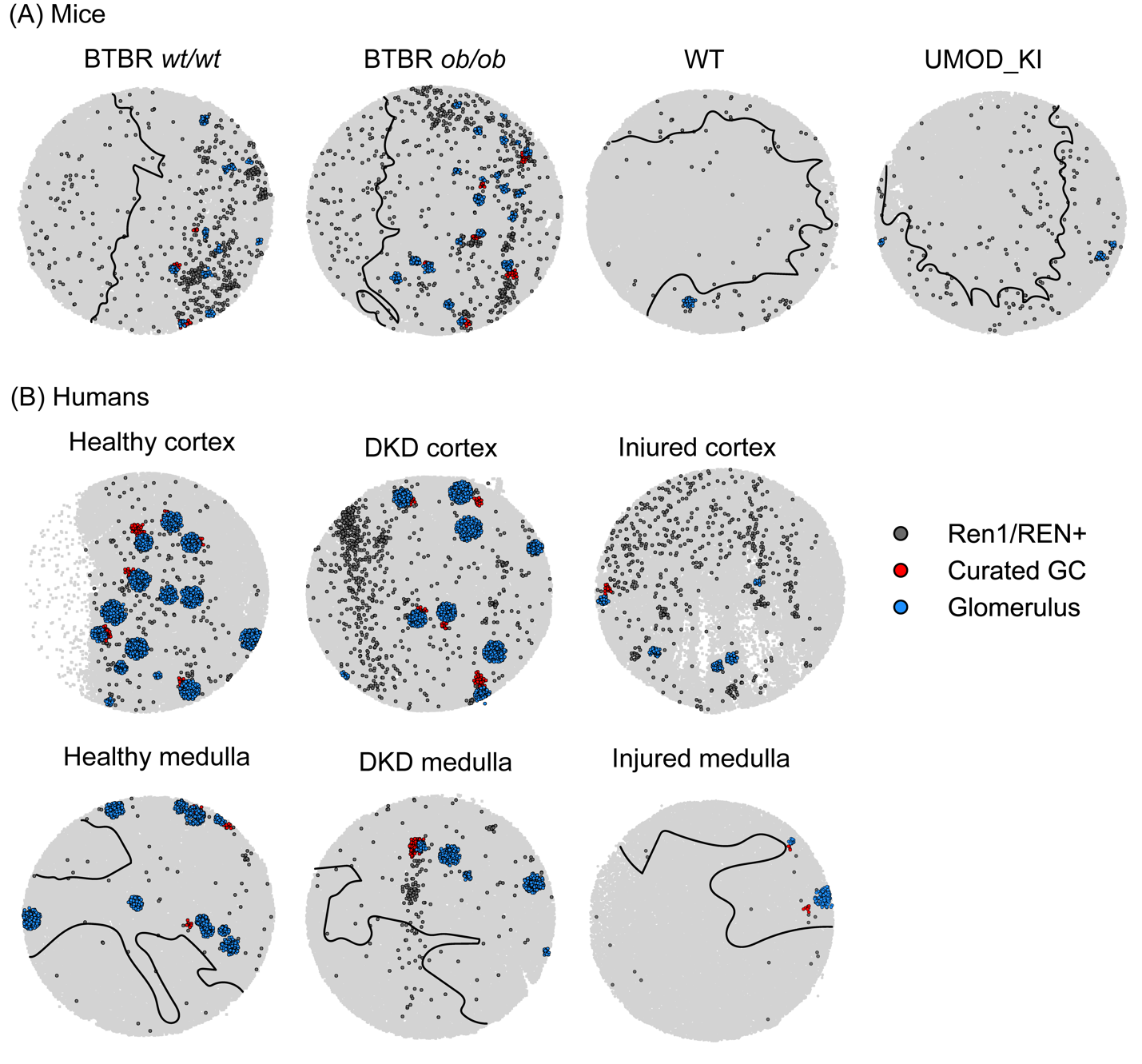

**Fig. S54. Granular cell curation in mice and humans.** Spatial distribution of *Ren1*/*REN*+ beads (dark grey) and curated granular cell calls defined as dense clusters of *Ren1*/*REN*+ beads by KNN filtration within close proximity to glomeruli (red) in (**A**) mice and (**B**) humans.

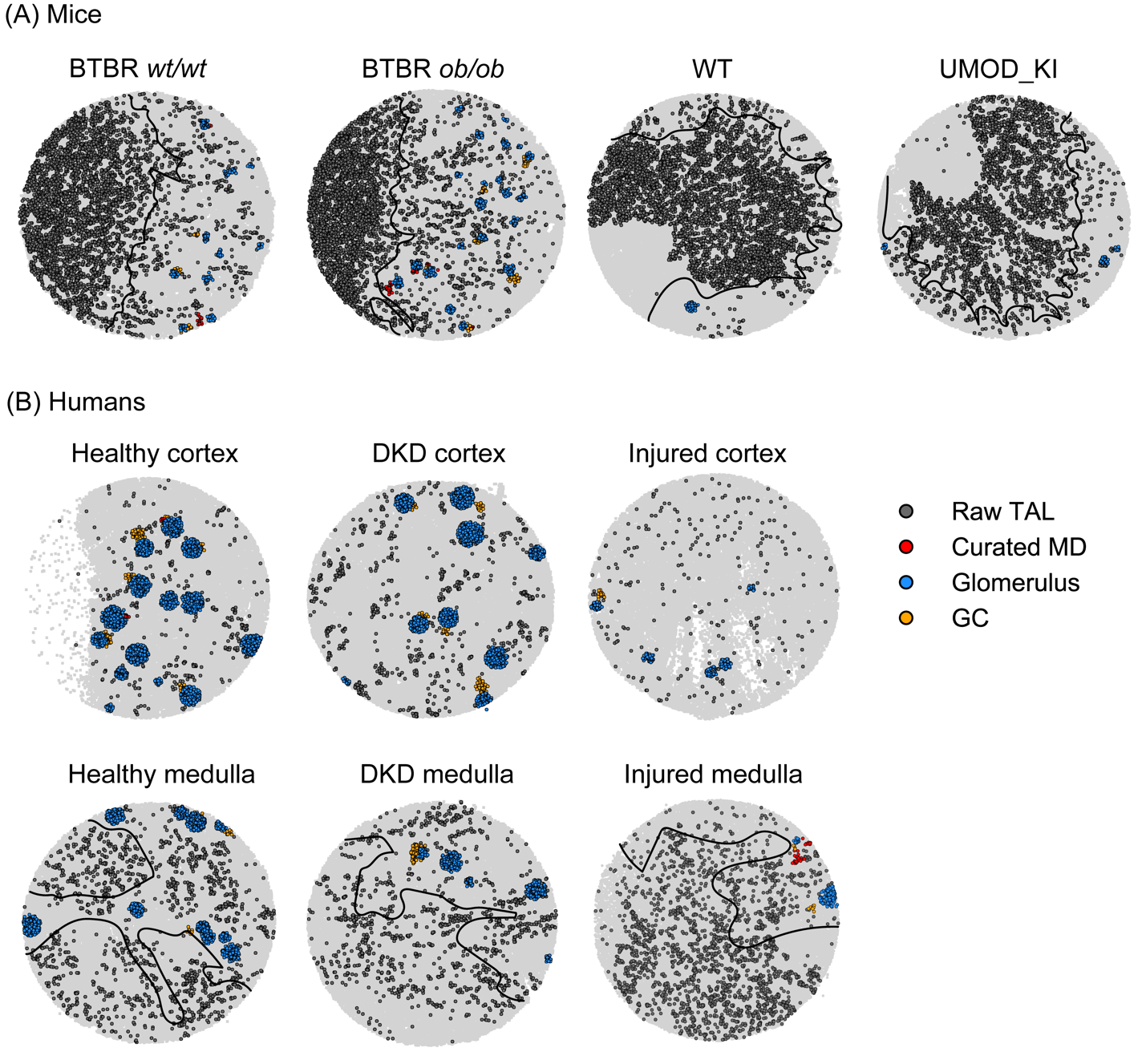

**Fig. S55. MD curation in mice and humans.** Spatial distribution of raw TAL calls from Seurat (dark grey) and curated MD calls defined as dense clusters of TAL beads by KNN filtration expressing MD markers and within close proximity to glomeruli (red) in (**A**) mice and (**B**) humans.

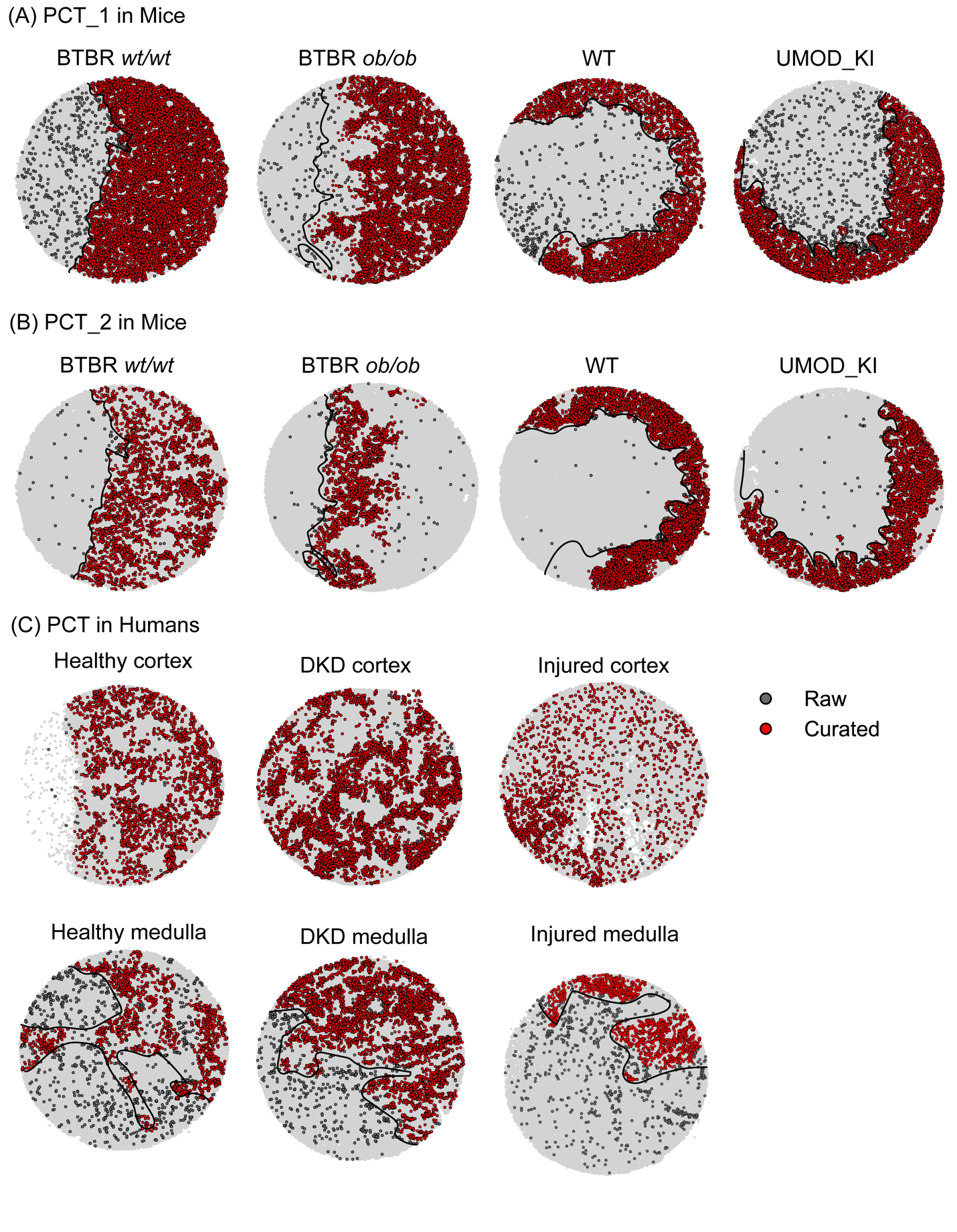

**Fig. S56. Raw versus KNN-curated PCT calls in mice and humans.** Spatial distribution of raw Seurat PCT calls (dark grey) (with spatial outliers and glomerular, GC, and MD areas removed), and (**A**) curated PCT_1 calls in mice, (**B**) curated PCT_2 calls in mice, and (**C**) curated PCT calls in humans from KNN filtration and removal of beads from medulla (red). Black outline shows approximate delineation of cortex and medulla regions of each array.

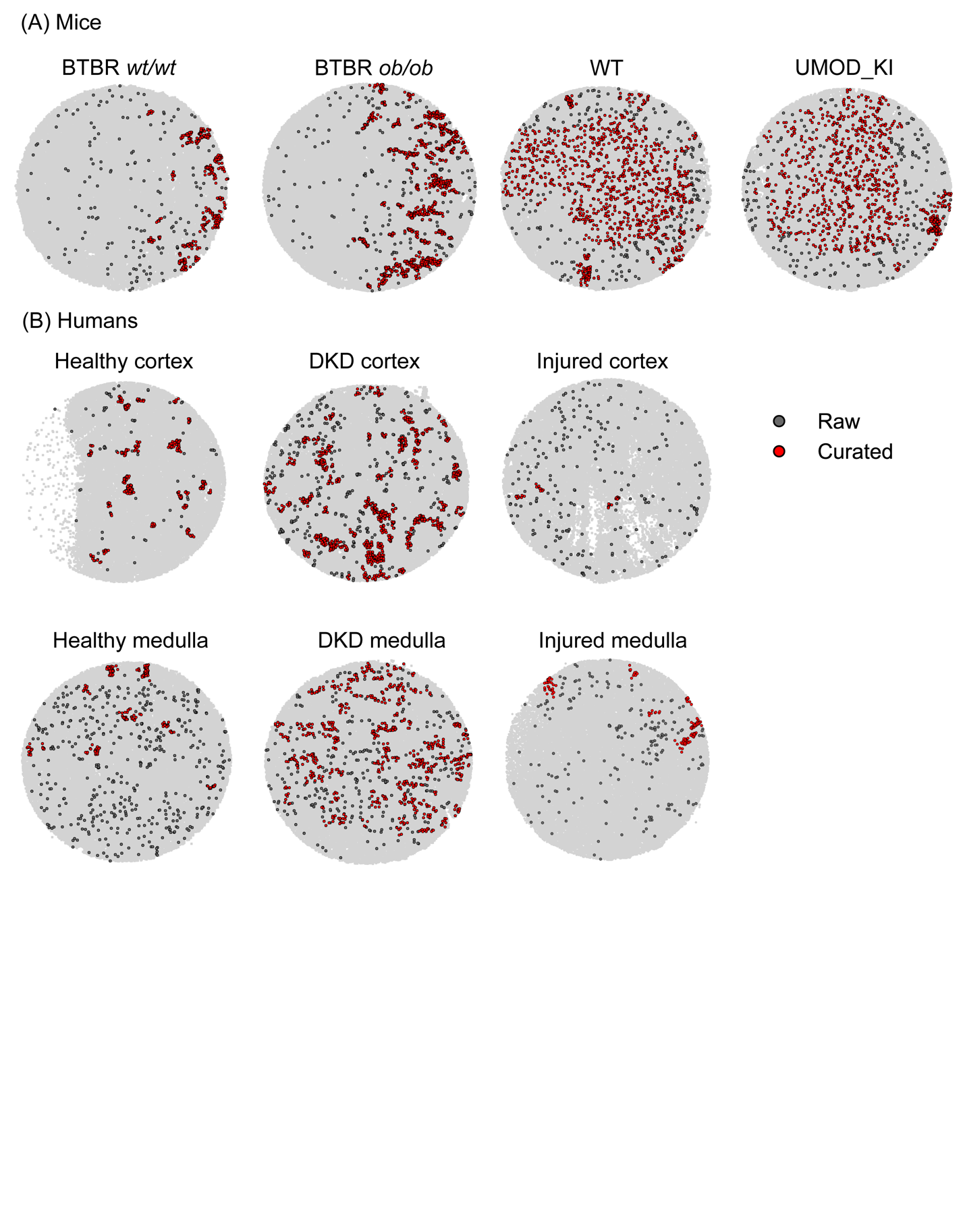

**Fig. S57. Raw versus KNN-curated DCT calls in mice and humans.** Spatial distribution of raw Seurat DCT calls (dark grey) (with spatial outliers and glomerular, GC, and MD areas removed), and curated DCT calls from KNN filtration (red) in (**A**) mice and (**B**) humans.

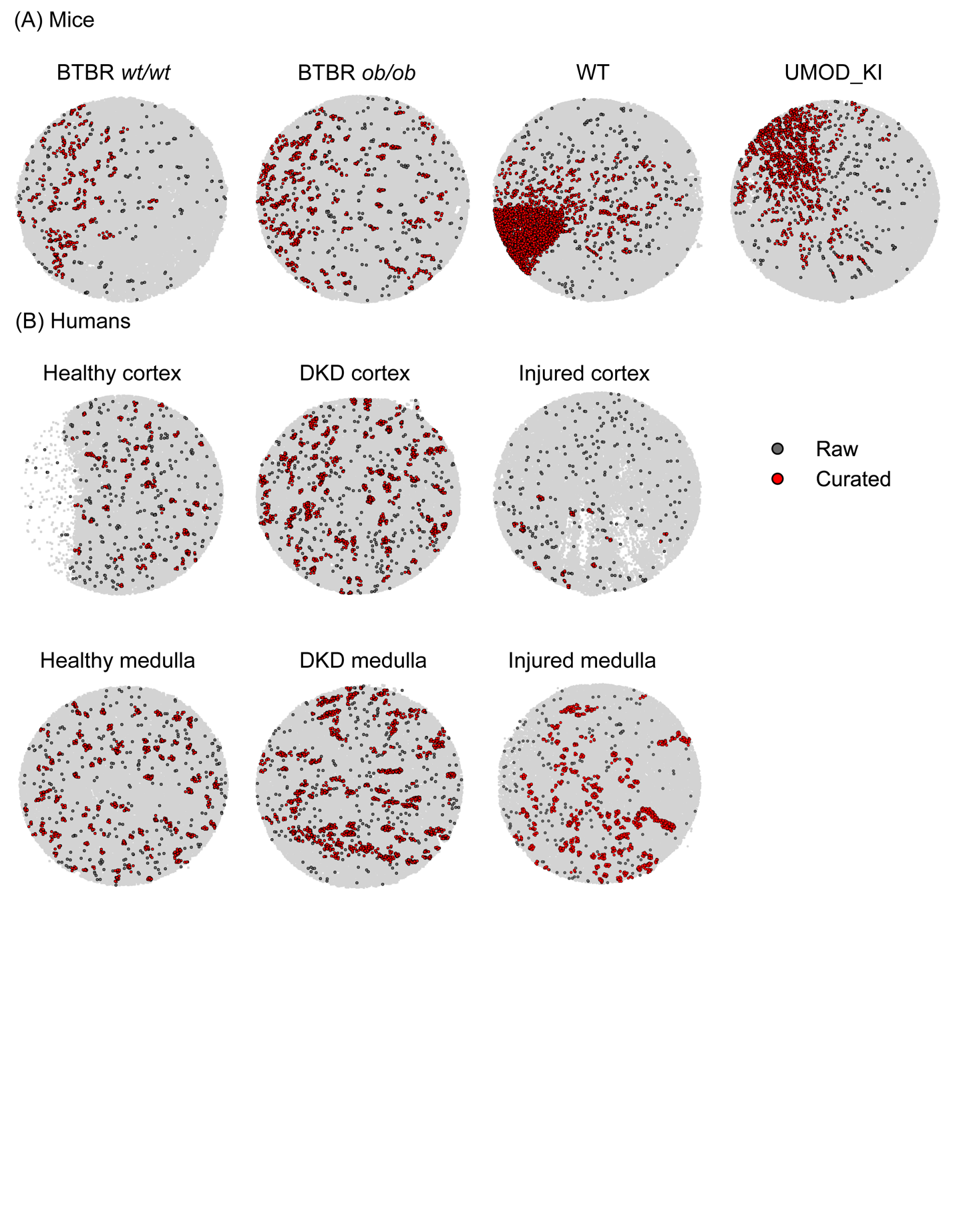

**Fig. S58. Raw versus KNN-curated CD-PC calls in mice and humans.** Spatial distribution of raw Seurat CD-PC calls (dark grey) (with spatial outliers and glomerular, GC, and MD areas removed), and curated CD-PC calls from KNN filtration (red) in (**A**) mice and (**B**) humans.

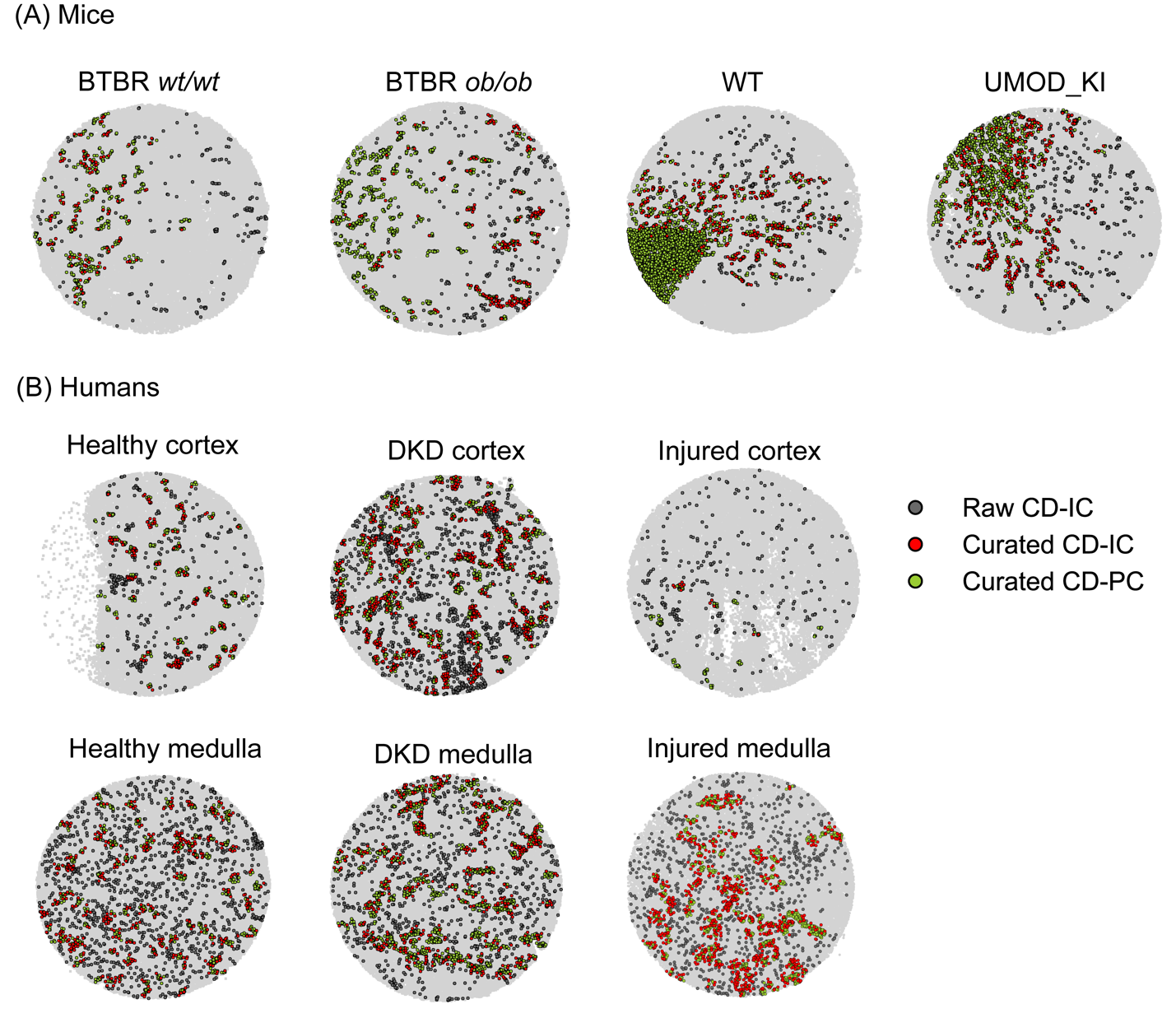

**Fig. S59. CD-IC curation in mice and humans.** Spatial distribution of raw Seurat CD-IC calls (dark grey) (with spatial outliers and glomerular, GC, and MD areas removed), curated CD-PC calls from KNN filtration (green), and curated CD-IC calls defined as those occurring within close proximity to CD-PC tubules (red) in (**A**) mice and (**B**) humans.

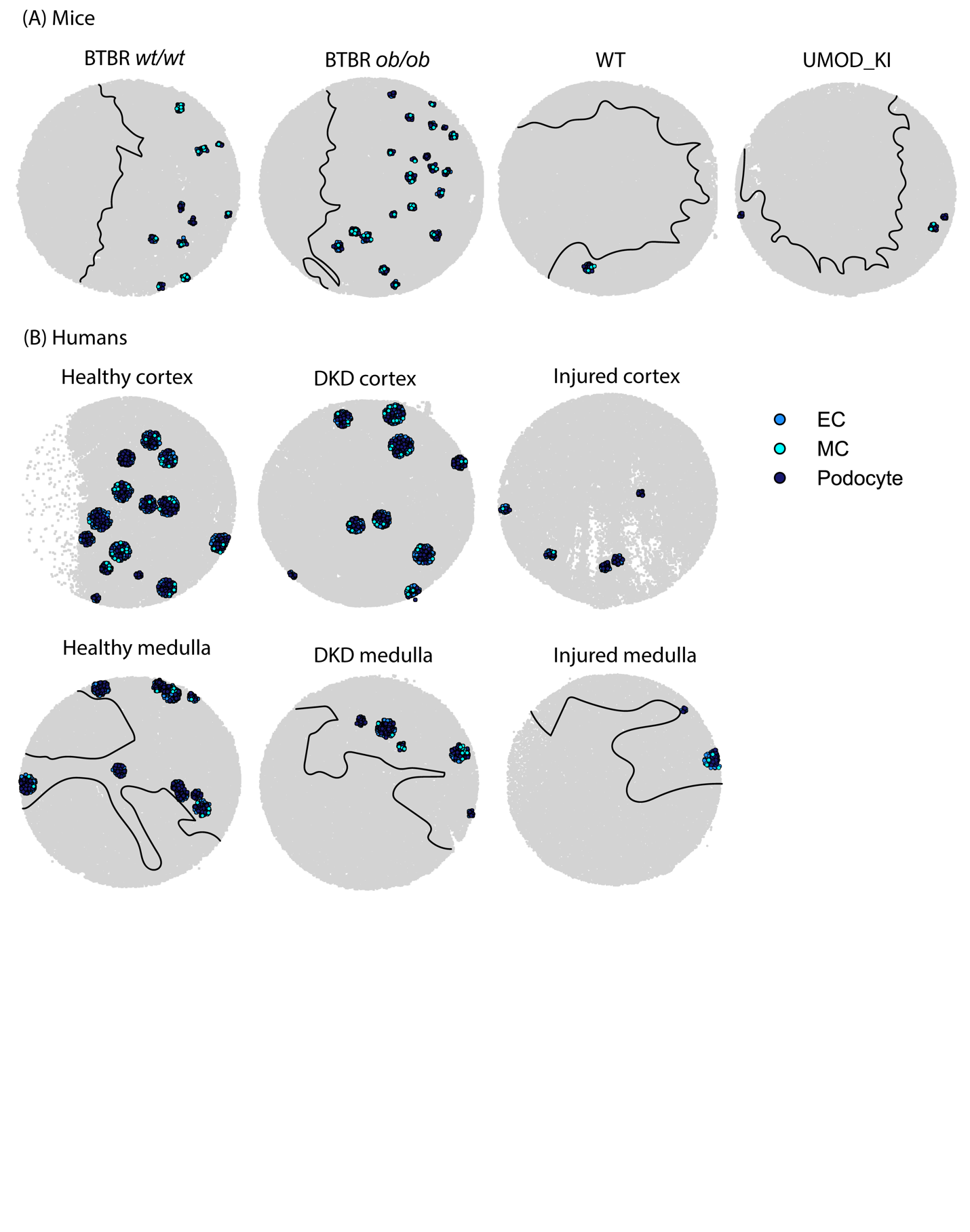

**Fig. S60. Cell type assignment in glomeruli in mice and humans.** Assignment of EC (blue) and MC (cyan) in glomerular areas defined by KNN-curated podocytes in (**A**) mice and (**B**) humans.

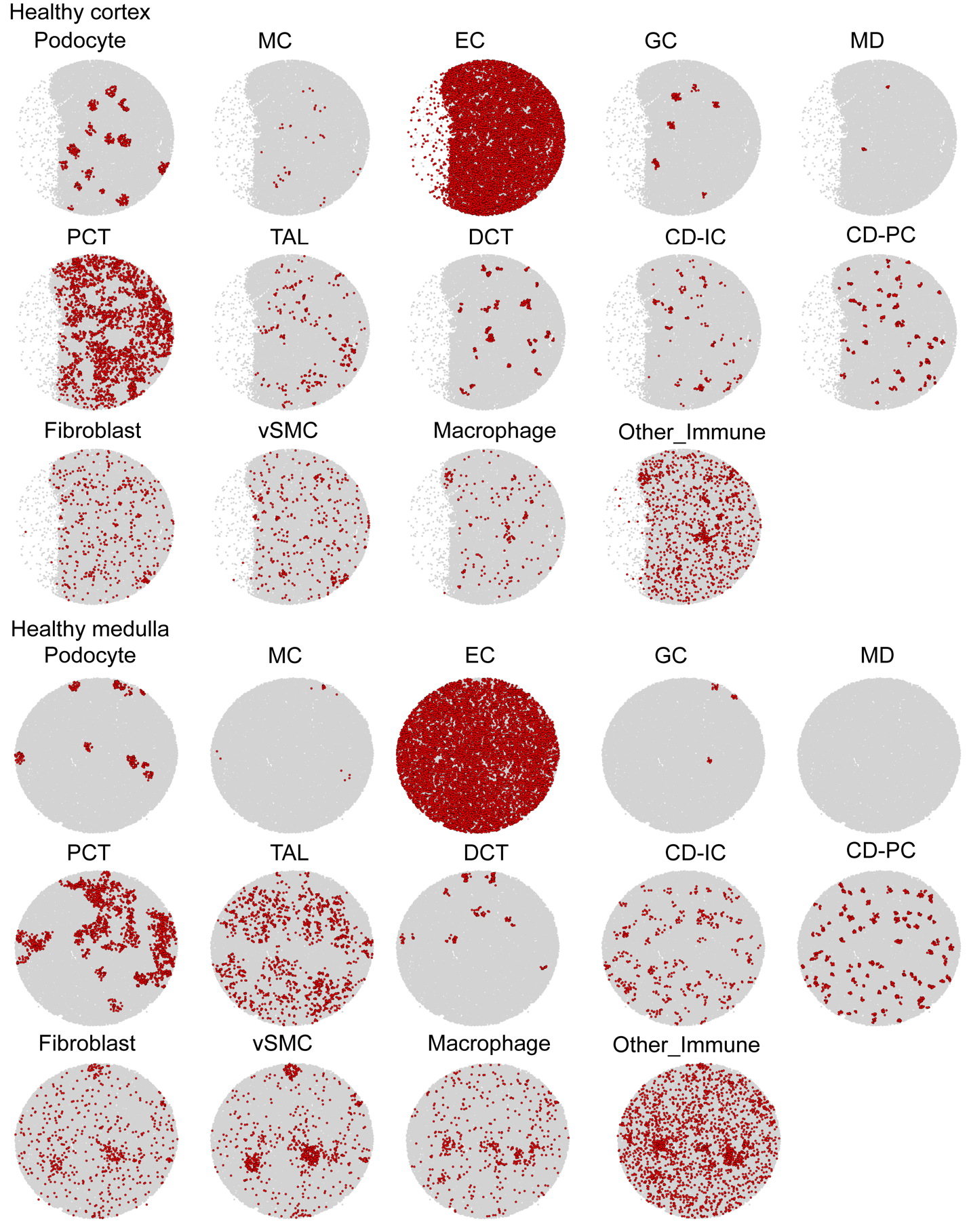

**Fig. S61. Individual cell type assignments in healthy human cortex and medulla.** Assignment of all curated cell types are shown for healthy human cortex (top) and medulla (bottom).

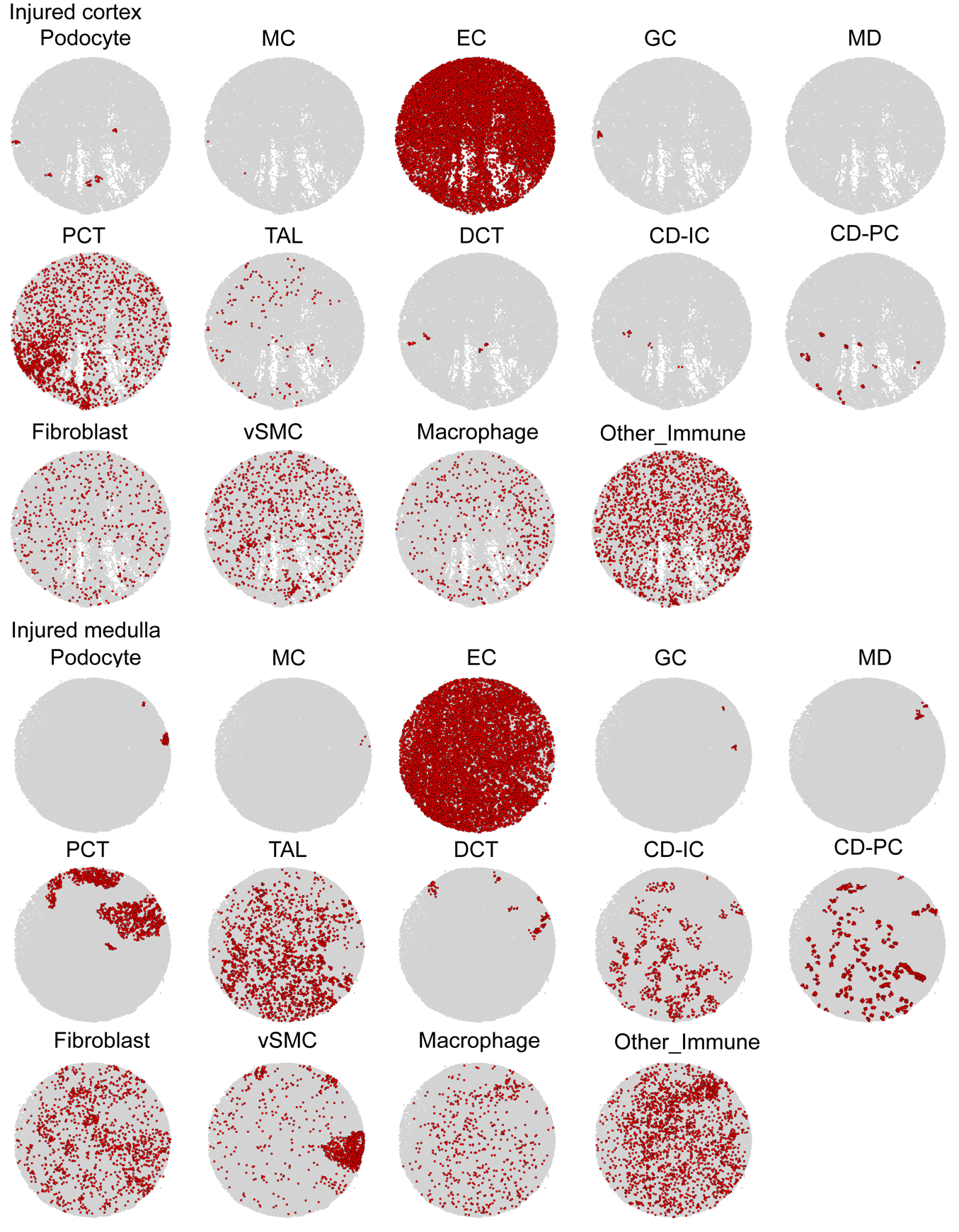

**Fig. S62. Individual cell type assignments in injured human cortex and medulla.** Assignment of all curated cell types are shown for injured human cortex (top) and medulla (bottom).

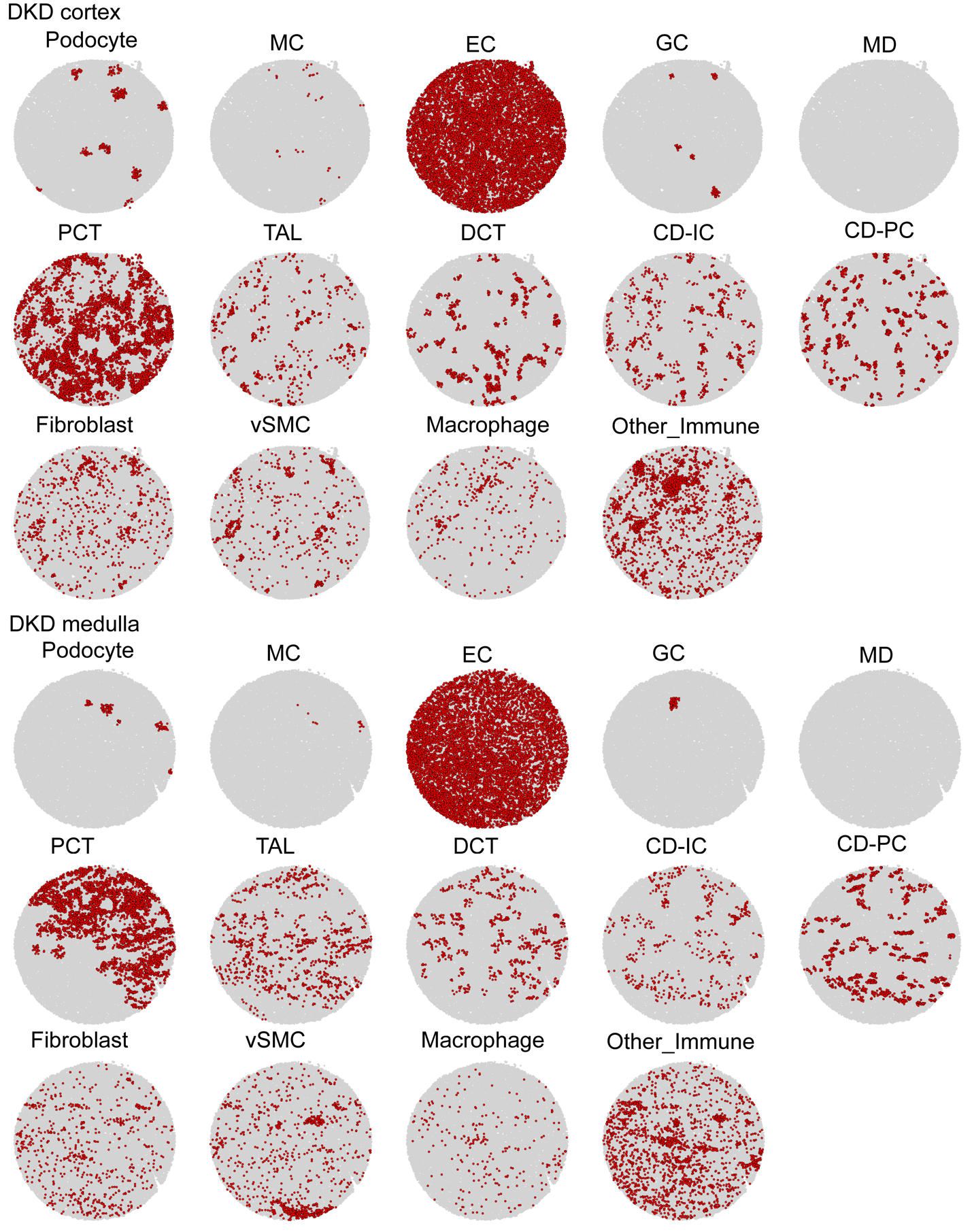

**Fig. S63. Individual cell type assignments in DKD human cortex and medulla.** Assignment of all curated cell types are shown for DKD human cortex (top) and medulla (bottom).

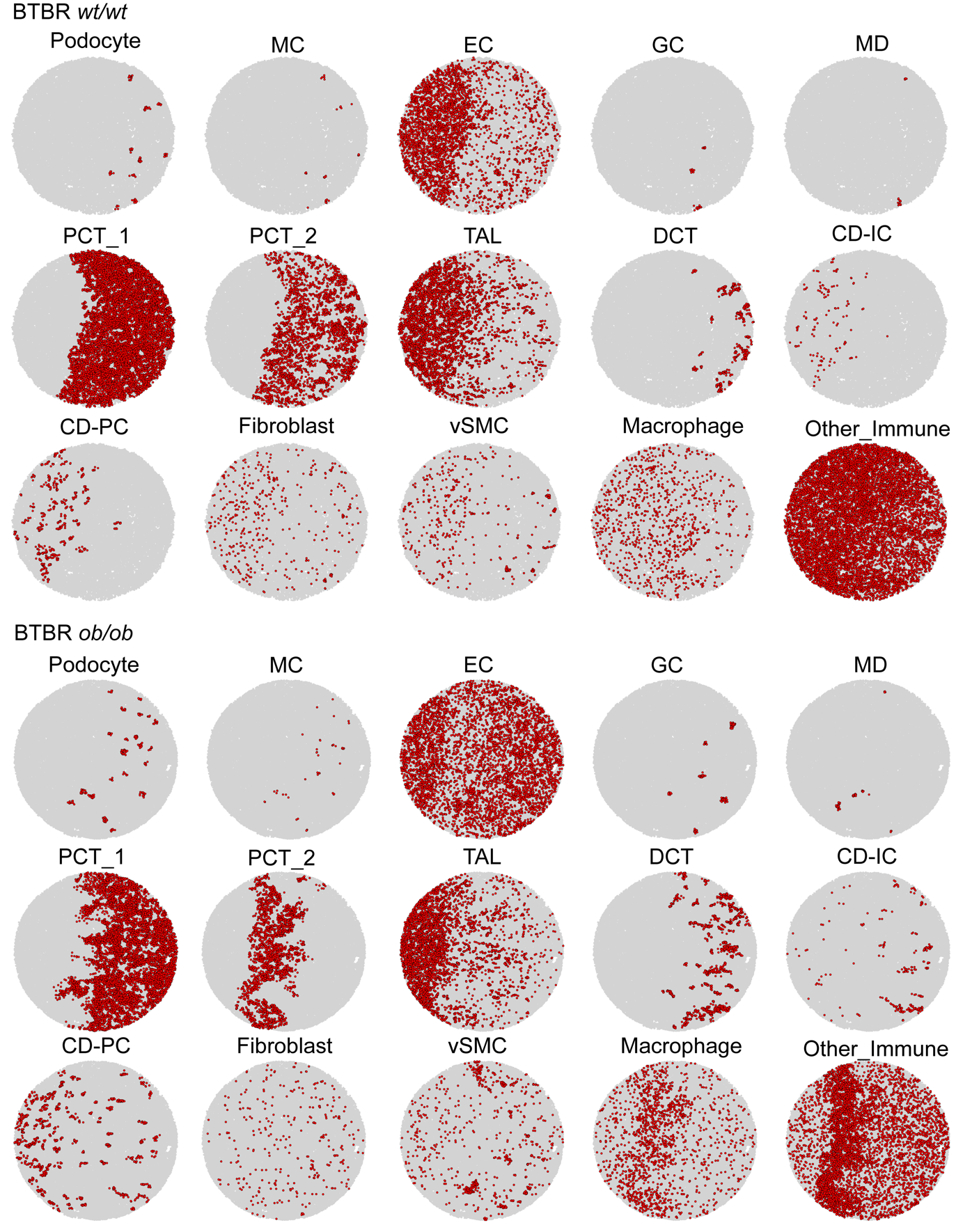

**Fig. S64. Individual cell type assignments in BTBR *wt/wt* and BTBR *ob/ob* kidney tissue.** Assignment of all curated cell types are shown for BTBR *wt/wt* (top) and BTBR *ob/ob* (bottom) kidney tissue.

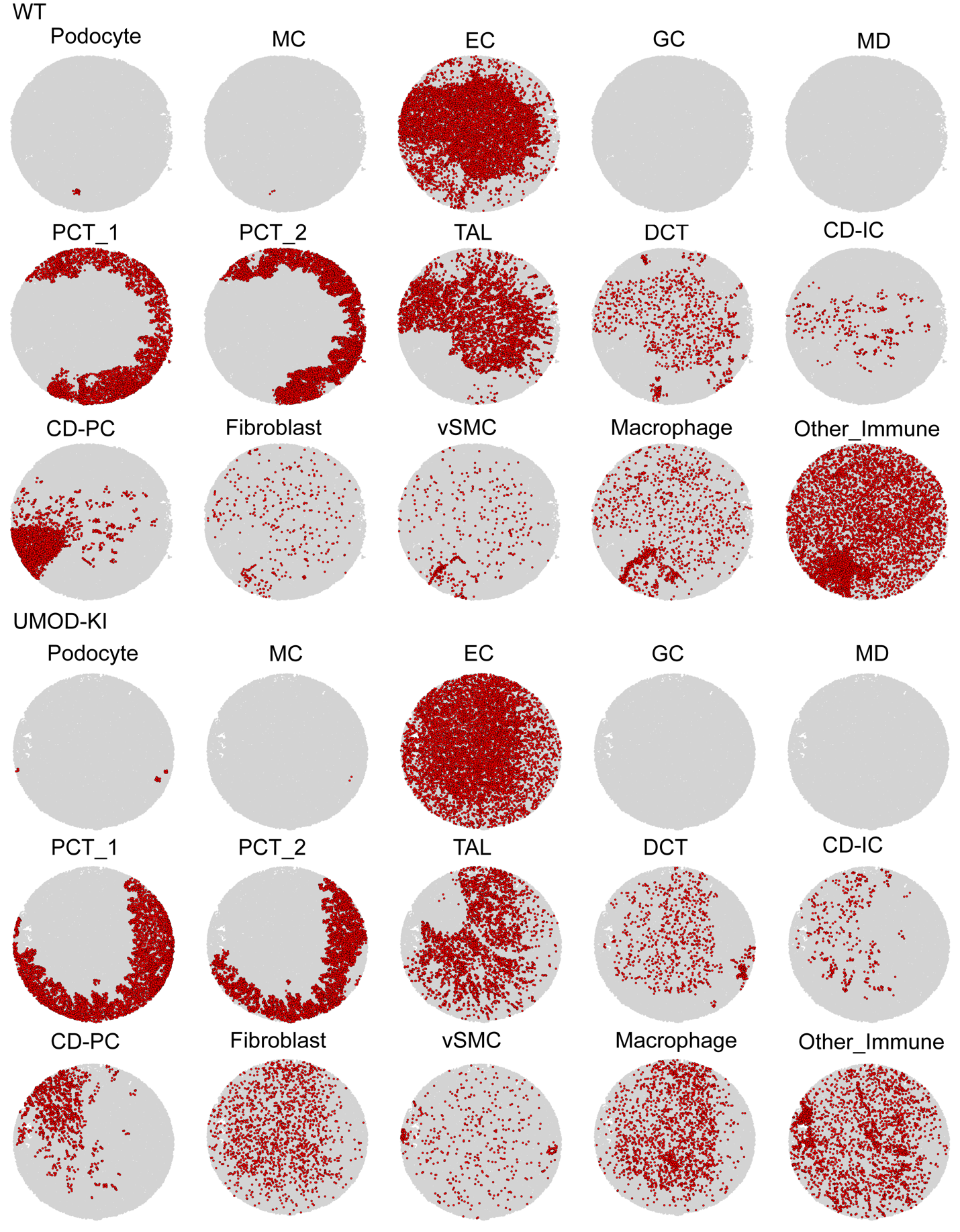

**Fig. S65. Individual cell type assignments in WT and UMOD-KI kidney tissue.** Assignment of all curated cell types are shown for WT (top) and UMOD-KI (bottom) kidney tissue.

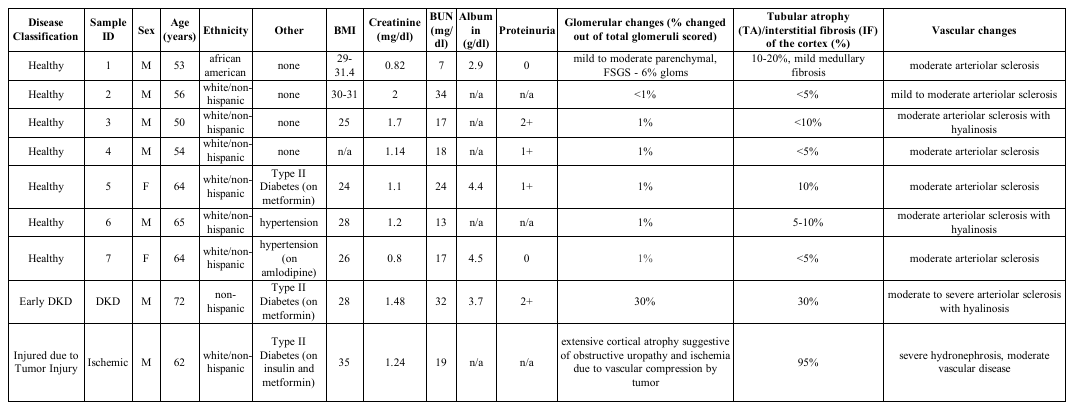

**Table. S1. Human tissue samples.**

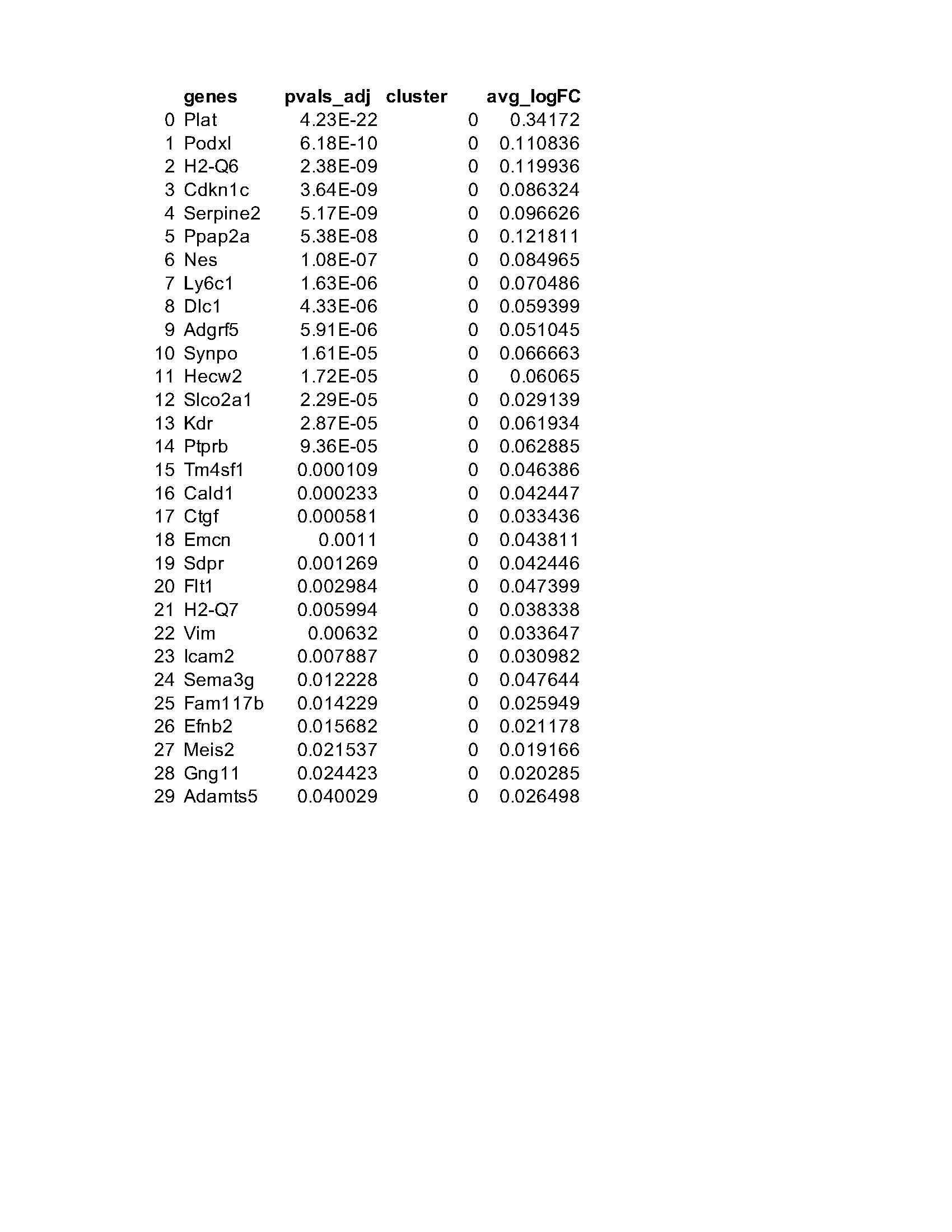

**Table S2: BTBR *ob/ob* glomerular EC DE disease genes.**

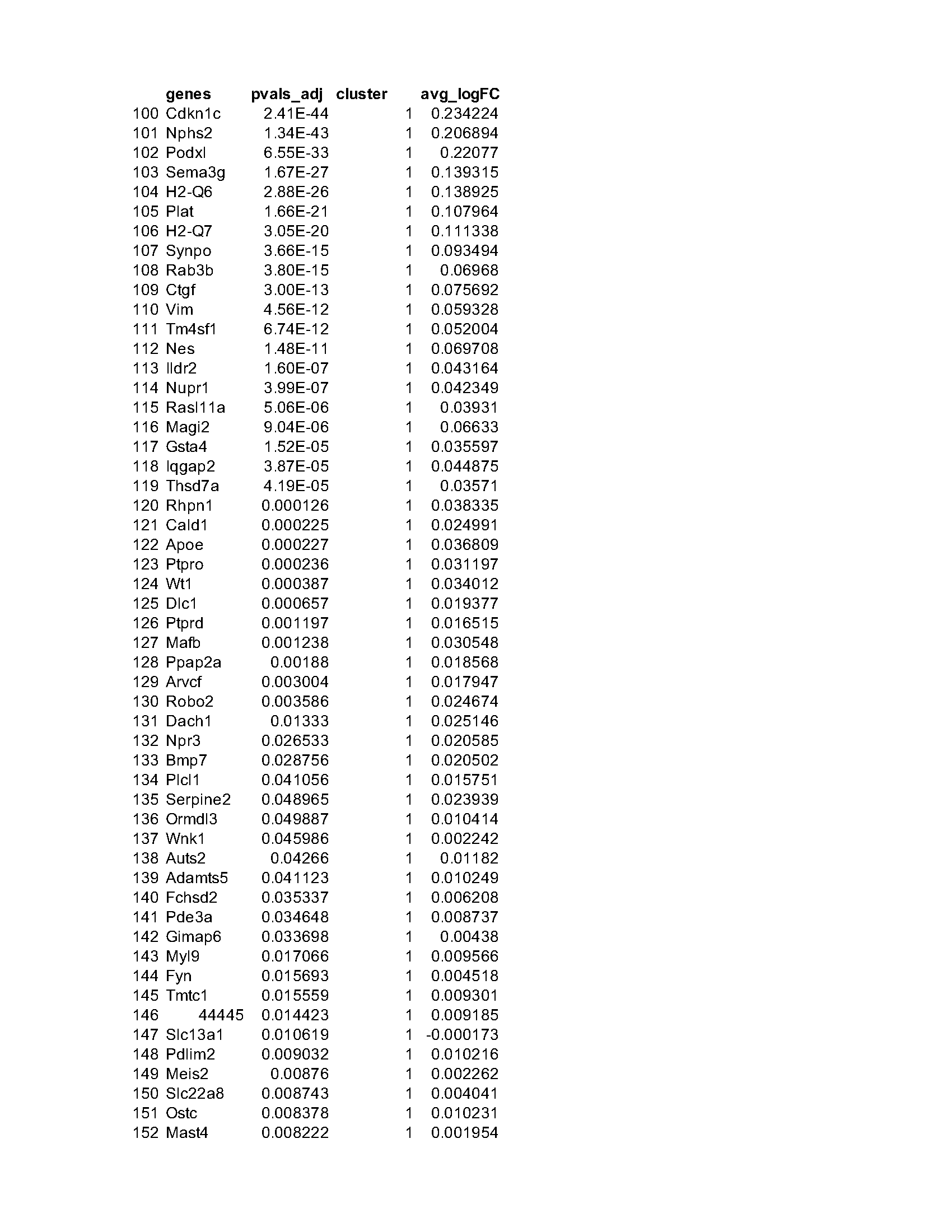

**Table S3: BTBR *ob/ob* glomerular podocyte DE disease genes.**

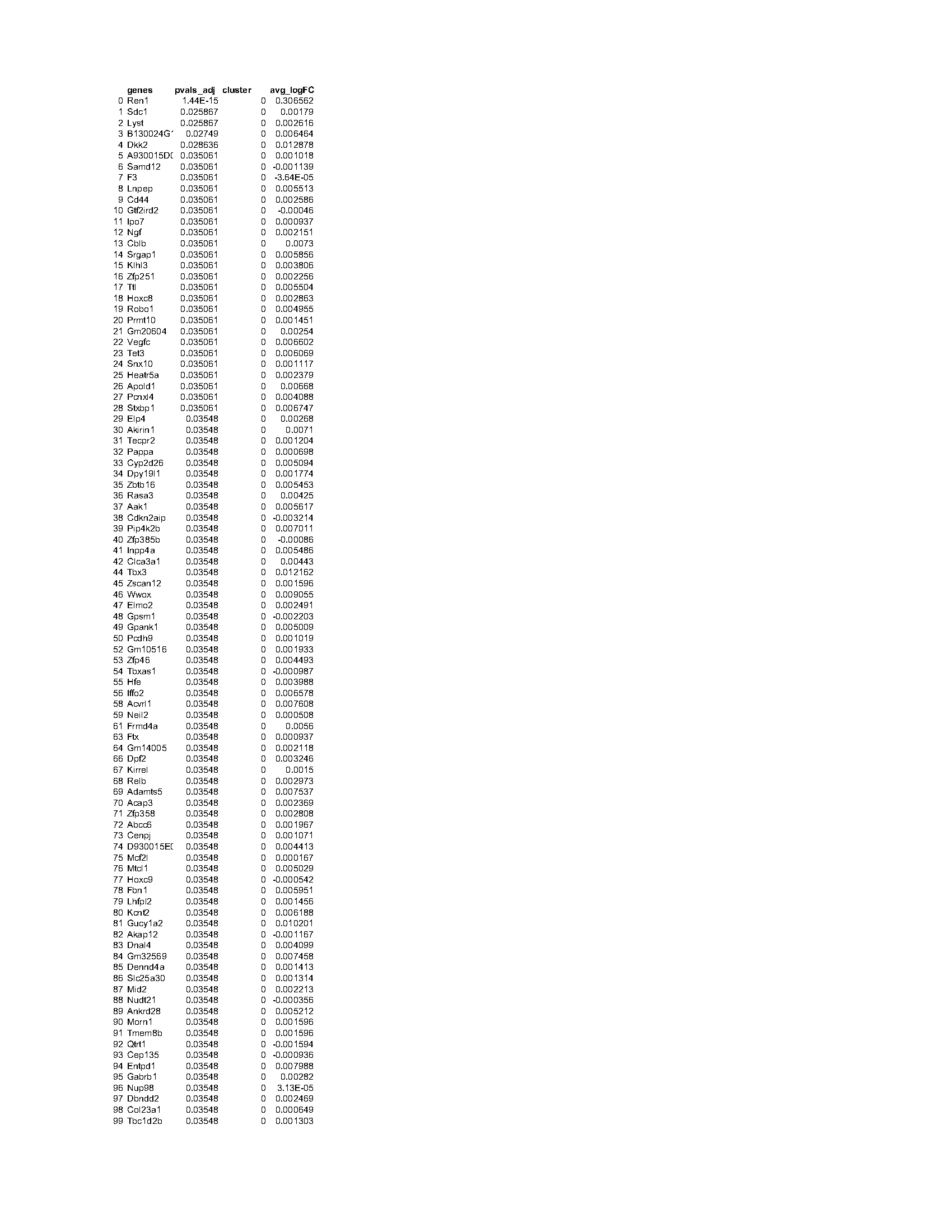

**Table S4: BTBR *ob/ob* GC DE disease genes.**

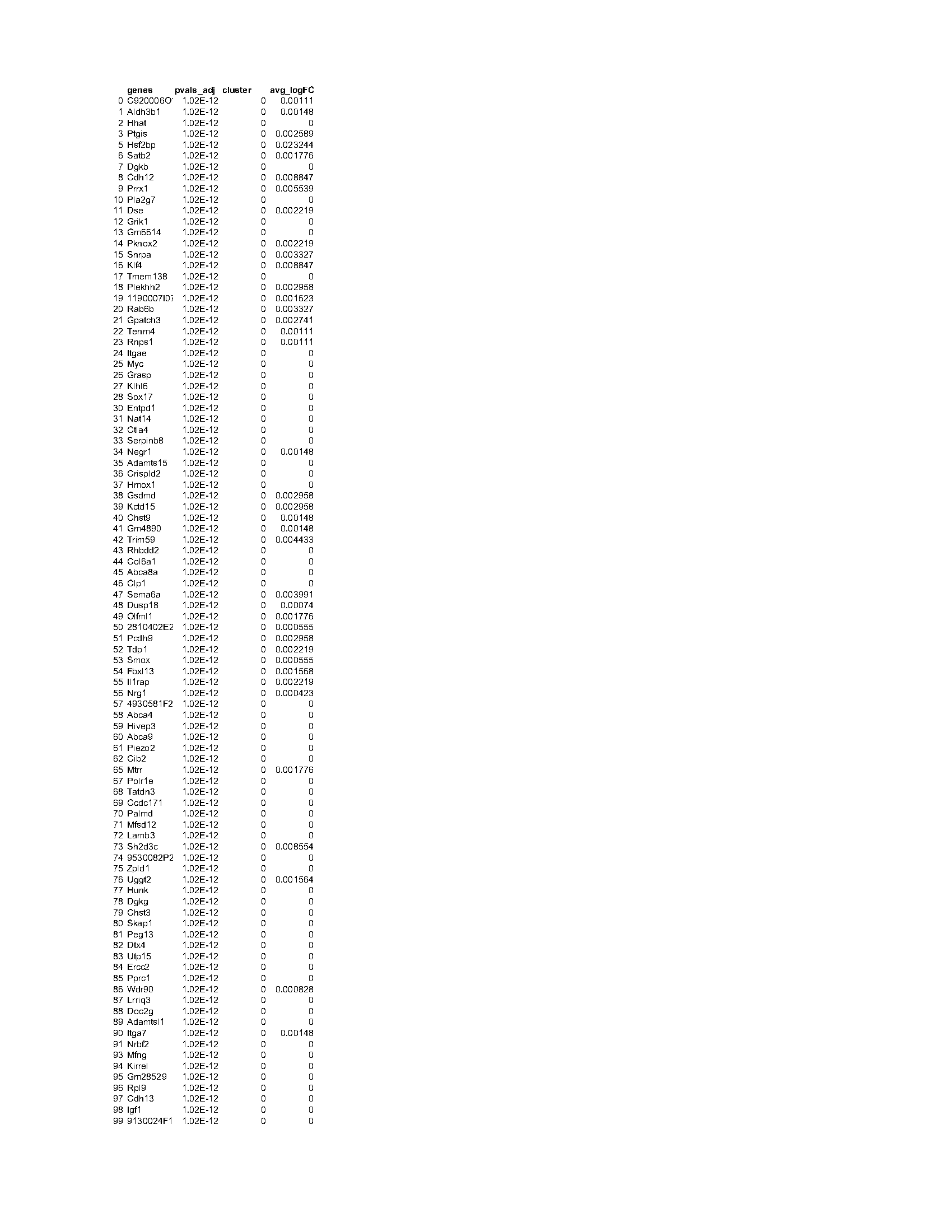

**Table S5: BTBR *ob/ob* CD-IC cortex DE disease genes.**

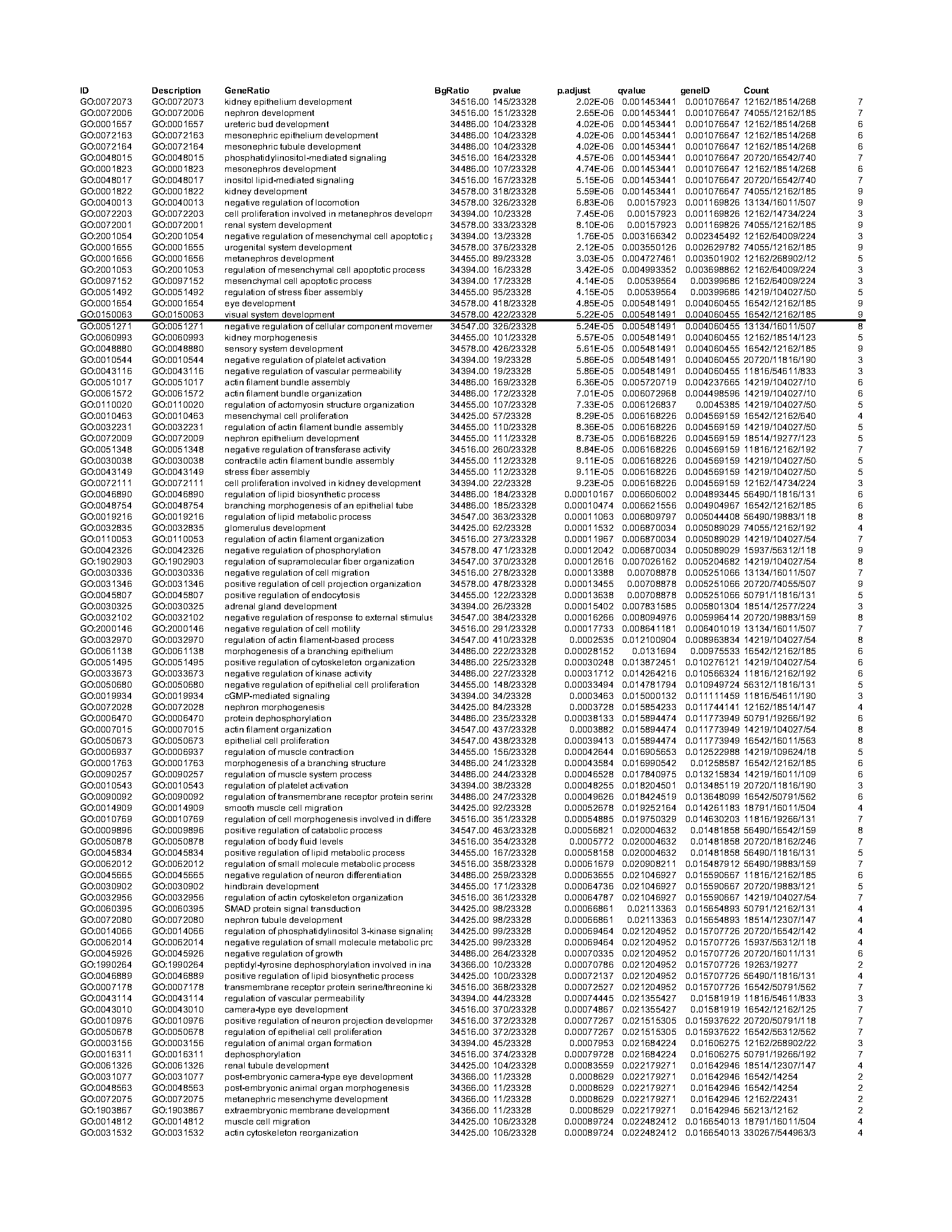

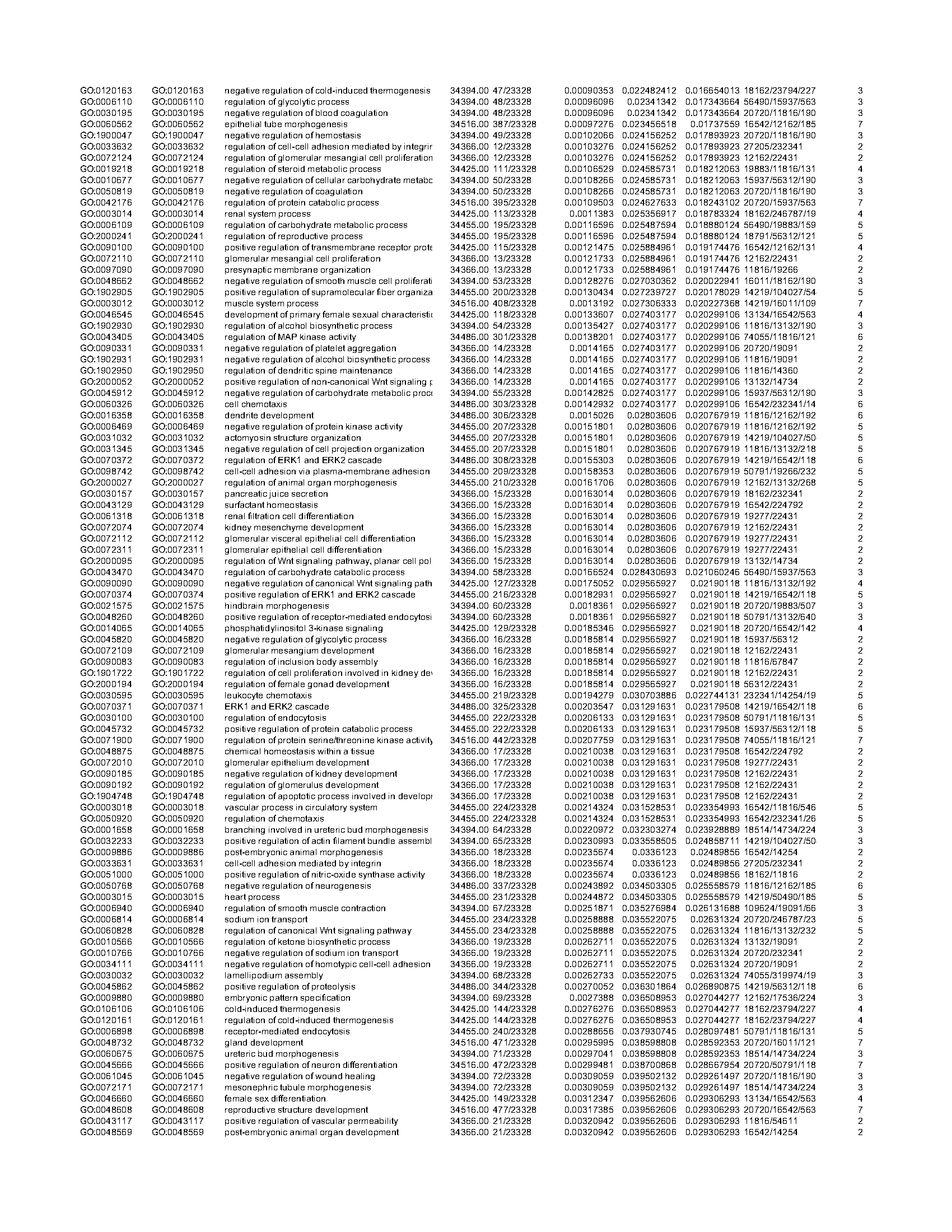

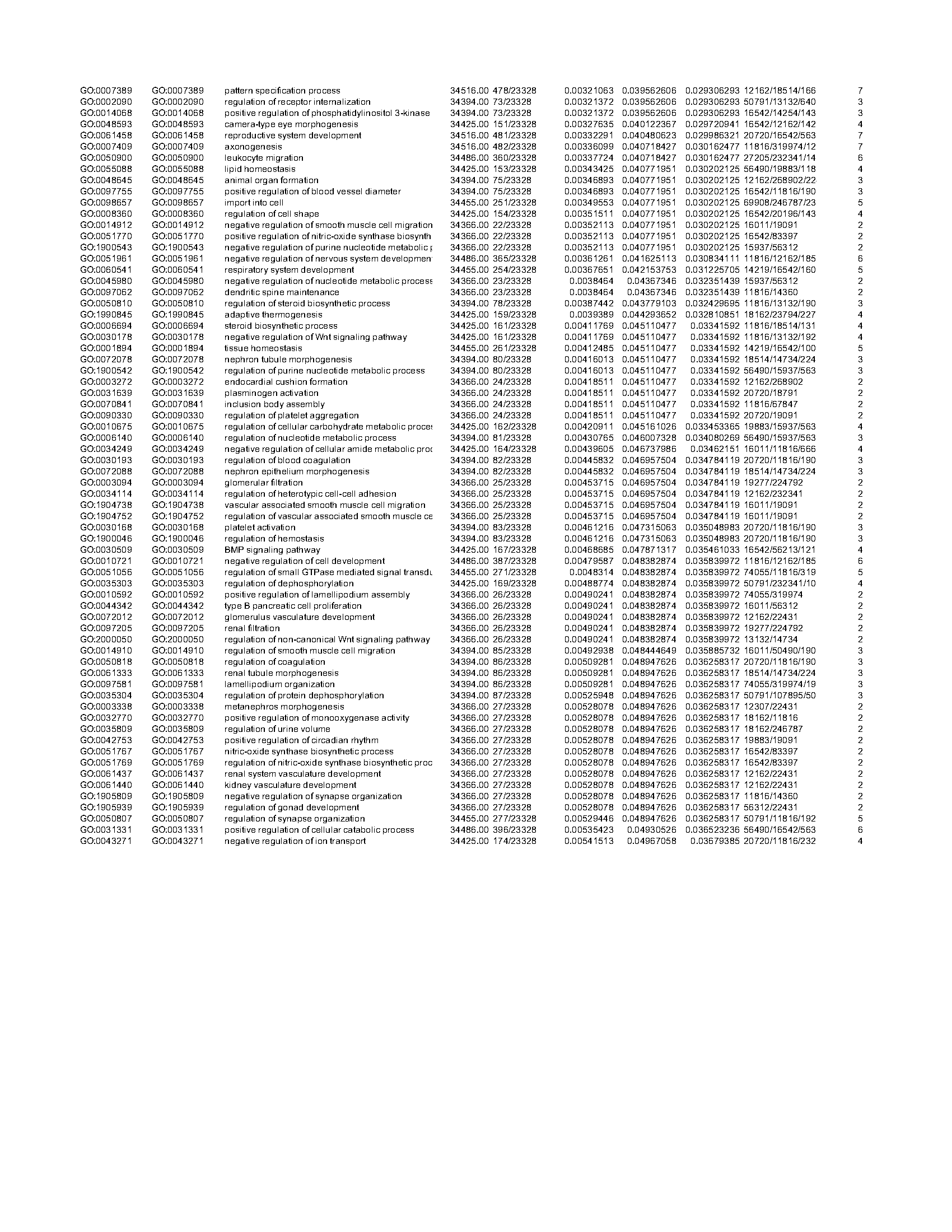

**Table S6: BTBR *ob/ob* GO analysis of glomerular podocyte DE disease genes.**

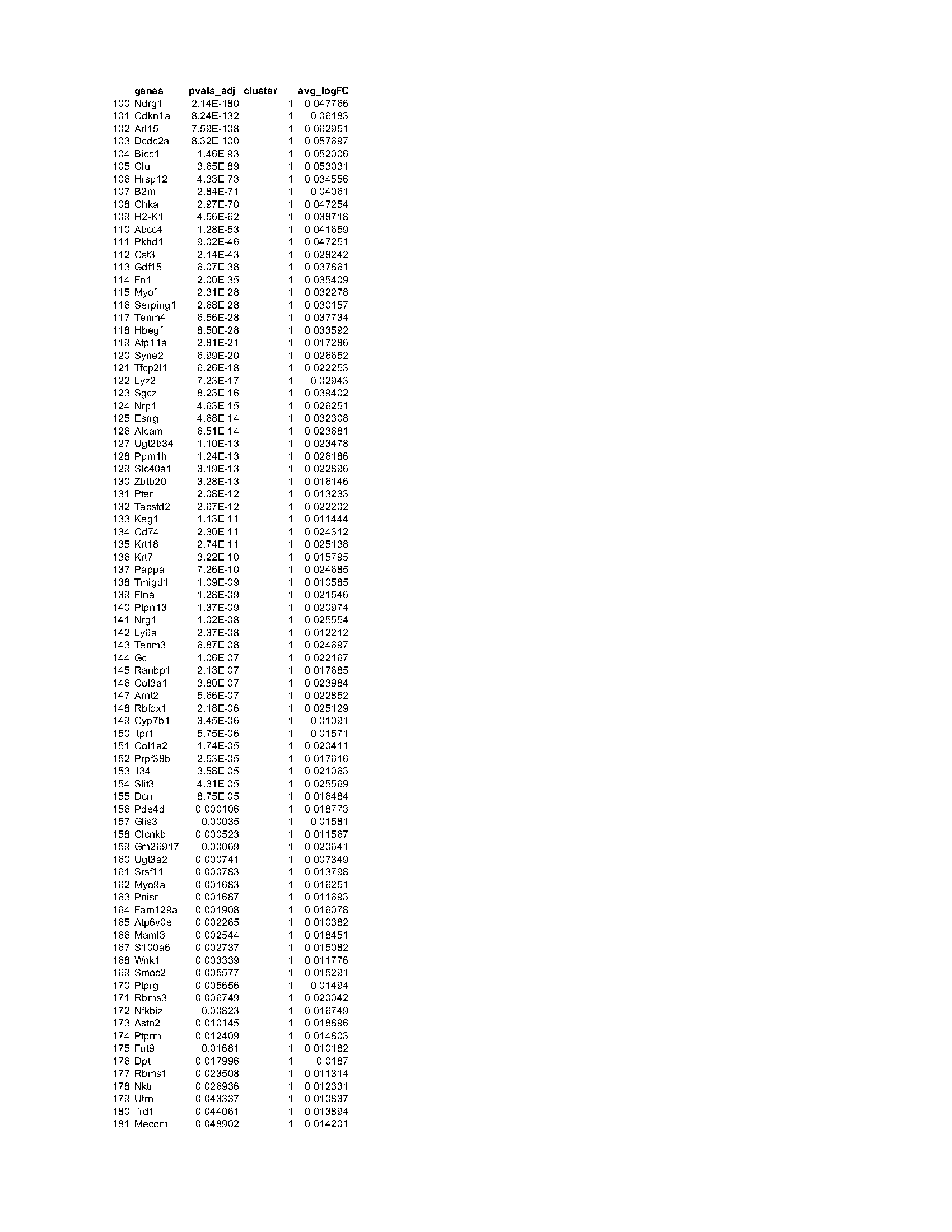

**Table S7: UMOD-KI TAL medulla DE disease genes.**

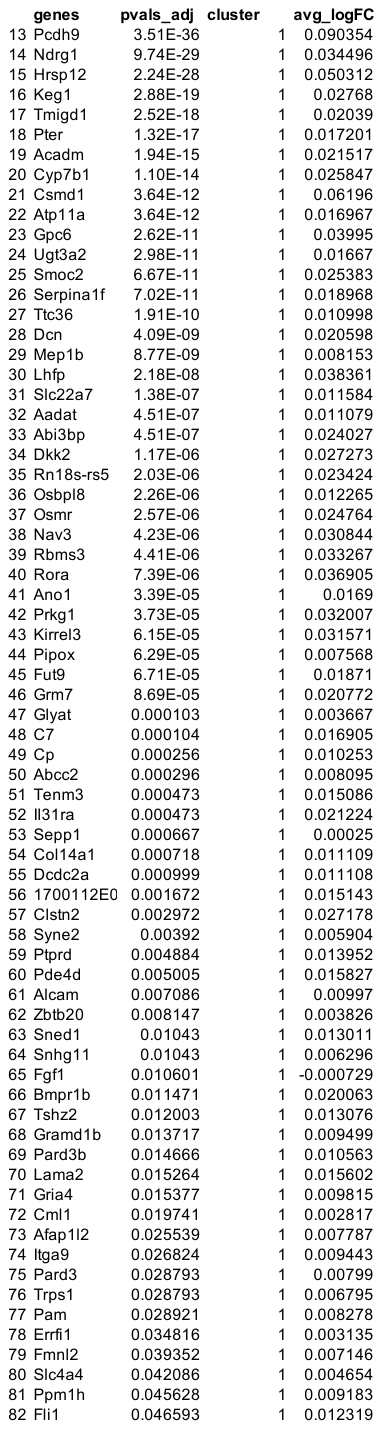

**Table S8: UMOD-KI fibroblast medulla DE disease genes.**

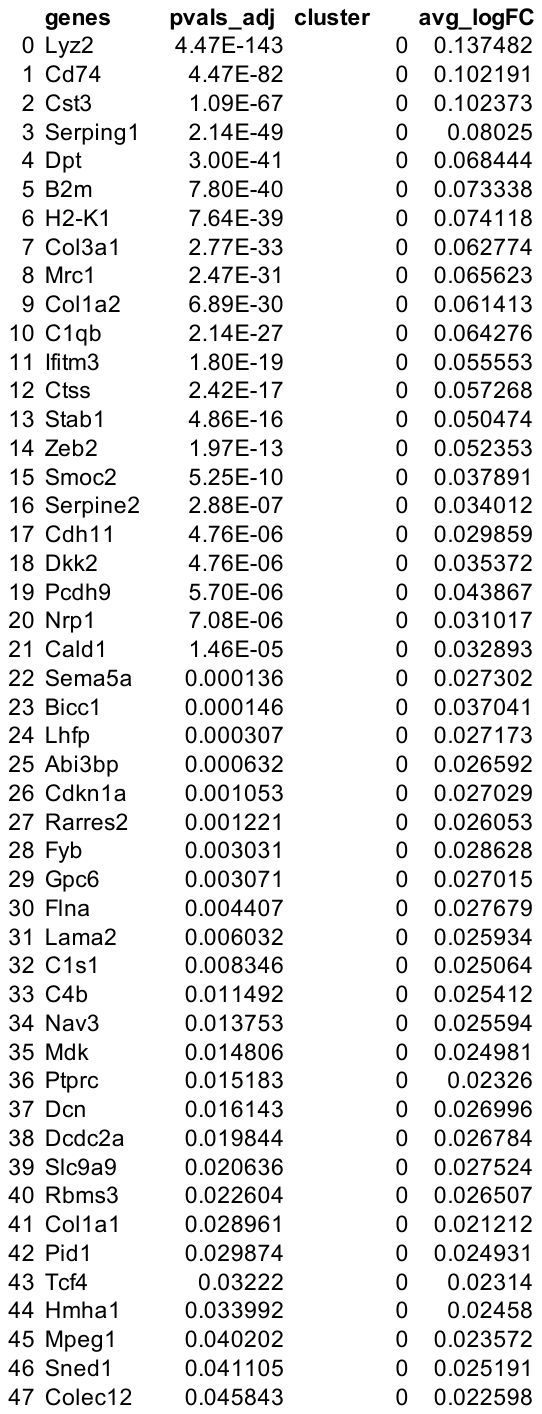

**Table S9: UMOD-KI macrophage medulla DE disease genes.**

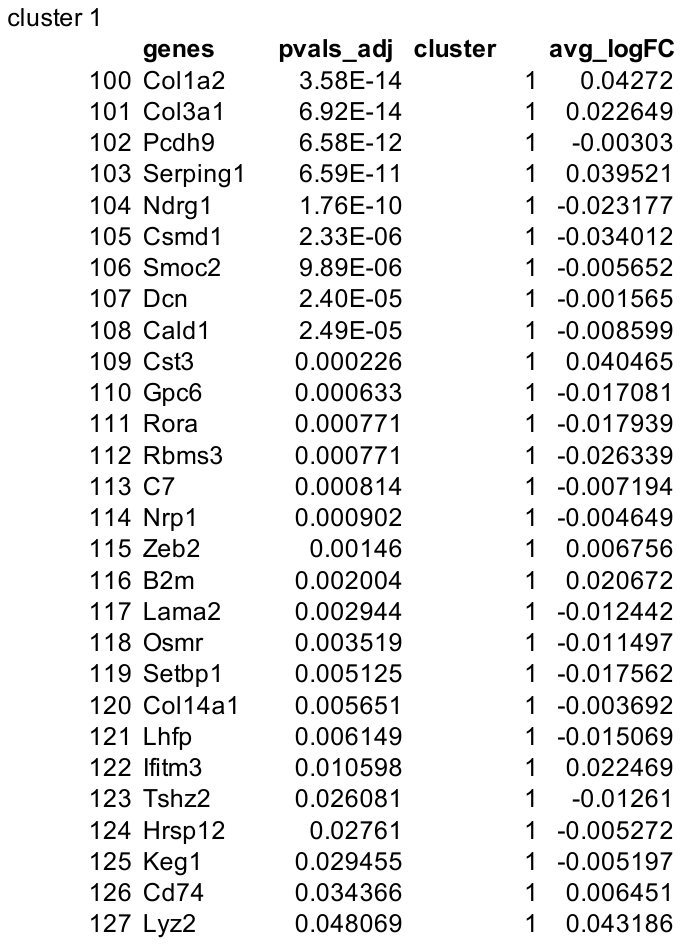

**Table S10: UMOD-KI fibroblast cortex DE disease genes.**

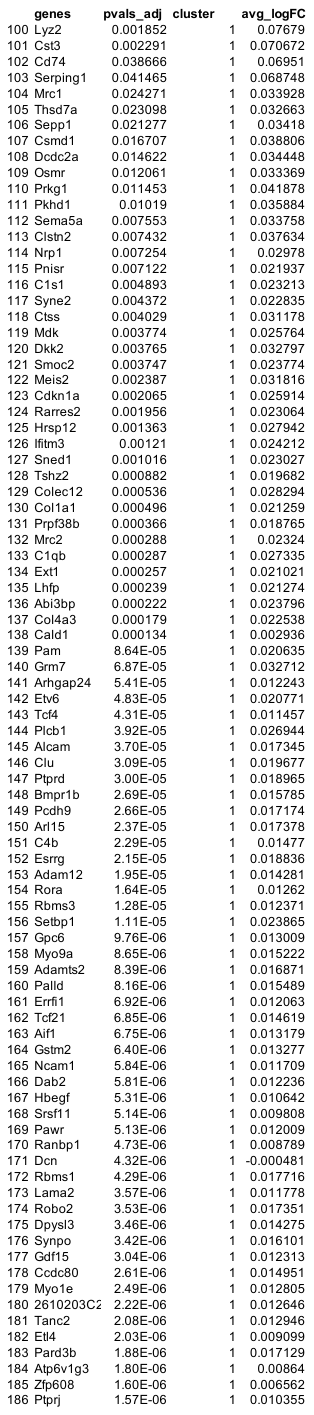

**Table S11: UMOD-KI vSMC medulla DE disease genes.**

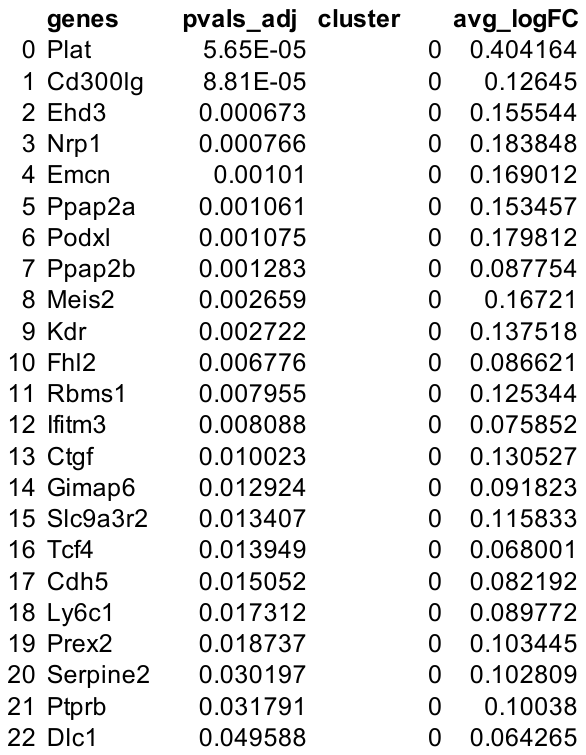

**Table S12: UMOD-KI glomerular EC DE disease genes.**

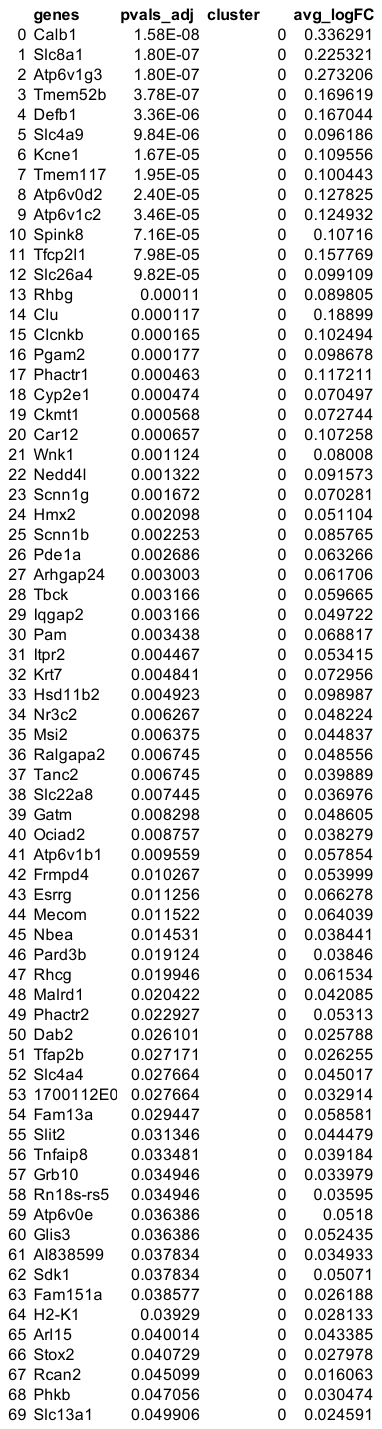

**Table S13: UMOD-KI CD-PC cortex DE disease genes.**

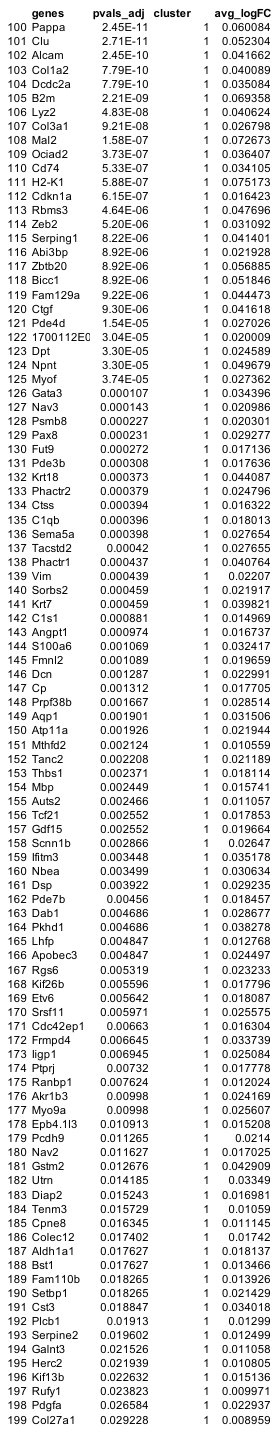

**Table S14: UMOD-KI CD-PC medulla DE disease genes.**
